# Ancestry-related differences in chromatin accessibility and gene expression of *APOE4* are associated with Alzheimer disease risk

**DOI:** 10.1101/2022.10.27.514114

**Authors:** Katrina Celis, Maria DM. Muniz Moreno, Farid Rajabli, Patrice Whitehead, Kara Hamilton-Nelson, Derek M. Dykxhoorn, Karen Nuytemans, Liyong Wang, Clifton L. Dalgard, Margaret Flanagan, Sandra Weintraub, Changiz Geula, Marla Gearing, David A. Bennett, Theresa Schuck, Fulai Jin, Margaret A. Pericak-Vance, Anthony J. Griswold, Juan I. Young, Jeffery M. Vance

## Abstract

**Background:** European local ancestry (ELA) surrounding *APOE4* is associated with a higher risk for Alzheimer Disease (AD) compared to African local ancestry (ALA). We previously demonstrated significantly higher *APOE4* expression in ELA vs ALA in the frontal cortex of *APOE4/4* AD patients. Differences in chromatin accessibility could contribute to these differences in *APOE4* expression.

**Methods:** We performed single nuclei Assays for Transposase Accessible Chromatin sequencing (snATAC-seq) and single nuclei RNA sequencing (snRNA-seq) from frozen frontal cortex of six ALA and six ELA AD patients, all homozygous for local ancestry and *APOE4*.

**Results:** We demonstrated that *APOE4*, including its promoter area, has greater chromatin accessibility in ELA vs ALA astrocytes. This increased accessibility in ELA astrocytes extended genome wide. Genes with increased accessibility and expression in ELA in astrocytes were enriched for synaptic function, cholesterol processing and astrocyte reactivity.

**Conclusion:** Our results suggest that increased chromatin accessibility of *APOE4* in astrocyte with the ELA contributes to the observed elevated *APOE4* expression, corresponding to the increased AD risk in ELA vs ALA *APOE4/4* carriers.

## BACKGROUND

Alzheimer disease (AD) is the most common neurodegenerative disorder and one of the leading causes of disability worldwide for individuals aged 75 and older^1^. The strongest genetic risk factor to develop late-onset AD is the apolipoprotein E4 (*APOE4*) allele located on chromosome 19^2,3^. However, the risk associated with *APOE4* differs dramatically between individuals of European vs African ancestries^4–7^ with *APOE4* carriers of European ancestry having substantially increased risk for AD compared to individuals with African ancestry.

This observed difference in risk has recently been shown in three independent studies to be due to differences in the genetic local ancestry (LA) region surrounding *APOE* rather than overall global European or African ancestry^8–10^. Using single nucleus RNA sequencing (snRNA-seq) of frontal cortex we have previously demonstrated that *APOE4/4* homozygous carriers with European local ancestry (ELA) expressed significantly higher *APOE* compared to those with African local ancestry (ALA), especially in astrocytes^11^, suggesting a potential mechanism underlying the differential AD risk seen between ancestries.

Control of gene expression is orchestrated by the integration of cis-regulatory modules, such as enhancer and promoter elements, along with transcription factors^12–14^. The cis-regulatory modules of actively transcribed genes are generally in ‘accessible’ euchromatin with low nucleosome occupancy and few high-order structures^15–17^, while ‘closed’ chromatin (heterochromatin) generally associates with transcriptional silencing^16,18,19^. Accordingly, ancestry-specific changes in accessibility of cis-regulatory elements that modulate the binding of transcription factors is a potential mechanism responsible for the difference in *APOE4* expression seen between ancestries.

The single nuclei Assay for Transposase Accessible Chromatin followed by sequencing (snATAC-seq), facilitates the creation of chromatin accessibility maps with single cellular resolution. For example, Corces et al^20^ demonstrated that open chromatin regions in promoter or enhancer areas are associated with increased gene expression through snATAC-seq. Investigation of chromatin accessibility profiles specifically in AD have mostly focused on pathological hallmarks like accumulation of Tau and amyloid-β^21^ or altered APP signaling pathways^22^. More recently, Morabito et al^23^ conducted an integrated analysis using snATAC-seq and snRNA-seq from brains of AD and cognitively normal individuals and reported that cell-type-specific chromatin accessibility changes in regulatory elements may be key to regulate gene expression changes in AD. However, to date, the role of chromatin accessibility in AD between different ancestries, specifically at the *APOE4* locus, has not been investigated.

Therefore, we performed snATAC-seq combined with snRNA-seq to investigate the cell specific patterns of chromatin accessibility in ELA and ALA *APOE4/4* brains. We observed increased accessibility in *APOE* ELA relative to ALA in astrocytes. Increased chromatin accessibility in ELA samples was also observed genome-wide in this cell type. The differentially accessible peaks at the *APOE* promoter are predicted to bind a subset of transcription factors that exhibit differential gene expression in ELA astrocyte samples. These data support the hypothesis that European ancestry at the *APOE* promoter and in the area around *APOE* have a more open chromatin conformation than African ancestry. We speculate that this difference is permissive for transcription factor binding and transcriptional activation of *APOE*. Thus, these findings provide novel insights into the molecular mechanisms of ancestry specific differences in AD risk.

## METHODS

### Brain Samples

Brain autopsy samples were obtained as part of a multi-center collaboration from four Alzheimer Disease Research Centers (ADRCs; Emory University, Northwestern University, Rush University Medical Center, and the University of Pennsylvania) and from the John P. Hussman Institute for Human Genomics (HIHG) at the University of Miami. All ADRC samples were initially selected from the National Alzheimer Coordinating Center (NACC) following filtering criteria previously described^11^. All selected samples were compliant with site-specific approved institutional review board protocols.

Genotyping arrays were processed for all samples to assess both global and local genetic ancestry (LA) in the *APOE* region, defined as within 1Mb on either side of *APOE* (chr19:44–46Mb)^8^. Ancestry analyses were performed as previously described^11^. Individuals homozygous for the ELA or ALA haplotype in this region were selected for Whole Genome Sequencing (WGS) at either the Center for Genome Technology (CGT) at the HIHG or The American Genome Center at Uniformed Services University of the Health Sciences (USUHS) using the TruSeq DNA PCR-Free library preparation kit and sequencing on the Novaseq 6000 (Illumina, San Diego, CA). WGS was analyzed through a bioinformatics pipeline including alignment to the GRCh38 genome build with BWA mem, duplicate marking and base-quality recalibration, and genotype calling with the HaplotypeCaller as recommended by the Genome Analysis Toolkit best practices.

### Nuclei Isolation for snATAC-seq

Nuclei were isolated from frozen frontal cortex brain tissue (Brodmann area 9) as described previously^24,25^. Briefly, ~100mg of frozen brain tissue was dissociated by Dounce homogenization to create a nuclear suspension. Nuclei were purified using an iodixanol gradient and washed in resuspension buffer (RSB). Nuclei were counted and, for each sample, 65,000 nuclei were aliquoted into a separate tube containing RSB with 0.1% Tween-20. Nuclei were assessed for quality using the Nexcelom Cellometer K2 and a propidium iodide fluorescent dye.

### snATAC-seq data generation and analysis

SnATAC-seq was performed at the HIHG CGT. A suspension of 65,000 nuclei was prepped according to the 10X Genomics Chromium Next GEM Single Cell ATAC 1.1 protocol. In short, single nuclei were isolated on a 10x Chromium Next GEM Chip H and library construction was completed by incorporation of Illumina adapters via PCR. Libraries were sequenced on an Illumina NovaSeq 6000 sequencer targeting 75,000 reads per nuclei in paired end 50bp sequencing reactions. Resulting FASTQ files were processed using the CellRanger ATAC software package v1.2 (10X Genomics) including cell identification, alignment to the GRCh38 human reference genome, and generation of fragment files for downstream analysis. snRNA-seq was performed as previously described^11^.

Integrated analysis of the snATAC-seq and snRNA-seq data was performed using the ArchR software package v1.0.1^26^. First, fragment files from CellRanger were converted to ARROW files to include cells with a minimum of 1000 fragments and a transcription start site (TSS) enrichment score of at least two. Doublet nuclei inference and removal were performed as described^26^. Clustering of cells was performed with five iterations of Iterative Latent Semantic Indexing (LSI)^27,28^ followed by batch correction with Harmony^29^. We then identified ATAC clusters using the FindClusters function from the Seurat v4.0 software package^30^ and refined the cell type definitions by integration with snRNA-seq from the same samples using the Seurat FindTransferAnchors function.

### SnATAC-seq peak calling and differential accessibility

To identify ancestry specific peaks within each cell cluster, we created pseudo-bulk replicates combining all ELA and all ALA samples separately and called peaks for each ancestry within each cluster using the addReproduciblePeakSet function in ArchR utilizing the default parameters of the MACS2 callpeak command v2.2.7.1^31^. Differential accessibility for each peak between ELA and ALA within each cluster was calculated using the ArchR getMarkerFeatures function.

### ATAC peak annotations

Peak annotation was performed with HOMER^32^ and GREAT^33^. HOMER (version 4.11^34^, was used to annotate peaks closest to transcription start sites (TSS) and to indicate the genomic region type of the peak (exon, intron, intergenic, or promoter-TSS). A peak was assigned to a gene promoter-TSS when the peak location was ±2kb from the TSS of the associated gene. GREAT (version 4.0.4, http://great.stanford.edu/public/html/) was used to refine peak annotations where the peak was near multiple genes or was intergenic. Distal enhancers were identified with the ELITE GeneHancer database from UCSC^35^ of enhancers and promoters and their inferred target genes supported by more than one source of evidence. We considered as distal enhancer peaks those annotated by GREAT or HOMER farther than ±2kb from a TSS and with coordinates overlapping one of these ELITE enhancers. All other peaks were considered as intergenic.

### Transcription Factor Analysis

Bedtools getfasta^36^ was used to extract the sequence of each peak from the reference human genome (refdata-cellranger-atac-GRCh38-1.2.0 version). The algorithm FIMO^37^ within MEME Suite version 5.4.1^38^ was used to identify known transcription factor (TF) binding motifs in the two *APOE* promoter differentially accessible peaks (DAP). We surveyed those peak sequences against experimentally defined transcription factor binding site databases JASPAR CORE 2022 (vertebrates non redundant)^39^, Jolma 2013 (human and mouse)^40^, and Swiss regulon (human and mouse)^41^, and selected those with false discovery rate (FDR) adjusted *p*-value <0.05. In addition, we used all the human specific JASPAR 2022 defined TFs of the UCSC HG38 table browser^42^ binding at those peak coordinates filtering by a JASPAR score ≥ 300.

### Functional Enrichment Analysis

Functional analysis of differentially expressed genes (DEG), differentially accessible genes (DAG), and both differentially expressed and accessible genes (DEG-DAG) was performed using enrichR^43^. Enrichment for cell type specific markers was performed using the cell type gene-set atlas lists from Azimuth 2021^44^ and PanglaoDB 2021^45^. Functional pathway enrichment was performed using as reference the human specific gene-set libraries KEGG 2021^46^, the Gene Ontology Biological Process 2021^47^, Reactome 2021^48^ and ENCODE Histone post-translational modifications^49^. Disease gene enrichment analysis used the gene-sets extracted from GeneSigDb^50^, HDSigDb^51^ and MsigDB^52^. The enrichment in chromosomal location was performed using the annotations of DEGs-DAGs per chromosome and using a Fischer exact test.

## RESULTS

### Brain Sample Characteristics

In total, twelve brain samples were used for the study (Table 1). Six ELA samples (European global ancestry >96%) were obtained from the HIHG Brain Bank (3), Rush University Medical Center (2) and Emory University (1). Six ALA samples (African global ancestry 68% to 88%) were obtained from Emory University (2), Northwestern University (1) and Rush University Medical Center (3). The samples included seven females (four ALA and three ELA) and five males (two ALA and three ELA). All samples had BRAAK staging scores ranging from IV to VI and had a mean age-of-death of 79 years. Whole genome sequencing confirmed the homozygous *APOE4/4* genotype and the absence of mutations in known Mendelian genes for AD (*PSEN1, PSEN2, APP, MAPT*) as well as the absence of known rare AD risk variants in *ABCA7, TREM2* and *SORL1*.

**Table 1.** Demographic characteristics of the samples

| Sample | Center | Sex | AOD | *APOE* genotype | Local Ancestry (Chr19: 44-46Mb) | % Of Global Ancestry | BRAAK staging score |
| --- | --- | --- | --- | --- | --- | --- | --- |
| 1 | Emory | Female | 82 | 4,4 | AF/AF | 87% AF | V |
| 2 | NW | Female | 85 | 4,4 | AF/AF | 86% AF | V |
| 3 | Emory | Female | 80 | 4,4 | AF/AF | 88% AF | VI |
| 4 | Rush | Male | 93 | 4,4 | AF/AF | 68% AF | V |
| 5 | Rush | Male | 83 | 4,4 | AF/AF | 79% AF | VI |
| 6 | Rush | Female | 77 | 4,4 | AF/AF | 85% AF | VI |
| 7 | UM/Duke | Male | 75 | 4,4 | EU/EU | 95% EU | IV |
| 8 | Rush | Male | 71 | 4,4 | EU/EU | 98% EU | VI |
| 9 | UM/Duke | Female | 70 | 4,4 | EU/EU | 95% EU | VI |
| 10 | Rush | Male | 76 | 4,4 | EU/EU | 97% EU | IV |
| 11 | UM/Duke | Female | 76 | 4,4 | EU/EU | 96% EU | V |
| 12 | Rush | Female | 86 | 4,4 | EU/EU | 98% EU | VI |

AOD: Age of Death; UM: University of Miami; NW: Northwestern University; RUSH: RUSH University Medical Center; AF: African; EU: European

### Cell type identification

We obtained data from a total of 60,306 nuclei (with a mean of 4,200 nuclei per sample) from snATAC-seq and 94,411 nuclei (with a mean of 7,868 nuclei per sample) from snRNA-seq. The snATAC-seq libraries had an average of ~13,000 fragments per nucleus and the snRNA-seq libraries had a median depth of ~116,000 reads per nucleus with on average ~1,900 genes/nucleus (Supplementary Table 1). The clusters with more than 500 nuclei in ELA and ALA showed no statistical differences in the number if nuclei per cluster between the ancestry groups (*p*-value= 0.497). Integration of the snATAC-seq and snRNA-seq resulted in the identification of 11 cell type clusters (Figure 1A) with no sample specific bias (Supplementary Figure 1A and 1B, Supplementary Table 2). The cell type identity of each cluster was determined using top 50 snATAC-seq predicted gene scores for marker genes of known cell types in the frontal cortex (Figure 1B). For example, we used *SLC17A7* for excitatory neurons; *SLC32A1* for inhibitory neurons; *MAG* for oligodendrocytes; *AIF1* for microglia; *GFAP* for astrocytes, *CSPG4* for oligodendrocyte precursor cell (OPCs) and *COL1A2* for vascular leptomeningeal cell (VLMC) (Supplementary Figure 2). We identified one oligodendrocyte cluster (55.6% of total cells), four excitatory neuron clusters (20.8% of the total cells), one inhibitory neuron cluster (8.3% of total cells), one microglia cluster (7.4% of total cells), one OPC cluster (3.9% of total cells), two astrocyte clusters (2.5% of total cells), and one vascular leptomeningeal cell (VLMC) cluster (1.5% of total cells) (Supplementary Table 2).

**Figure 1.**
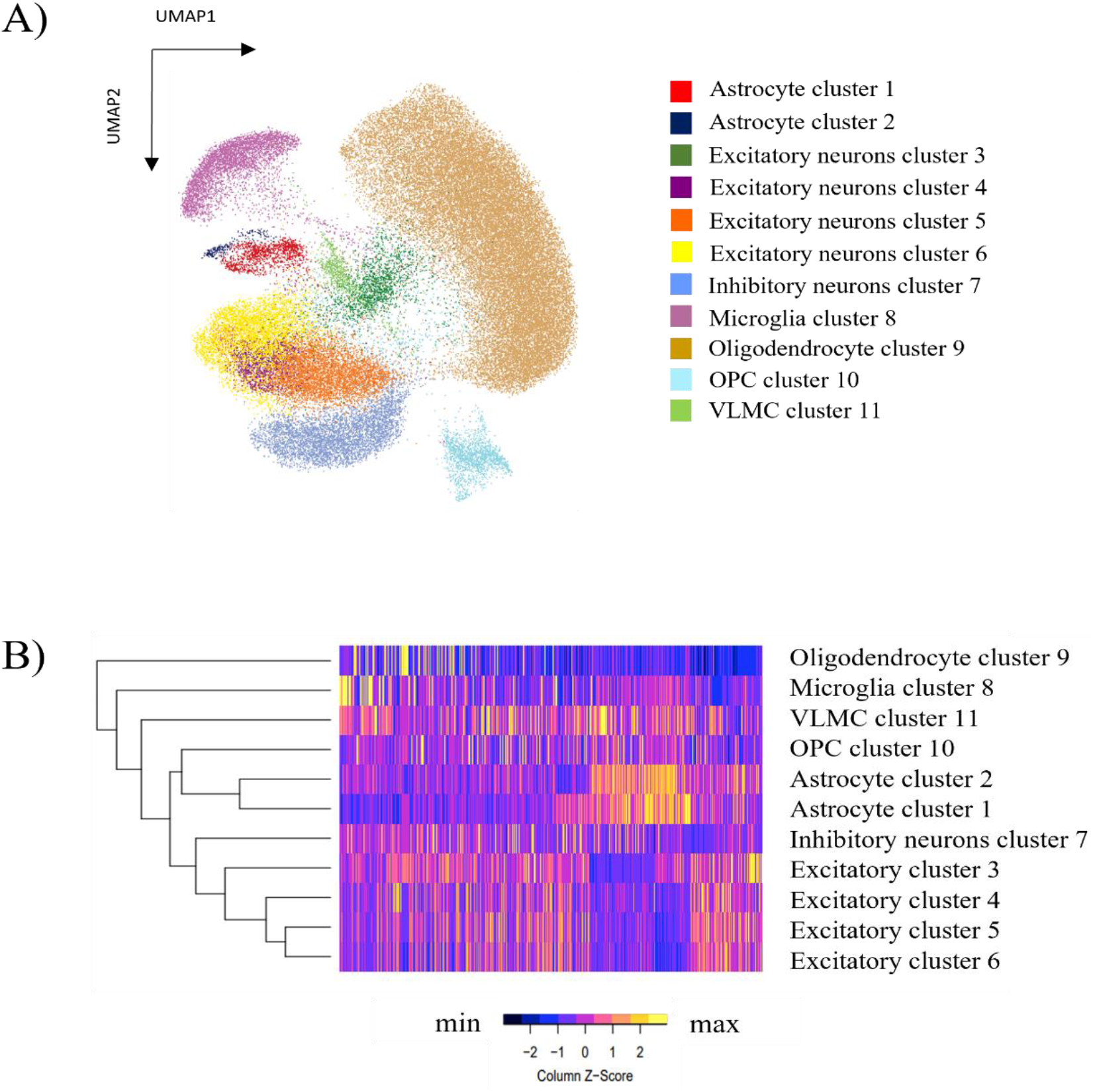
Visualization of the integrated single nuclei ATAC and RNA sequencing clusters and cell identification across the samples. A) UMAP reduction plot using resolution of 0.4 resulting in 11 cell type clusters from the integration of snATAC-seq and snRNA-seq data. B) Heatmap showing cell cluster identification by chromatin accessibility patterns using the top 50 snATAC-Seq predicted gene scores for markers genes of the known cortex cell types.

To further characterize the two astrocyte cell clusters, we determined the genes that identified astrocyte cluster 1 and astrocyte cluster 2 and investigated their expression in the single cell atlas of the Entorhinal Cortex in Human Alzheimer’s Disease (ECHAD)^53^ and astrocyte transcriptomic data from frontal cortex^54^. We observed that the genes characterizing astrocyte cluster 1 were primarily expressed in astrocyte subclusters a4 to a8, while the genes from our astrocyte cluster 2 were primarily expressed in astrocyte subclusters a2 and a3 from ECHAD, which correspond to reactive astrocytes subpopulations (Supplementary Figure 3)^54^.

### Accessibility analysis at the APOE locus and LA

We first investigated whether the ancestry-related differential expression of *APOE4* previously described^11^ was recapitulated when additional samples from Rush University Medical Center were included in our analyses. Indeed, *APOE4* was differentially expressed when considering expression over all cell types between ancestries with greater expression in ELA (FC= 1.31; *p*-value=9.66E^−219^), except one excitatory neurons cluster with higher expression in ALA (FC= 1.28; *p*-value=5.87E^06^). This pattern of increased expression in ELA samples was true for four cell types (excitatory and inhibitory neurons, astrocytes and microglia) (Supplementary Table 3). Astrocytes had the highest fold change difference in ELA (FC = 1.56) and most significant *p*-value (*p*-value=1.24E^−129^).

We next determined whether the *APOE* expression difference in astrocytes is associated with differences in chromatin accessibility. The snATAC-seq and snRNA-seq integrated UMAP showed that *APOE* expression as well as accessibility was highest in the astrocyte cluster 1 (Figure 2A). Comparison of accessibility between ancestries revealed two peaks with significantly increased accessibility in ELA at the *APOE* promoter in astrocyte cluster 1 (Figure 2B). These peaks were located at −19bp and −1990bp upstream of the *APOE* transcription start site (TSS) (FC: 2.57 and 4.52; FDR: 0.001 and 0.02, respectively). Notably, although *APOE* showed significantly increased expression in several cell types, significant differences in accessibility were only detected in astrocyte cluster 1 (Supplementary Figure 4).

**Figure 2.**
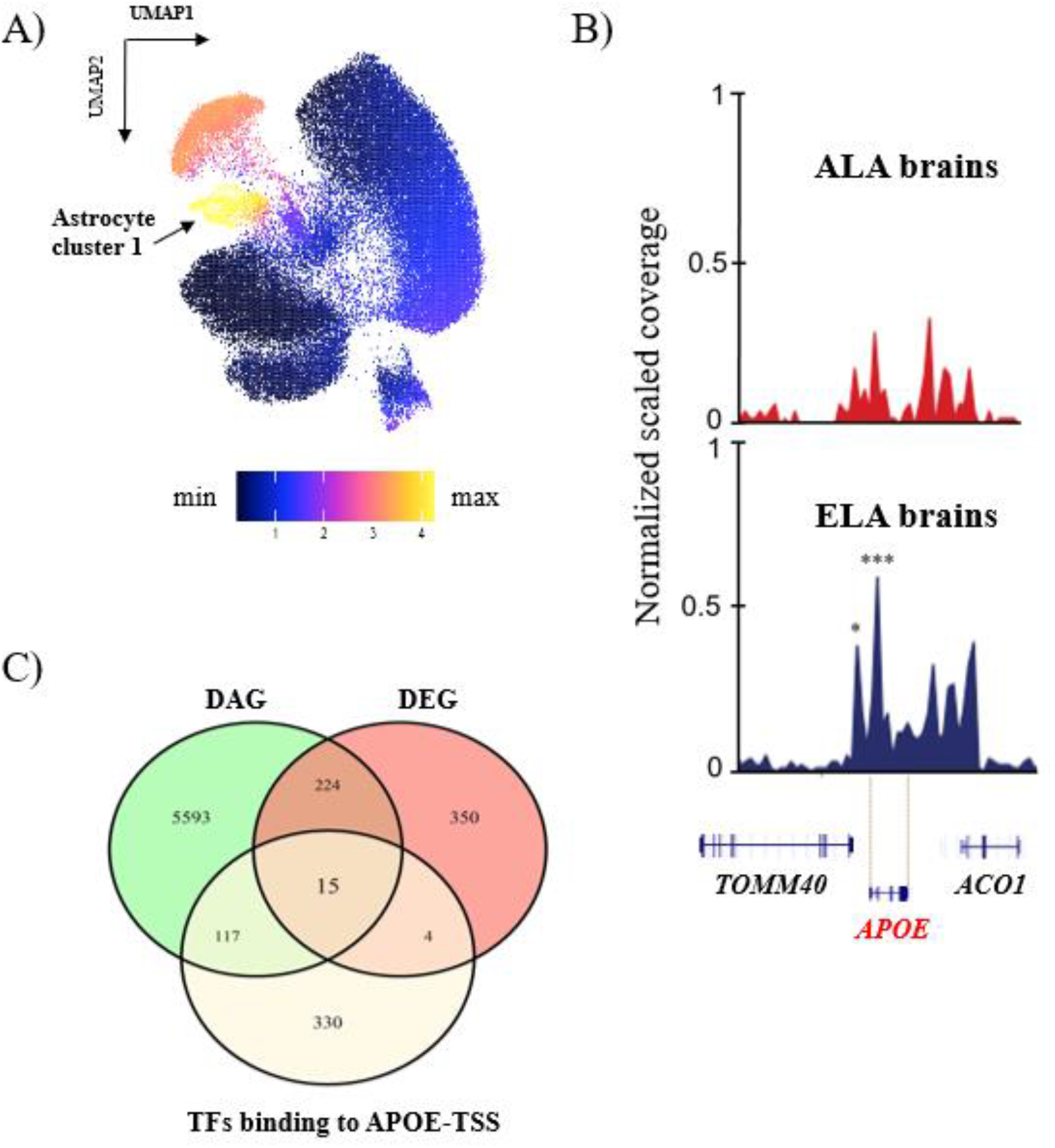
Differential chromatin accessibility and expression of *APOE* in Astrocyte cluster 1. A) *APOE* expression represented in clusters generated by the integrated snATAC-seq and snRNA-seq data; B) Visualization of chromatin differential accessible peaks in the *APOE* locus between ancestries from Astrocyte cluster 1. *Represents significantly differentially accessible peaks between ancestries (*FDR=0.02; ***FDR=0.001). C) Venn diagram showing transcription factors biding to *APOE* that are differentially expressed (DEG) and differentially accessible (DAG).

To gain insight into the molecular mechanisms involved in the increased accessibility and increased expression of *APOE*, we determined which known transcription factor binding sites were present in the differentially accessible peaks and analyzed the accessibility and gene expression of those transcription factors. The two differentially accessible peaks at the *APOE* promoter overlap predicted binding sites for a set of 15 transcription factors (*GLI2, NPAS2, KLF15, HIF1A, LHX2, RXRA, MXI1, FOS, JUNB, KLF2, PURA, SREBF1, TEAD1, KLF6, ZBTB7C*) that are also differentially expressed and have differentially accessible genomic region between ancestries (Supplemental Table 4). From these 15 putative *APOE* binding transcription factors, *PURA* and *SREBF1* have increased accessibility and expression in ELA samples, *KLF6* has increased accessibility and expression in ALA samples and the other 12 transcription factors have increased accessibility in ELA and increased expression in ALA.

We identified 32 additional differentially accessible peaks with greater accessibility in ELA than ALA astrocytes in gene promoters (within 2kb from the TSS) in the LA region surrounding *APOE* (chr19:44-46MB), which corresponds to 19 additional genes, including the *APOE* proximal genes *TOMM40* and *APOC1* (Table 2). No genes in the defined LA region were more accessible in the ALA. In addition, by overlapping the differentially accessible peaks in the *APOE* LA region with previously classified enhancers (ELITE GeneHancer from USCS) we identified 32 additional peaks among 23 LA genes, all with increased accessibility in ELA brains (Supplementary Table 5).

**Table 2.** Differentially accessible peaks in promoter regions of Local Ancestry genes in Astrocyte cluster 1.

| Gene | Chromosome | Peak location | Distance to TSS | FC | FDR | Region |
| --- | --- | --- | --- | --- | --- | --- |
| *CTB-171A8.1* | chr19 | 44718456 | -21 | 3.66 | 0.003 | promoter-TSS |
| *BCL3* | chr19 | 44748309 | 12 | 2.77 | 0.007 | promoter-TSS |
| *BCAM* | chr19 | 44808802 | -7 | 5.11 | 0.043 | promoter-TSS |
| *BCAM* | chr19 | 44808802 | -51 | 5.11 | 0.043 | promoter-TSS |
| *CTB-129P6.4* | chr19 | 44890205 | 421 | 2.70 | 0.002 | promoter-TSS |
| *TOMM40* | chr19 | 44890205 | -781 | 2.70 | 0.002 | promoter-TSS |
| *APOE\** | chr19 | 44903514 | -1990 | 4.52 | 0.022 | promoter-TSS |
| *APOE\** | chr19 | 44905485 | -19 | 2.57 | 0.001 | promoter-TSS |
| *APOC1* | chr19 | 44914897 | 278 | 2.05 | 0.000 | promoter-TSS |
| *APOC1* | chr19 | 44914897 | 900 | 2.05 | 0.000 | promoter-TSS |
| *APOC1P1* | chr19 | 44926992 | 266 | 2.23 | 0.025 | promoter-TSS |
| *CLPTM1* | chr19 | 44955080 | 84 | 3.61 | 0.007 | promoter-TSS |
| *CLPTM1* | chr19 | 44955080 | 84 | 3.61 | 0.007 | promoter-TSS |
| *CTB-179K24.3* | chr19 | 45091377 | -1236 | 2.19 | 0.027 | promoter-TSS |
| *PPP1R37* | chr19 | 45091377 | -1335 | 2.19 | 0.027 | promoter-TSS |
| *MARK4\** | chr19 | 45294609 | 504 | 3.65 | 0.011 | promoter-TSS |
| *ERCC2* | chr19 | 45370425 | -88 | 2.40 | 0.002 | promoter-TSS |
| *ERCC2* | chr19 | 45370425 | -57 | 2.40 | 0.002 | promoter-TSS |
| *PPP1R13L* | chr19 | 45405279 | -491 | 3.68 | 0.004 | promoter-TSS |
| *CD3EAP* | chr19 | 45405279 | -1162 | 3.68 | 0.004 | promoter-TSS |
| *PPP1R13L* | chr19 | 45405279 | 820 | 3.68 | 0.004 | promoter-TSS |
| *FOSB* | chr19 | 45467194 | -551 | 2.61 | 0.024 | promoter-TSS |
| *VASP* | chr19 | 45506348 | -818 | 3.58 | 0.020 | promoter-TSS |
| *OPA3* | chr19 | 45584758 | -166 | 2.16 | 0.000 | promoter-TSS |
| *EML2\** | chr19 | 45644874 | 505 | 7.63 | 0.032 | promoter-TSS |
| *EML2\** | chr19 | 45645399 | -20 | 3.28 | 0.002 | promoter-TSS |
| *FBXO46* | chr19 | 45730087 | 567 | 4.28 | 0.025 | promoter-TSS |
| *FBXO46* | chr19 | 45730692 | -38 | 2.01 | 0.028 | promoter-TSS |
| *FBXO46* | chr19 | 45730692 | -38 | 2.01 | 0.028 | promoter-TSS |
| *RSPH6A* | chr19 | 45815742 | -673 | 3.10 | 0.033 | promoter-TSS |
| *IRF2BP1* | chr19 | 45885449 | 471 | 4.72 | 0.025 | promoter-TSS |
| *NOVA2* | chr19 | 45973825 | -529 | 5.11 | 0.004 | promoter-TSS |
| *CCDC61* | chr19 | 45995053 | -158 | 2.38 | 0.000 | promoter-TSS |
| *CCDC61* | chr19 | 45995053 | -164 | 2.38 | 0.000 | promoter-TSS |

FC: Fold change; TSS: Transcription Starting Site; Negative value in Distance to TSS reflects peaks falling upstream of TSS

### Genome-wide differential accessibility and expression

Since the global ancestry of the ELA samples were predominantly European and the European local ancestry blocks are uniformly distributed among ALA samples (Supplementary Figure 5), we performed genome-wide differential accessibility analysis. In total we identified 5,154 significant (FDR ≤ 0.05 and FC ≥ 2) differentially accessible peaks between the ancestries in three cell types (astrocytes, excitatory neurons and microglia), representing less than 1% of all called ATAC peaks. 99% of these differentially accessible peaks were seen in the astrocyte cluster 1, with an overall increased chromatin accessibility in the European ancestry blocks compared to African ancestry blocks (more accessible in ELA: 4,546 peaks; more accessible in ALA: 107 peaks) (Figure 3A). This astrocyte specific accessibility difference was widely spread across all chromosomes (Figure 3B). Among all differentially accessible peaks in all chromosomes, 25.8% (2,248 peaks) are in promoters and TSS of genes, 33.5 % (2,920 peaks) are in exons, introns or close to the transcription terminating site (TTS), 34.1% (2,970 peaks) are in intergenic regions and 6.6% (574 peaks) are in distal ELITE enhancers (Figure 3C), altogether corresponding to 6,067 genes.

**Figure 3.**
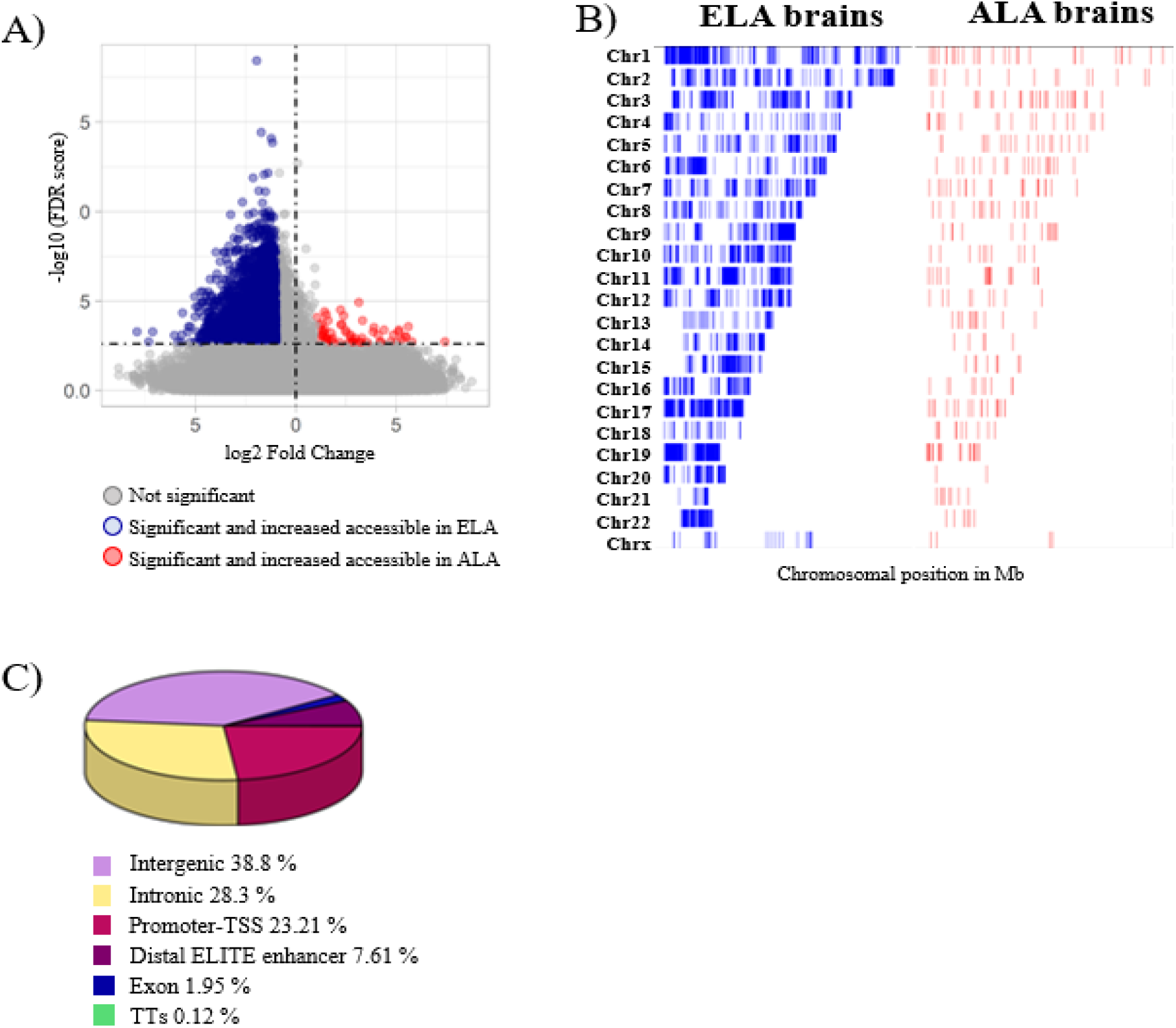
Chromatin accessibility differences between ancestries genome-wide in Astrocyte cluster 1. A) Volcano plot representation of global chromatin accessibility difference between ancestries in Astrocyte cluster 1; B) Chromatin accessibility differences across the genome displayed by chromosomes. Blue is increased accessibility in Europeans, Red is increased accessibility in African ancestry; C) Pie chart showing the differentially accessible peak region distribution in Astrocyte cluster 1.

Sixteen percent of the differentially accessible peaks were found on chromosome 19, a significant enrichment as compared to a random chromosomal distribution of peaks (Fischer exact test adjusted *p*-value=2,501E^−31^) with 36.4% of these differentially accessible peaks in the promoter regions of chromosome 19 genes (Supplementary Figure 6).

### Pathway enrichment of differentially accessible and expressed genes

To gain mechanistic insights from the integrative snRNA-seq and snATAC-seq approach, we determined the genome-wide overlap between DEG in the astrocyte clusters identified by snRNA-seq and genes associated with differentially accessible peaks identified by snATAC-seq (DAG). From the 6,067 genes with differentially accessible peaks, 239 genes were both differentially expressed and accessible (DEG-DAG) between the ancestries (Supplementary table 6). However only 55 genes had concordant differences in expression and accessibility: 48 with increased accessibility and expression in ELA (including *APOE*) and 7 with reduced expression and accessibility in ELA. This set of 55 DEG-DAG in astrocytes were used to compute a functional enrichment analysis.

The analysis of the KEGG Molecular functional pathways showed enrichment in signaling pathways including specifically: “Calcium signaling” (adjusted *p*-value=0.01) and “MAPK signaling pathway” (adjusted *p*-value=0.02). The GO gene-sets highlighted alterations in the biological process of cholesterol metabolism, including “regulation of cholesterol metabolic process” (GO:0090181, adjusted *p*-value=0.04) and “regulation of cholesterol biosynthetic process” (GO:0045540, adjusted *p*-value=0.04), pathways linked to synapses and axonal transport (GO:0097106, GO:0090129 and GO:0090128 adjusted *p*-value 0.03; GO:0008090, GO:0001941, GO:0045773, GO:0150104 adjusted *p*-value=0.04), spine development (GO:0060999, GO:0060998 adjusted *p*-value=0.04) and pathways linked to glial cell and astrocyte projection (GO:0097386, GO:0097449 adjusted *p*-value 0.02) (Supplementary Table 7).

Using databases of disease associated genes from published transcriptomic studies revealed the highest overlap with lipopolysaccharide (LPS) treated astrocytes to induce reactivity (adjusted *p*-value =8.16E^−10^, GSE75246)^55^ and with genes showing altered expression in brains of Huntington Disease (HD) patients (adjusted *p*-value =2.26E^−19^, GSE79666)^56^. Thus, we tested the enrichment in the HD-relevant database HDSigDB and found significant enrichment in gene changes specifically in astrocytes of HD patients (adjusted *p*-value =2.92E^−19^)^57^, particularly with genes up-regulated in astrocytes of HD patients (adjusted *p*-value=2.45E^−17^)^57^. The overlapping genes between the ancestry DEG-DAG and genes up-regulated in astrocytes of HD patients include a molecular signature of reactive astrocyte markers also observed in the HD striatal astroglia data, but also extends to pathways related to glutamatergic synapse (GO:0098978) and signaling (hsa04020; hsa04015; hsa04022). In addition, the ENCODE database of brain histone post-translational modifications identified an enrichment of DEG-DAG genes in those genes having H3K27me3 signals in astrocytes (H3K27me3 astrocytes Hg19, adjusted *p*-value = 0.02) (Supplementary Table 7).

### Chromatin accessibility in AD candidate genes

We also assessed chromatin accessibility differences between the ancestries in AD associated genes, including Mendelian genes (*APP*, *PS1*, *PS2*, and *MAPT*) and genes suggested by GWAS and rare variant association studies across ancestries^58,59^. 36 differentially accessible peaks between the ancestries were identified in eight out of 140 AD-associated genes analyzed. Aside from *APOE*, nine differentially accessible peaks were observed to fall in the promoter areas of three genes, *SORL1, VRK3* and *ABCA7*, all with increased accessibility in ELA regions (Table 3). Ten differentially accessible peaks fall in intergenic regions close to seven AD genes, all with increase accessibility in ELA. In addition, fifteen peaks are found in the gene body of four AD-associated genes (*CLU, ACER3, SORL1, SPHK1*), also with increased accessibility in ELA (Table 3). Nonetheless, none of these genes containing differentially accessible peaks between local ancestries showed differentially expression between ancestries.

**Table 3.** Differentially accessible peaks in Astrocyte cluster 1 in Alzheimer disease GWAS candidate genes.

| Gene | Chromosome | Peak location | Cell cluster | Distance to TSS | FC | FDR | Region |
| --- | --- | --- | --- | --- | --- | --- | --- |
| *PTK2B* | chr8 | 27340471.00 | Astrocyte 1 | 29239 | 2.33 | 0.026 | intron |
| *CLU* | chr8 | 27596089.00 | Astrocyte 1 | 18692 | 4.23 | 0.019 | intergenic |
| *CLU* | chr8 | 27597084.00 | Astrocyte 1 | 17697 | 2.56 | 0.035 | exon |
| *CLU* | chr8 | 27597084.00 | Astrocyte 1 | 17697 | 2.56 | 0.035 | intron |
| *CLU* | chr8 | 27597917.00 | Astrocyte 1 | 16864 | 18.11 | 0.008 | exon |
| *CLU* | chr8 | 27597917.00 | Astrocyte 1 | 16864 | 18.11 | 0.008 | intron |
| *CLU* | chr8 | 27607577.00 | Astrocyte 1 | 7204 | 6.69 | 0.032 | intron |
| *ACER3* | chr11 | 76799174.00 | Astrocyte 1 | -61449 | 2.41 | 0.008 | intergenic |
| *SORL1* | chr11 | 121566449.00 | Astrocyte 1 | 114496 | 3.54 | 0.044 | exon |
| *SORL1* | chr11 | 121566449.00 | Astrocyte 1 | 114496 | 3.54 | 0.044 | intron |
| *SORL1* | chr11 | 121566449.00 | Astrocyte 1 | -26 | 3.54 | 0.044 | promoter-TSS |
| *SORL1* | chr11 | 121591362.00 | Astrocyte 1 | 139409 | 7.33 | 0.011 | intron |
| *SORL1* | chr11 | 121591362.00 | Astrocyte 1 | 1193 | 7.33 | 0.011 | promoter-TSS |
| *SORL1* | chr11 | 121596337.00 | Astrocyte 1 | 6168 | 6.49 | 0.046 | intron |
| *SORL1* | chr11 | 121596337.00 | Astrocyte 1 | 144384 | 6.49 | 0.046 | intron |
| *SORL1* | chr11 | 121695992.00 | Astrocyte 1 | 105823 | 3.06 | 0.039 | intergenic |
| *SORL1* | chr11 | 121695992.00 | Astrocyte 1 | 244039 | 3.06 | 0.039 | intergenic |
| *SORL1* | chr11 | 121795198.00 | Astrocyte 1 | 205029 | 3.10 | 0.027 | intergenic |
| *SORL1* | chr11 | 121795198.00 | Astrocyte 1 | 343245 | 3.10 | 0.027 | intergenic |
| *SORL1* | chr11 | 121929505.00 | Astrocyte 1 | 477552 | 3.49 | 0.004 | intergenic |
| *SORL1* | chr11 | 121932043.00 | Astrocyte 1 | 480090 | 11.93 | 0.031 | intergenic |
| *SPHK1* | chr17 | 76397939.00 | Astrocyte 1 | 13539 | 4.18 | 0.041 | intergenic |
| *SPHK1* | chr17 | 76397939.00 | Astrocyte 1 | 12897 | 4.18 | 0.041 | intron |
| *SPHK1* | chr17 | 76414406.00 | Astrocyte 1 | 30006 | 15.90 | 0.000 | intergenic |
| *ABCA7* | chr19 | 1039766.00 | Astrocyte 1 | -87 | 2.43 | 0.001 | promoter-TSS |
| *ABCA7* | chr19 | 1040877.00 | Astrocyte 1 | -119 | 4.09 | 0.002 | promoter-TSS |
| *ABCA7* | chr19 | 1040877.00 | Astrocyte 1 | 1024 | 4.09 | 0.002 | promoter-TSS |
| *VRK3* | chr19 | 50024717.00 | Astrocyte 1 | 360 | 5.86 | 0.043 | promoter-TSS |
| *VRK3* | chr19 | 50024717.00 | Astrocyte 1 | 979 | 5.86 | 0.043 | promoter-TSS |
| *VRK3* | chr19 | 50025268.00 | Astrocyte 1 | -116 | 3.41 | 0.002 | promoter-TSS |
| *VRK3* | chr19 | 50025268.00 | Astrocyte 1 | 428 | 3.41 | 0.002 | promoter-TSS |
| *VRK3* | chr19 | 50050285.00 | Astrocyte 1 | -24589 | 5.27 | 0.002 | intergenic |

FC: Fold change; TSS: Transcription Starting Site; Negative value in Distance to TSS reflects peaks falling upstream of TSS

## DISCUSSION

To understand the complex mechanisms by which *APOE4* confers differential risk for AD in the context of genetic ancestry, we evaluated both gene expression and chromatin accessibility profile differences between homozygous *APOE4/4* ELA and ALA Alzheimer disease brains at single cell resolution. We confirmed our previous finding^11^ that *APOE* is significantly more expressed in astrocytes of ELA compared to ALA. In fact, *APOE* is the most differentially expressed gene in astrocytes in the 2Mb local ancestry region surrounding *APOE*. Beyond being the most differentially expressed, the promoter of *APOE* in astrocytes showed a significant increase in accessibility in ELA compared to ALA. These results suggest that the increased *APOE4* expression previously reported in ELA *APOE4* carriers^11^ may be partly due to differences in chromatin remodeling at the promoter of *APOE4* between ancestries. We have also shown using Capture Chromatin Conformation analysis and massively parallel reporter assays that specific DNA sequence differences in areas physically interacting with the *APOE* promoter have functional effects in microglia and astrocytes, with greater expression occurring in ELA sequence variants than ALA^60^. Thus, given the long-time span of AD, it appears that multiple mechanisms affecting *APOE4* gene expression in astrocytes may be activated at different times in life or by stress and could affect the differences in AD risk seen between ancestral homozygous *APOE4* carriers. This emphasizes that *APOE* loci is under a complex regulatory environment that involves cell type-specific processes and strengthen the primary functional role of *APOE* in astrocytes^61^.

Though our principal research question focused on the underlying mechanisms leading to the higher expression of *APOE* in ELA brains using snATAC-seq, this study also provides a global view on the chromatin landscape of AD *APOE4/4* carriers from different genetic ancestries. We observed in astrocyte cluster 1 a significantly higher chromatin accessibility in ELA brains genome wide. Notably, only a small percent of those significant peaks of higher chromatin accessibility overlap with the significant differentially expressed genes identified by snRNA-seq. Conversely, only 40% of the significant differentially expressed genes showed significant changes in accessibility between the ancestries. This is in accordance with the growing understanding that gene expression depends on multiple regulatory elements^62^, many of which are located far from the gene locus itself^23^. Some of the DEG-DAG show a positive correlation between expression and accessibility, suggesting a functional relationship. However, we also observed an anti-correlation in accessibility-expression pairs. This anti-correlation could represent binding by repressors leading to a decrease in expression as previously described^63,64^. In addition, highly accessible sites not associated with enhanced expression could represent poised enhancers/promoter with no active transcription occurring^65^. Besides this known biological variability in accessibility and expression correlation in this study the snATAC and snRNA were derived from separate, but adjacent, brain punches. Thus, differences in the tissue used could contribute to some discordance between these two outcomes. Future studies with paired snATAC-seq and snRNA-seq from the same nuclei such as the 10X Multiome will minimize any of these potential tissue location effects, if present. In fact, while we do identify differential expression of *APOE* between ancestries in several cell types (e.g., microglia), we only see significant differential accessibility of its promoter and immediate region in astrocytes. Finally, a recent study using mesenchymal progenitor cells has reported that *APOE* accumulation in the nucleus can disrupt and destabilize heterochromatin, which would increase chromatin accessibility as we have seen here in the ELA with its higher *APOE* expression^66^. As astrocytes are the primary cell with *APOE* expression, this could explain the finding of increased global accessibility primarily in this cell type.

Astrocytes have well established roles in providing neuronal metabolic support and neuroprotection, but also play important roles in synapse formation and function, ion signaling homeostasis, integrity of the blood brain barrier, tissue repair and complex brain functions such as memory and sleep^67–71^. Thus, it is not surprising that a growing body of evidence suggests a role for astrocytes in AD pathology^72^. Our data suggest that astrocytes are involved in the modulation of *APOE4*-afforded AD risk and pinpoint this cell type as one of the most sensitive to differences in genetic ancestry in our study. The significant enrichment of astrocytic ancestry-associated DEG-DAG in genes upregulated in HD astrocytes suggest that the observed differences associated with diverse ancestry entail functional properties that are potentially common to other neurodegenerative disorders. In addition to this astrocyte-ancestry association, we also evidence a difference in DEG-DAG between our astrocyte clusters 1 and 2, when we use astrocyte subclusters described by Grubman et al^53^ and also reference genes identified by Sadick et al^54^. We observed that our astrocyte cluster 1 was most similar in genetic expression to astrocytes subclusters a4 and a8, clusters where Grubman et al^53^ also found that *APOE* was upregulated in AD; while our astrocyte subcluster 2 was comparable in genetic expression to astrocytes subclusters a2 where *APOE* was downregulated, further supporting our findings.

Interestingly, there was a significant enrichment of differential chromatin accessibility and expression on chromosome 19 relative to the rest of the genome (Supplemental Figure 5) in astrocyte cluster 1. These results correlate with the observation of a comparatively high number of ATAC-seq peaks in chromosome 19^73^, potentially due to the high gene density seen on chromosome 19, GC content and DNA binding proteins compared to other chromosomes in the genome^74–76^. It is possible the high gene density and the structural nature of chromosome 19 could modulate the differential binding of chromatin modifying enzymes, which may explain the enrichment in differentially accessible chromatin and gene expression in this specific chromosome observed between ancestries.

Our analysis of transcription factors with binding sites in the differentially accessible peaks at the promoter and upstream of *APOE* that were both differentially expressed and accessible between ancestries suggests transcriptional modulators of *APOE* expression differences. Of the 15 transcription factors we identified, only *SREBF1* and *PURA* showed increased accessibility (intergenic and intronic) and expression in ELA samples and *KLF6* showed increased accessibility (distal enhancer) and expression in ALA samples. A polymorphism in *SREBF1* has been shown to influence the risk of AD specifically in *APOE4* carriers^77^, while *PURA* has been shown to regulate expression of AD and Aβ clearance related genes^78^. On the other hand, *KLF6* is found to be differentially expressed in AD brains and implicated in Aβ-induced oxidative stress^79^. Their role with *APOE* activation has not been previously described, although *SREBF1* is a major regulator of lipid metabolism.

Although many genetic risk factors for disease are expected to be shared among populations, genomic diversity among ancestries can provide new opportunities for discoveries regarding disease susceptibility. In the case of AD, understanding why the *APOE4* allele confers a lower risk to African ancestry carriers versus European ancestry carriers is instrumental for the identification of potential AD therapeutics. Here we provide chromatin accessibility and gene expression data from AD *APOE4* carriers of African and European ancestries at single cell resolution. Expansion of this study into other *APOE* genotypes and more diverse ancestries could provide further insight into the importance of brain astrocytes and the differential accessibility and expression of *APOE*, as well as genome-wide changes, implicating specific molecular pathways and regulatory proteins in this risk difference. Finally, these findings support the concept of reducing *APOE4* expression as a potential therapeutic pathway for AD.

## CONCLUSIONS

Our results provide novel compelling insights to understand the AD risk difference known to exist between European and African *APOE4* local ancestry carriers. Here we demonstrate that differences in chromatin accessibility between the African and European *APOE* locus could explain the expression differences in *APOE* expression in astrocytes and supports the concept that this contributes to the risk difference between African American and European populations for AD in *APOE4* carriers. We found that this increase in accessibility in the region surrounding *APOE4* in European astrocytes extended genome-wide, suggesting a wider mechanism of regulatory dysfunction is occurring in these cells. Not only is it important to include diverse ancestries to ensure all individuals are represented in research, but also we have shown here and in previous work that including diversity in research can provide additional windows of opportunity to elucidate disease and biological mechanisms. Understanding regulatory dysfunction in AD has not been well studied and should be and area of future research in AD.

## LIST OF ABBREVIATIONS

AD: Alzheimer Disease
ABCA7: ATP Binding Cassette Subfamily A Member 7
ACER3: Alkaline Ceramidase 3
ADRC: Alzheimer Disease Research Centers
AIF1: Allograft inflammatory factor 1
ALA: African Local Ancestry
APOC1: Apolipoprotein C1
APOE: Apolipoprotein E
APOE4: Apolipoprotein E allele 4
APOE4/4: Homozygous for Apolipoprotein E allele 4
APP: Amyloid Beta Precursor Protein
ATAC: Assay for Transposase-Accessible Chromatin
Azimth: App for reference-based single-cell analysis
Aβ: Amyloid beta
BRAAK: Braak staging (pathology score for Alzheimer Disease)
CGT: Center for Genome Technology
CLU: Clusterin
COL1A2: Collagen Type I Alpha 2 Chain
CSPG4: Chondroitin Sulfate Proteoglycan 4
DAG: Differentially Accessible Genes
DAP: Differentially Accessible Peaks
DEG: Differentially Expressed Genes
DEG-DAG: Differentially Expressed and Accessible Genes
ECHAD: Entorhinal Cortex in Human Alzheimer’s Disease
ELA: European Local Ancestry
ELITE: enhancer gene with both a high likelihood enhancer definition and a strong enhancer–gene association
ENCODE: Encyclopedia of DNA Elements
FC: Fold Change
FDR: false discovery rate
FOS: Fos Proto-Oncogene, AP-1 Transcription Factor Subunit
GC: Guanine-Cytosine content
GeneSigDb: Gene Signature Data Base
GFAP: Glial Fibrillary Acidic Protein
GLI2: GLI Family Zinc Finger 2
GO: Gene Ontology
GRCh38: Genome Reference Consortium Human Build 38
GREAT: Genomic Regions Enrichment of Annotations Tool
H3K27me3: tri-methylation of lysine 27 on histone H3 protein
HD: Huntington Disease
HDSigDb: Huntigton Disease Signature Database
HIF1A: Hypoxia Inducible Factor 1 Subunit Alpha
HIHG: John P. Hussman Institute for Human Genomics
HOMER: Hypergeometric Optimization of Motif EnRichment
JASPAR: transcription factor binding profile database
JOLMA: Human specific transcription factor binding
JUNB: JunB Proto-Oncogene, AP-1 Transcription Factor Subunit
KEGG: Kyoto Encyclopedia of Genes and Genomes
KLF15: Kruppel Like Factor 15
KLF2: Kruppel Like Factor 2
KLF6: Kruppel Like Factor 6
LA: Local Ancestry
LHX2: LIM Homeobox 2
LPS: lipopolysaccharide
LSI: Iterative Latent Semantic Indexing
MAG: Myelin Associated Glycoprotein
MAPT: microtubule-associated protein tau
MB: Million Base pair
Mb: Mega base
MsigDB: Molecular Signatures Database
MXI1: MAX interactor-1
NACC: National Alzheimer Coordinating Center
NPAS2: Neuronal PAS Domain Protein 2
OPCs: oligodendrocyte precursor cells
PanglaoDB: Single-cell sequencing mouse and human database
PCR: Polymerase chain reaction
PSEN1: Presenillin-1
PSEN2: Presenillin-2
PURA: Purine Rich Element Binding Protein A
Reactome: pathway database
RSB: Resuspension Buffer
RXRA: Retinoid X Receptor Alpha
SLC17A7: Solute Carrier Family 17 Member 7
SLC32A1: Solute Carrier Family 32 Member 1
snATAC-seq: single nuclei Assay for Transposase-Accessible Chromatin sequencing
snRNA-seq: single nuclei RNA sequencing
SORL1: Sortilin Related Receptor 1
SPHK1: Sphingosine Kinase 1
SREBF1: Sterol Regulatory Element Binding Transcription Factor 1
TEAD1: TEA Domain Transcription Factor 1
TF: Transcription Factor
TOMM40: Translocase Of Outer Mitochondrial Membrane 40
TREM2: Triggering Receptor Expressed On Myeloid Cells 2
TSS: Transcription Starting Site
TTS: transcription terminating site
UCSC: University of California Santa Cruz
USUHS: Uniformed Services University of the Health Sciences
VLMC: vascular leptomeningeal cell
VRK3: VRK Serine/Threonine Kinase 3
ZBTB7C: Zinc Finger And BTB Domain Containing 7C
WGS: Whole Genome Sequencing

## DECLARATIONS

### Ethics approval and consent to participate

We consent for publication.

### Consent for publication

We consent for publication.

### Availability of data and material

Data are available through the National Institute on Aging Genetics of Alzheimer’s Disease Data Storage Site (NIAGADS) Data Sharing Service (DSS): https://dss.niagads.org/datasets/ng00067/

### Author contributions

K.C, J.M.V, J.I.Y and A.J.G overall study design. K.C performed experiments. M.D.M.M and A.J.G performed bioinformatics analysis. F.R and K.H performed ancestry analysis. P.W. and C. L.D performed genotyping arrays and whole genome sequencing. M.F, S.W, C.G, M.G, D. A.B, T.S and M.A.P provided brain samples and advice. D.M.D, K.N, L.W and F.J provided insight for the functional analysis. K.C, J.M.V, J.I.Y, A.J.G and M.D.M.M wrote the manuscript with input from all authors. All authors read and approved the final manuscript.

## Acknowledgements

We acknowledge the collaboration of Dr. John Q. Trojanowski in this study, whose untimely death is a great loss to both his family, colleagues and to the field of neurodegeneration research.

## Supplementary Figure Legends

**Supplementary Figure 1.** SnATAC-seq data analysis for sample bias.

**Supplementary Figure 2.** SnATAC-seq cell marker cluster identification by chromatin accessibility.

**Supplementary Figure 3.** Astrocyte subtype classification.

**Supplementary Figure 4.** *APOE* chromatin accessibility in the other cell clusters where *APOE* is highly expressed but not differentially accessible between the ancestries.

**Supplementary Figure 5.** Visualization of local ancestry across all chromosomes per individual sample.

**Supplementary Figure 6.** Enrichment of differentially accessible peak between ancestries in chromosome 19.

## Supplementary Table Legends

**Supplementary Table 1.** Summary of snATAC and snRNA sequencing results.

**Supplementary Table 2.** Total cell count and proportion per cluster in the integrated sequencing analysis.

**Supplementary Table 3.** snRNA sequencing clusters with significant *APOE4* differential expression between ancestries.

**Supplementary Table 4.** TFs (DEGs-DAGs) binding to differentially accessible peaks at the promoter of *APOE*.

**Supplementary Table 5.** *APOE* Local ancestry accessibility analysis.

**Supplementary Table 6.** Differentially accessible and expressed genes (DEG-DAG) in astrocyte cluster 1.

**Supplementary Table 7.** Subset of the differential functional analysis of the astrocyte cluster 1 DEG-DAG with positive correlation between expression and accessibility using EnrichR software.

