## supplementary material for "Ancestry-related differences in chromatin accessibility and gene expression of *APOE4* are associated with Alzheimer disease risk"

**Supplementary Table 1.** Summary of snATAC and snRNA sequencing results.

| SnATAC-seq | | | | | SnRNA-seq | | |
|---|---|---|---|---|---|---|---|
| Sample | Raw # of cells | Median # of fragments per nucleus | Fraction of fragments overlapping any targeted region | Fraction of transposition events in peaks in cell barcodes | Raw # of cells | Median # of reads per nucleus | Median # of genes per nucleus |
| 1 | 7393 | 9270 | 0.492 | 0.176 | 6660 | 126054 | 2677 |
| 2 | 3571 | 6327 | 0.431 | 0.075 | 5984 | 180736 | 1331 |
| 3 | 5717 | 23659 | 0.532 | 0.214 | 3233 | 322773 | 1388 |
| 4 | 3553 | 837 | 0.475 | 0.166 | 12472 | 78945 | 1820 |
| 5 | 2421 | 14581 | 0.481 | 0.186 | 6599 | 101072 | 1718 |
| 6 | 8556 | 10863 | 0.541 | 0.234 | 16506 | 66903 | 2702 |
| 7 | 2336 | 21660 | 0.569 | 0.255 | 8853 | 86528 | 1253 |
| 8 | 4773 | 8838 | 0.536 | 0.256 | 10437 | 94609 | 2553 |
| 9 | 4842 | 9742 | 0.463 | 0.13 | 7100 | 126487 | 1759 |
| 10 | 1030 | 14782 | 0.515 | 0.205 | 10,953 | 62,612 | 1664 |
| 11 | 5375 | 10312 | 0.52 | 0.193 | 10492 | 96055 | 3332 |
| 12 | 832 | 16893 | 0.451 | 0.105 | 9643 | 110324 | 2395 |

#: number

**Supplementary Table 2.** Total cell count and proportion per cluster in the integrated sequencing analysis.

| | | AF | | | | | | | EU | | | | | | | | | |
|---|---|---|---|---|---|---|---|---|---|---|---|---|---|---|---|---|---|---|
| Cluster | Cell Type | SAMP 1 | SAMP2 | SAMP3 | SAMP4 | SAMP5 | SAMP6 | AF proportion | SAMP7 | SAMP8 | SAMP9 | SAMP10 | SAMP11 | SAMP12 | EU proportion | AF/EU ratio | Total nuclei | Total proportion of nuclei |
| 1 | Astrocytes | 210 | 137 | 82 | 13 | 74 | 151 | 0.547 | 153 | 48 | 93 | 20 | 216 | 23 | 0.453 | 1.206 | 1220 | 2.1 |
| 2 | Astrocytes | 29 | 51 | 65 | 0 | 34 | 1 | 0.804 | 0 | 4 | 2 | 1 | 0 | 37 | 0.196 | 4.091 | 224 | 0.4 |
| 3 | Excitatory Neurons | 21 | 574 | 38 | 7 | 35 | 176 | 0.515 | 19 | 51 | 208 | 8 | 451 | 65 | 0.485 | 1.061 | 1653 | 2.8 |
| 4 | Excitatory Neurons | 86 | 14 | 7 | 13 | 14 | 385 | 0.524 | 264 | 62 | 31 | 32 | 72 | 10 | 0.476 | 1.102 | 990 | 1.7 |
| 5 | Excitatory Neurons | 936 | 629 | 299 | 100 | 273 | 896 | 0.567 | 115 | 503 | 779 | 236 | 189 | 574 | 0.433 | 1.308 | 5529 | 9.4 |
| 6 | Excitatory Neurons | 916 | 136 | 46 | 77 | 50 | 729 | 0.487 | 42 | 207 | 675 | 87 | 922 | 126 | 0.513 | 0.949 | 4013 | 6.8 |
| 7 | Inhibitory Neurons | 571 | 323 | 291 | 201 | 126 | 619 | 0.437 | 413 | 302 | 924 | 238 | 638 | 230 | 0.563 | 0.776 | 4876 | 8.3 |
| 8 | Microglia | 572 | 553 | 342 | 73 | 137 | 525 | 0.506 | 325 | 486 | 333 | 64 | 642 | 299 | 0.494 | 1.025 | 4351 | 7.4 |
| 9 | Oligodendrocytes | 4859 | 2804 | 3982 | 553 | 2023 | 4847 | 0.582 | 942 | 3373 | 2387 | 453 | 2728 | 3796 | 0.418 | 1.394 | 32747 | 55.8 |
| 10 | OPC | 70 | 257 | 236 | 57 | 75 | 401 | 0.484 | 242 | 122 | 274 | 95 | 251 | 184 | 0.516 | 0.938 | 2264 | 3.9 |
| 11 | VLMC | 42 | 74 | 59 | 22 | 21 | 81 | 0.350 | 45 | 54 | 79 | 11 | 321 | 45 | 0.650 | 0.539 | 854 | 1.5 |

AF: African; EU: European; SAMP: sample

**Supplementary Table 3.** snRNA sequencing clusters with significant *APOE4* differential expression between ancestries.

| Gene | FC | FDR | Cell type |
|---|---|---|---|
| *APOE* | 1.556 | 1.24E-129 | Astrocytes |
| *APOE* | 1.392 | 4.62E-07 | Astrocytes |
| *APOE* | 1.399 | 3.30E-73 | Excitatory Neurons |
| *APOE* | -1.209 | 4.73E-23 | Excitatory Neurons |
| *APOE* | 1.191 | 2.48E-02 | Excitatory Neurons |
| *APOE* | 1.289 | 5.87E-06 | Excitatory Neurons |
| *APOE* | 1.200 | 2.35E-02 | Inhibitory Neurons |
| *APOE* | 1.273 | 1.41E-03 | Inhibitory Neurons |
| *APOE* | 1.433 | 6.58E-03 | Inhibitory Neurons |
| *APOE* | 1.415 | 3.89E-02 | Microglia |

FC: fold change; Negative FC reflects higher expression in African local ancestry brains

**Supplementary Table 4.** TFs (DEGs-DAGs) binding to differentially accessible peaks at the promoter of *APOE*

| Gene | Chromosome | Peak location | Cell cluster | Distance to TSS | FC (snATAC-seq) | FDR (snATAC-seq) | Region | Differentially | FC (snRNA-seq) | FDR (snRNA-seq) | ELITE enhancer (GeneHancer Db) | Gene Association Methods ELITE enhancer |
|---|---|---|---|---|---|---|---|---|---|---|---|---|
| *GLI2* | chr2 | 120758032 | Astrocyte 1 | -34127 | 4.363519487 | 0.002 | intergenic; intronic | DEGs, DAGs | -1.21 | 5.592E-53 | NEG | NEG |
| *NPAS2* | chr2 | 100818483 | Astrocyte 1 | -1419 | 2.520744875 | 0.050 | promoter-TSS | DEGs, DAGs | -1.55 | 4.592E-57 | NEG | NEG |
| *KLF15* | chr3 | 126341652 | Astrocyte 1 | 15540 | 2.696252091 | 0.035 | TTS; intergenic | DEGs, DAGs | -1.21 | 1.078E-55 | NEG | NEG |
| *KLF15* | chr3 | 126277893 | Astrocyte 1 | 79299 | 7.947060238 | 0.008 | intergenic | DEGs, DAGs | -1.21 | 1.078E-55 | NEG | NEG |
| *KLF15* | chr3 | 126208618 | Astrocyte 1 | 148574 | 4.357686047 | 0.024 | intergenic | DEGs, DAGs | -1.21 | 1.078E-55 | NEG | NEG |
| *KLF15* | chr3 | 126213000 | Astrocyte 1 | 144192 | 2.000902751 | 0.035 | intergenic | DEGs, DAGs | -1.21 | 1.078E-55 | NEG | NEG |
| *LHX2* | chr9 | 124011402 | Astrocyte 1 | 42 | 2.38695679 | 0.000 | promoter-TSS | DEGs, DAGs | -1.39 | 7.392E-66 | NEG | NEG |
| *LHX2* | chr9 | 124013712 | Astrocyte 1 | 2352 | 4.449551054 | 0.042 | exon; intron | DEGs, DAGs | -1.39 | 7.392E-66 | NEG | NEG |
| *LHX2* | chr9 | 124018110 | Astrocyte 1 | 6750 | 2.503939163 | 0.049 | intron | DEGs, DAGs | -1.39 | 7.392E-66 | NEG | NEG |
| *LHX2* | chr9 | 124188229 | Astrocyte 1 | 176869 | 5.029629361 | 0.010 | intergenic | DEGs, DAGs | -1.39 | 7.392E-66 | NEG | NEG |
| *LHX2* | chr9 | 124042056 | Astrocyte 1 | 30696 | 2.878977792 | 0.010 | intergenic | DEGs, DAGs | -1.39 | 7.392E-66 | NEG | NEG |
| *LHX2* | chr9 | 124110256 | Astrocyte 1 | 98896 | 4.70672574 | 0.024 | intergenic | DEGs, DAGs | -1.39 | 7.392E-66 | NEG | NEG |
| *LHX2* | chr9 | 124223400 | Astrocyte 1 | 212040 | 3.315235163 | 0.037 | intergenic | DEGs, DAGs | -1.39 | 7.392E-66 | NEG | NEG |
| *RXRA* | chr9 | 134348401 | Astrocyte 1 | 22071 | 3.62378105 | 0.006 | intron | DEGs, DAGs | -1.22 | 5.850E-67 | NEG | NEG |
| *RXRA* | chr9 | 134344653 | Astrocyte 1 | 18323 | 2.216024313 | 0.007 | intron | DEGs, DAGs | -1.22 | 5.850E-67 | NEG | NEG |
| *RXRA* | chr9 | 134331104 | Astrocyte 1 | 4774 | 2.188097796 | 0.016 | intron | DEGs, DAGs | -1.22 | 5.850E-67 | NEG | NEG |
| *RXRA* | chr9 | 134338784 | Astrocyte 1 | 12454 | 3.39686537 | 0.019 | intron | DEGs, DAGs | -1.22 | 5.850E-67 | NEG | NEG |
| *RXRA* | chr9 | 134326812 | Astrocyte 1 | 482 | 7.823652651 | 0.023 | promoter-TSS | DEGs, DAGs | -1.22 | 5.850E-67 | NEG | NEG |
| *RXRA* | chr9 | 134316329 | Astrocyte 1 | -10001 | 3.429170767 | 0.039 | Intergenic | DEGs, DAGs | -1.22 | 5.850E-67 | NEG | NEG |
| *RXRA* | chr9 | 134340714 | Astrocyte 1 | 14384 | 15.34732166 | 0.043 | intron | DEGs, DAGs | -1.22 | 5.850E-67 | NEG | NEG |
| *RXRA* | chr9 | 134163620 | Astrocyte 1 | -162710 | 11.67990375 | 0.002 | Intergenic | DEGs, DAGs | -1.22 | 5.850E-67 | NEG | NEG |
| *RXRA* | chr9 | 134370038 | Astrocyte 1 | 43708 | 5.793401092 | 0.001 | intron | DEGs, DAGs | -1.22 | 5.850E-67 | NEG | NEG |
| *RXRA* | chr9 | 134361775 | Astrocyte 1 | 35445 | 6.575046056 | 0.006 | intron | DEGs, DAGs | -1.22 | 5.850E-67 | NEG | NEG |
| *RXRA* | chr9 | 134434494 | Astrocyte 1 | 108164 | 2.966122147 | 0.008 | intron | DEGs, DAGs | -1.22 | 5.850E-67 | NEG | NEG |
| *RXRA* | chr9 | 134354442 | Astrocyte 1 | 28112 | 6.275306825 | 0.024 | intron | DEGs, DAGs | -1.22 | 5.850E-67 | NEG | NEG |
| *RXRA* | chr9 | 134385086 | Astrocyte 1 | 58756 | 2.581741167 | 0.024 | intron | DEGs, DAGs | -1.22 | 5.850E-67 | NEG | NEG |
| *RXRA* | chr9 | 134484264 | Astrocyte 1 | 157934 | 2.353452296 | 0.045 | Intergenic | DEGs, DAGs | -1.22 | 5.850E-67 | NEG | NEG |
| *MXI1* | chr10 | 110212064 | Astrocyte 1 | 2083 | 3.778389575 | 0.047 | intron | DEGs, DAGs | -1.25 | 6.794E-114 | NEG | NEG |

| | | | | | | | | | | | | |
|---|---|---|---|---|---|---|---|---|---|---|---|---|
| *MXI1* | chr10 | 110141878 | Astrocyte 1 | -65477 | 3.310311704 | 0.018 | Intergenic | DEGs, DAGs | -1.25 | 6.794E-114 | NEG | NEG |
| *MXI1* | chr10 | 110212064 | Astrocyte 1 | 4709 | 3.778389575 | 0.047 | intron | DEGs, DAGs | -1.25 | 6.794E-114 | NEG | NEG |
| *FOS* | chr14 | 75276950 | Astrocyte 1 | -1574 | 2.306953157 | 0.014 | promoter-TSS | DEGs, DAGs | -1.55 | 1.822E-60 | NEG | NEG |
| *FOS* | chr14 | 75277563 | Astrocyte 1 | -961 | 3.276642374 | 0.024 | promoter-TSS | DEGs, DAGs | -1.55 | 1.822E-60 | NEG | NEG |
| *FOS* | chr14 | 75278122 | Astrocyte 1 | -402 | 2.09918502 | 0.045 | promoter-TSS | DEGs, DAGs | -1.55 | 1.822E-60 | NEG | NEG |
| *ZBTB7C* | chr18 | 48408812 | Astrocyte 1 | 360 | 3.963677132 | 0.009 | promoter-TSS | DEGs, DAGs | -1.20 | 1.238E-44 | NEG | NEG |
| *ZBTB7C* | chr18 | 48408812 | Astrocyte 1 | -243496 | 3.963677104 | 0.009 | exon; intron | DEGs, DAGs | -1.20 | 1.238E-44 | NEG | NEG |
| *JUNB* | chr19 | 12790638 | Astrocyte 1 | -608 | 2.227239295 | 0.020 | promoter-TSS | DEGs, DAGs | -1.51 | 1.074E-62 | NEG | NEG |
| *JUNB* | chr19 | 12792004 | Astrocyte 1 | 758 | 2.043594319 | 0.029 | promoter-TSS | DEGs, DAGs | -1.51 | 1.074E-62 | NEG | NEG |
| *KLF2* | chr19 | 16272869 | Astrocyte 1 | -51698 | 3.975325519 | 0.003 | Intergenic | DEGs, DAGs | -1.39 | 2.756E-85 | NEG | NEG |
| *KLF2* | chr19 | 16457812 | Astrocyte 1 | 133245 | 6.826173104 | 0.014 | Intergenic | DEGs, DAGs | -1.39 | 2.756E-85 | NEG | NEG |
| *KLF2* | chr19 | 16460392 | Astrocyte 1 | 135825 | 3.231276551 | 0.024 | Intergenic | DEGs, DAGs | -1.39 | 2.756E-85 | NEG | NEG |
| *PURA* | chr5 | 140108994 | Astrocyte 1 | -4879 | 4.453306534 | 0.020 | intron | DEGs, DAGs | 1.25 | 6.955E-61 | NEG | NEG |
| *SREBF1* | chr17 | 17877952 | Astrocyte 1 | -41215 | 6.086119959 | 0.025 | Intergenic | DEGs, DAGs | 1.27 | 7.140E-76 | NEG | NEG |
| *TEAD1* | chr11 | 12941548 | Astrocyte 1 | 267207 | 6.428975042 | 0.013 | exon; intron | DEGs, DAGs | -1.23 | 5.493E-75 | NEG | NEG |
| *TEAD1* | chr11 | 12845625 | Astrocyte 1 | 171284 | 3.75344429 | 0.046 | intron | DEGs, DAGs | -1.23 | 5.493E-75 | NEG | NEG |
| *KLF6* | chr10 | 3538892 | Astrocyte 1 | 246133 | -11.66457283 | 0.044 | Intergenic | DEGs, DAGs | -1.21 | 3.251E-17 | NEG | NEG |
| *FOS* | chr14 | 75296006 | Astrocyte 1 | 17482 | 4.581231794 | 0.016 | Distal enhancer | DEGs, DAGs | -1.55 | 1.822E-60 | GH14J075273; GH14J075293 | eRNA_co-expression; eQTLs |
| *JUNB* | chr19 | 12782256 | Astrocyte 1 | -8990 | 3.961706738 | 0.004 | Distal enhancer | DEGs, DAGs | -1.51 | 1.074E-62 | GH19J012781; GH19J013160; GH19J013148 | eRNA_co-expression; eQTLs, |
| *JUNB* | chr19 | 12782903 | Astrocyte 1 | -8343 | 3.470709101 | 0.035 | Distal enhancer | DEGs, DAGs | -1.51 | 1.074E-62 | GH19J012781; GH19J013160; GH19J013148 | eRNA_co-expression; eQTLs |
| *JUNB* | chr19 | 12796444 | Astrocyte 1 | 5198 | 6.13381936 | 0.011 | Distal enhancer | DEGs, DAGs | -1.51 | 1.074E-62 | GH19J012781; GH19J013160; GH19J013148 | eRNA_co-expression; eQTLs |
| *HIF1A* | chr14 | 61574030 | Astrocyte 1 | -123342 | -5.185814892 | 0.008 | Distal enhancer | DEGs, DAGs | 1.30 | 4.916E-82 | GH14J061465; GH14J061468; GH14J061489; GH14J061510; GH14J061518; GH14J061530; GH14J061551; GH14J061580; GH14J061593; GH14J061599; GH14J061601; GH14J061619; GH14J061641; GH14J061660; GH14J061747; GH14J061717 | C-HiC, eRNA_co-expression; eQTLs |

FC: Fold change; DEGs: Differentially Expressed Genes; DAGs: Differentially Accessible Genes; TSS: Transcription Starting Site; Negative value in Distance to TSS reflects peaks falling upstream of TSS; Negative FC reflects higher accessibility or expression in Afircan local ancestry; NEG: not ELITE enhancers found

**Supplementary Table 5.** *APOE* Local ancestry accessibility analysis

| Gene | FC | FDR | Distance to TSS | Region |
|---|---|---|---|---|
| *AC007191.4* | 5.88 | 0.044 | 10056 | intergenic |
| *APOC1* | 2.23 | 0.025 | 12995 | intergenic |
| *APOC4* | 2.23 | 0.025 | -14996 | intergenic |
| *BCAM* | 4.27 | 0.019 | -12250 | intergenic |
| *BCAM* | 9.08 | 0.024 | -8440 | intergenic |
| *BCL3* | 3.66 | 0.003 | -29841 | intergenic |
| *CBLC* | 4.27 | 0.019 | 18984 | intron |
| *CBLC* | 9.08 | 0.024 | 22794 | exon, intergenic |
| *CEACAM16* | 3.66 | 0.003 | 19555 | intergenic |
| *CKM* | 3.65 | 0.011 | 28118 | intergenic |
| *CKM* | 5.62 | 0.029 | 12819 | intron |
| *CKM* | 6.75 | 0.008 | -3303 | intergenic |
| *ERCC1* | 2.76 | 0.000 | -20885 | intron |
| *ERCC1* | 3.08 | 0.026 | -16451 | intron |
| *FBXO46* | 5.88 | 0.044 | 23579 | intergenic |
| *FOSB* | 2.34 | 0.008 | 17439 | intergenic |
| *FOSB* | 2.76 | 0.000 | -23544 | Distal enhancer |
| *FOSB* | 3.08 | 0.026 | -27978 | Distal enhancer |
| *GIPR* | 4.86 | 0.002 | 13457 | exon, intron |
| *LLNLR-231D4.1* | 4.27 | 0.019 | -6848 | intron |
| *MARK4* | 3.65 | 0.011 | 43601 | intron |
| *MARK4* | 5.62 | 0.029 | 58900 | intergenic |
| *MIR6088* | 2.76 | 0.000 | 7797 | intron |
| *MIR6088* | 3.08 | 0.026 | 3363 | intron |
| *MIR642B* | 4.86 | 0.002 | -6693 | exon |
| *OPA3* | 4.23 | 0.017 | 60969 | intergenic |
| *PPM1N* | 2.34 | 0.008 | -3343 | exon |
| *QPCTL* | 5.88 | 0.044 | 14842 | intergenic |
| *RTN2* | 2.34 | 0.008 | 11627 | exon, intergenic |
| *SNRPD2* | 4.86 | 0.002 | 10632 | intergenic |
| *VASP* | 4.23 | 0.017 | 16457 | intron, exon |
| *VASP* | 4.23 | 0.017 | 16457 | exon |

FC: Fold change; TSS: Transcription Starting Site; Negative value in Distance to TSS reflects peaks falling upstream of TSS

**Supplementary Table 6.** Differentially accessible and expressed genes (DEG-DAG) in astrocyte cluster 1

| Gene | Chromosome | Peak location | Cell cluster | Distance to TSS | FC (snATAC-seq) | FDR (snATAC-seq) | Region | Differentially | FC (snRNA-seq) | FDR (snRNA-seq) | ELITE enhancer (GeneHancer Db) | geneAssociationMethods ELITE enhancer |
|---|---|---|---|---|---|---|---|---|---|---|---|---|
| *CAPZB* | chr1 | 19405759 | Astrocyte 1 | 13742 | 5.47 | 0.003 | intron | DEGs, DAGs | -1.254 | 1.72E-60 | NEG | NEG |
| *KCNN3* | chr1 | 154866993 | Astrocyte 1 | 1076 | 2.97 | 0.004 | promoter-TSS | DEGs, DAGs | -1.256 | 1.59E-33 | NEG | NEG |
| *MACF1* | chr1 | 39204515 | Astrocyte 1 | 519 | 17.26 | 0.008 | promoter-TSS | DEGs, DAGs | 1.266 | 2.76E-75 | NEG | NEG |
| *MACF1* | chr1 | 39090243 | Astrocyte 1 | 6326 | 2.40 | 0.009 | intron | DEGs, DAGs | 1.266 | 2.76E-75 | NEG | NEG |
| *ATP1A2* | chr1 | 160122920 | Astrocyte 1 | 7393 | 5.84 | 0.011 | intron | DEGs, DAGs | 1.507 | 3.27E-138 | NEG | NEG |
| *TXNIP* | chr1 | 145985936 | Astrocyte 1 | 9577 | 3.06 | 0.011 | Intergenic | DEGs, DAGs | -1.238 | 4.28E-40 | NEG | NEG |
| *KCNN3* | chr1 | 154776183 | Astrocyte 1 | 83410 | 3.84 | 0.011 | intron | DEGs, DAGs | -1.256 | 1.59E-33 | NEG | NEG |
| *RNF220* | chr1 | 44423504 | Astrocyte 1 | 18466 | 2.75 | 0.014 | intron | DEGs, DAGs | -1.365 | 1.25E-53 | NEG | NEG |
| *KCNN3* | chr1 | 154751500 | Astrocyte 1 | 108093 | 2.33 | 0.015 | intron | DEGs, DAGs | -1.256 | 1.59E-33 | NEG | NEG |
| *CNTN2* | chr1 | 205061544 | Astrocyte 1 | 18598 | 2.35 | 0.016 | intron | DEGs, DAGs | -1.204 | 5.98E-09 | NEG | NEG |
| *NIPAL3* | chr1 | 24415383 | Astrocyte 1 | -161 | 2.44 | 0.024 | promoter-TSS | DEGs, DAGs | -1.312 | 6.48E-33 | NEG | NEG |
| *MACF1* | chr1 | 39082823 | Astrocyte 1 | -989 | 6.76 | 0.025 | promoter-TSS | DEGs, DAGs | 1.266 | 2.76E-75 | NEG | NEG |
| *CSRP1* | chr1 | 201494211 | Astrocyte 1 | 2112 | 4.37 | 0.026 | intron | DEGs, DAGs | -1.335 | 4.90E-71 | NEG | NEG |
| *PIK3C2B* | chr1 | 204494520 | Astrocyte 1 | -4346 | 2.07 | 0.029 | Intergenic | DEGs, DAGs | -1.203 | 1.30E-27 | NEG | NEG |
| *PIK3C2B* | chr1 | 204473218 | Astrocyte 1 | 16878 | 5.63 | 0.039 | intron | DEGs, DAGs | -1.203 | 1.30E-27 | NEG | NEG |
| *KAZN* | chr1 | 14930548 | Astrocyte 1 | 998 | 10.33 | 0.044 | promoter-TSS | DEGs, DAGs | -1.370 | 3.41E-43 | NEG | NEG |
| *AK5* | chr1 | 77489988 | Astrocyte 1 | -41869 | 2.58 | 0.045 | intron | DEGs, DAGs | -1.296 | 2.34E-26 | NEG | NEG |
| *ACOT11* | chr1 | 54579410 | Astrocyte 1 | 31432 | 2.23 | 0.046 | intron | DEGs, DAGs | -1.244 | 1.85E-35 | NEG | NEG |
| *TTC7A* | chr2 | 46940895 | Astrocyte 1 | -29 | 5.83 | 0.000 | promoter-TSS | DEGs, DAGs | -1.221 | 1.84E-61 | NEG | NEG |
| *GLI2* | chr2 | 120758032 | Astrocyte 1 | -34127 | 4.36 | 0.002 | Intergenic | DEGs, DAGs | -1.211 | 5.59E-53 | NEG | NEG |
| *DPYSL5* | chr2 | 26846287 | Astrocyte 1 | -1564 | 3.91 | 0.004 | promoter-TSS | DEGs, DAGs | -1.289 | 1.46E-24 | NEG | NEG |
| *EFHD1* | chr2 | 232659199 | Astrocyte 1 | -3284 | 3.34 | 0.006 | intron | DEGs, DAGs | -1.315 | 1.40E-66 | NEG | NEG |
| *EFHD1* | chr2 | 232669038 | Astrocyte 1 | 6546 | 10.89 | 0.013 | intron | DEGs, DAGs | -1.315 | 1.40E-66 | NEG | NEG |
| *AGAP1* | chr2 | 235638486 | Astrocyte 1 | -30868 | 12.91 | 0.015 | intron | DEGs, DAGs | -1.270 | 3.22E-66 | NEG | NEG |
| *PRKCE* | chr2 | 45874186 | Astrocyte 1 | -126466 | 4.77 | 0.018 | intron | DEGs, DAGs | -1.219 | 4.54E-49 | NEG | NEG |
| *SFXN5* | chr2 | 73085572 | Astrocyte 1 | -13986 | 3.74 | 0.019 | intron | DEGs, DAGs | -1.236 | 1.99E-25 | NEG | NEG |
| *NRXN1* | chr2 | 50651690 | Astrocyte 1 | 4470 | 2.12 | 0.020 | intron | DEGs, DAGs | 1.383 | 1.48E-83 | NEG | NEG |
| *DNMT3A* | chr2 | 25256938 | Astrocyte 1 | -4877 | 3.25 | 0.035 | intron | DEGs, DAGs | -1.415 | 3.81E-61 | NEG | NEG |
| *IQCA1* | chr2 | 236485530 | Astrocyte 1 | 17917 | 5.21 | 0.045 | intron | DEGs, DAGs | 1.737 | 1.61E-171 | NEG | NEG |
| *AGAP1* | chr2 | 235494677 | Astrocyte 1 | 820 | 3.18 | 0.046 | promoter-TSS | DEGs, DAGs | -1.270 | 3.22E-66 | NEG | NEG |

| | | | | | | | | | | | | |
|---|---|---|---|---|---|---|---|---|---|---|---|---|
| *DYSF* | chr2 | 71672072 | Astrocyte 1 | 205620 | 2.83 | 0.047 | intron | DEGs, DAGs | -1.253 | 6.88E-21 | NEG | NEG |
| *NPAS2* | chr2 | 100818483 | Astrocyte 1 | -1419 | 2.52 | 0.050 | promoter-TSS | DEGs, DAGs | -1.552 | 4.59E-57 | NEG | NEG |
| *IGSF11* | chr3 | 118991743 | Astrocyte 1 | 42735 | 2.55 | 0.003 | intron | DEGs, DAGs | 1.413 | 2.49E-98 | NEG | NEG |
| *MRAS* | chr3 | 138357470 | Astrocyte 1 | 8458 | 2.51 | 0.011 | intron | DEGs, DAGs | -1.451 | 3.05E-64 | NEG | NEG |
| *IGSF11* | chr3 | 119034692 | Astrocyte 1 | -54 | 2.02 | 0.012 | promoter-TSS | DEGs, DAGs | 1.413 | 2.49E-98 | NEG | NEG |
| *TMEM108* | chr3 | 133041158 | Astrocyte 1 | 3006 | 2.48 | 0.014 | intron | DEGs, DAGs | 1.294 | 1.26E-60 | NEG | NEG |
| *CACNA2D3* | chr3 | 54146596 | Astrocyte 1 | 23385 | 3.17 | 0.027 | intron | DEGs, DAGs | 1.523 | 8.67E-65 | NEG | NEG |
| *MRAS* | chr3 | 138359412 | Astrocyte 1 | 10400 | 2.23 | 0.028 | intron | DEGs, DAGs | -1.451 | 3.05E-64 | NEG | NEG |
| *LRIG1* | chr3 | 66481842 | Astrocyte 1 | 18840 | 2.20 | 0.033 | intron | DEGs, DAGs | -1.224 | 1.39E-118 | NEG | NEG |
| *KLF15* | chr3 | 126341652 | Astrocyte 1 | 15540 | 2.70 | 0.035 | TTS | DEGs, DAGs | -1.209 | 1.08E-55 | NEG | NEG |
| *FGFR3* | chr4 | 1797770 | Astrocyte 1 | 4124 | 2.07 | 0.002 | intron | DEGs, DAGs | 1.330 | 1.07E-88 | NEG | NEG |
| *APBB2* | chr4 | 40845172 | Astrocyte 1 | 11733 | 2.07 | 0.015 | intron | DEGs, DAGs | -1.215 | 4.72E-59 | NEG | NEG |
| *FGFR3* | chr4 | 1808860 | Astrocyte 1 | 15214 | 3.51 | 0.036 | TTS | DEGs, DAGs | 1.330 | 1.07E-88 | NEG | NEG |
| *CPE* | chr4 | 165361033 | Astrocyte 1 | -17659 | 2.27 | 0.045 | Intergenic | DEGs, DAGs | 1.290 | 2.26E-55 | NEG | NEG |
| *FGFR3* | chr4 | 1788491 | Astrocyte 1 | -4552 | 6.51 | 0.045 | Intergenic | DEGs, DAGs | 1.330 | 1.07E-88 | NEG | NEG |
| *ERGIC1* | chr5 | 172867295 | Astrocyte 1 | 33134 | 4.48 | 0.001 | intron | DEGs, DAGs | -1.227 | 6.40E-52 | NEG | NEG |
| *FGF1* | chr5 | 142678528 | Astrocyte 1 | 6894 | 2.09 | 0.006 | intron | DEGs, DAGs | -1.299 | 1.17E-52 | NEG | NEG |
| *WWC1* | chr5 | 168334631 | Astrocyte 1 | 42731 | 3.21 | 0.009 | intron | DEGs, DAGs | -1.410 | 5.06E-70 | NEG | NEG |
| *SH3PXD2B* | chr5 | 172452765 | Astrocyte 1 | 1343 | 5.01 | 0.013 | promoter-TSS | DEGs, DAGs | -1.286 | 4.62E-47 | NEG | NEG |
| *FGF1* | chr5 | 142611006 | Astrocyte 1 | 10086 | 2.21 | 0.018 | intron | DEGs, DAGs | -1.299 | 1.17E-52 | NEG | NEG |
| *SH3PXD2B* | chr5 | 172497022 | Astrocyte 1 | -42749 | 2.10 | 0.046 | Intergenic | DEGs, DAGs | -1.286 | 4.62E-47 | NEG | NEG |
| *SLC22A23* | chr6 | 3456974 | Astrocyte 1 | -202 | 2.34 | 0.005 | promoter-TSS | DEGs, DAGs | -1.391 | 1.05E-91 | NEG | NEG |
| *DST* | chr6 | 56552089 | Astrocyte 1 | 90521 | 22.51 | 0.043 | exon | DEGs, DAGs | 1.296 | 1.70E-82 | NEG | NEG |
| *KLHL32* | chr6 | 97048443 | Astrocyte 1 | 38494 | -7.01 | 0.044 | intron | DEGs, DAGs | -1.198 | 8.80E-79 | NEG | NEG |
| *VEGFA* | chr6 | 43770739 | Astrocyte 1 | -94 | 2.39 | 0.048 | promoter-TSS | DEGs, DAGs | 1.284 | 1.68E-130 | NEG | NEG |
| *HSPB1* | chr7 | 76314257 | Astrocyte 1 | 10930 | 2.23 | 0.000 | Intergenic | DEGs, DAGs | 1.422 | 1.56E-31 | NEG | NEG |
| *COX19* | chr7 | 975366 | Astrocyte 1 | -17 | 3.84 | 0.002 | promoter-TSS | DEGs, DAGs | -1.302 | 2.99E-47 | NEG | NEG |
| *GTF2IRD1* | chr7 | 74430833 | Astrocyte 1 | -22707 | 2.76 | 0.005 | Intergenic | DEGs, DAGs | -1.348 | 5.20E-58 | NEG | NEG |
| *CUX1* | chr7 | 101733828 | Astrocyte 1 | -81826 | 3.04 | 0.010 | Intergenic | DEGs, DAGs | -1.271 | 8.56E-65 | NEG | NEG |
| *GTF2IRD1* | chr7 | 74542465 | Astrocyte 1 | 36723 | 9.78 | 0.012 | intron | DEGs, DAGs | -1.348 | 5.20E-58 | NEG | NEG |
| *HSPB1* | chr7 | 76309982 | Astrocyte 1 | 6655 | 3.13 | 0.026 | Intergenic | DEGs, DAGs | 1.422 | 1.56E-31 | NEG | NEG |
| *HSPB1* | chr7 | 76305500 | Astrocyte 1 | 2173 | 3.73 | 0.031 | Intergenic | DEGs, DAGs | 1.422 | 1.56E-31 | NEG | NEG |
| *CLIP2* | chr7 | 74379882 | Astrocyte 1 | 62585 | 2.34 | 0.031 | intron | DEGs, DAGs | -1.528 | 1.83E-58 | NEG | NEG |
| *ADRA1A* | chr8 | 26863491 | Astrocyte 1 | 1235 | 6.36 | 0.033 | promoter-TSS | DEGs, DAGs | 1.276 | 9.55E-62 | NEG | NEG |

| | | | | | | | | | | | | |
|---|---|---|---|---|---|---|---|---|---|---|---|---|
| *SCARA3* | chr8 | 27633557 | Astrocyte 1 | -61 | 2.51 | 0.043 | promoter-TSS | DEGs, DAGs | -1.510 | 1.88E-117 | NEG | NEG |
| *LHX2* | chr9 | 124011402 | Astrocyte 1 | 42 | 2.39 | 0.000 | promoter-TSS | DEGs, DAGs | -1.389 | 7.39E-66 | NEG | NEG |
| *NEK6* | chr9 | 124295915 | Astrocyte 1 | 3224 | 3.95 | 0.000 | intron | DEGs, DAGs | -1.406 | 2.80E-71 | NEG | NEG |
| *NEK6* | chr9 | 124291677 | Astrocyte 1 | -44 | 4.41 | 0.002 | promoter-TSS | DEGs, DAGs | -1.406 | 2.80E-71 | NEG | NEG |
| *LCNL1* | chr9 | 136980874 | Astrocyte 1 | -1869 | 4.50 | 0.004 | promoter-TSS | DEGs, DAGs | 1.273 | 3.08E-122 | NEG | NEG |
| *RXRA* | chr9 | 134348401 | Astrocyte 1 | 22071 | 3.62 | 0.006 | intron | DEGs, DAGs | -1.215 | 5.85E-67 | NEG | NEG |
| *RXRA* | chr9 | 134344653 | Astrocyte 1 | 18323 | 2.22 | 0.007 | intron | DEGs, DAGs | -1.215 | 5.85E-67 | NEG | NEG |
| *NEK6* | chr9 | 124188229 | Astrocyte 1 | -69127 | 5.03 | 0.010 | Intergenic | DEGs, DAGs | -1.406 | 2.80E-71 | NEG | NEG |
| *GSN* | chr9 | 121317720 | Astrocyte 1 | 18177 | 3.38 | 0.011 | intron | DEGs, DAGs | -1.420 | 3.57E-112 | NEG | NEG |
| *LCNL1* | chr9 | 136981583 | Astrocyte 1 | -1160 | 6.08 | 0.011 | promoter-TSS | DEGs, DAGs | 1.273 | 3.08E-122 | NEG | NEG |
| *RXRA* | chr9 | 134331104 | Astrocyte 1 | 4774 | 2.19 | 0.016 | intron | DEGs, DAGs | -1.215 | 5.85E-67 | NEG | NEG |
| *GSN* | chr9 | 121289239 | Astrocyte 1 | 2897 | 3.43 | 0.017 | intron | DEGs, DAGs | -1.420 | 3.57E-112 | NEG | NEG |
| *ZER1* | chr9 | 128771825 | Astrocyte 1 | -182 | 2.11 | 0.019 | promoter-TSS | DEGs, DAGs | -1.398 | 1.74E-68 | NEG | NEG |
| *RXRA* | chr9 | 134338784 | Astrocyte 1 | 12454 | 3.40 | 0.019 | intron | DEGs, DAGs | -1.215 | 5.85E-67 | NEG | NEG |
| *NEK6* | chr9 | 124343043 | Astrocyte 1 | 50352 | 3.04 | 0.022 | intron | DEGs, DAGs | -1.406 | 2.80E-71 | NEG | NEG |
| *RXRA* | chr9 | 134326812 | Astrocyte 1 | 482 | 7.82 | 0.023 | promoter-TSS | DEGs, DAGs | -1.215 | 5.85E-67 | NEG | NEG |
| *LCNL1* | chr9 | 136982235 | Astrocyte 1 | -508 | 2.30 | 0.026 | promoter-TSS | DEGs, DAGs | 1.273 | 3.08E-122 | NEG | NEG |
| *LCNL1* | chr9 | 136980104 | Astrocyte 1 | -2639 | 3.47 | 0.031 | intron | DEGs, DAGs | 1.273 | 3.08E-122 | NEG | NEG |
| *RXRA* | chr9 | 134316329 | Astrocyte 1 | -10001 | 3.43 | 0.039 | Intergenic | DEGs, DAGs | -1.215 | 5.85E-67 | NEG | NEG |
| *LHX2* | chr9 | 124013712 | Astrocyte 1 | 2352 | 4.45 | 0.042 | exon | DEGs, DAGs | -1.389 | 7.39E-66 | NEG | NEG |
| *RXRA* | chr9 | 134340714 | Astrocyte 1 | 14384 | 15.35 | 0.043 | intron | DEGs, DAGs | -1.215 | 5.85E-67 | NEG | NEG |
| *NEK6* | chr9 | 124259376 | Astrocyte 1 | 1375 | 2.27 | 0.043 | promoter-TSS | DEGs, DAGs | -1.406 | 2.80E-71 | NEG | NEG |
| *RGS3* | chr9 | 113566609 | Astrocyte 1 | 1905 | 2.59 | 0.044 | promoter-TSS | DEGs, DAGs | -1.240 | 4.11E-53 | NEG | NEG |
| *COL27A1* | chr9 | 114153635 | Astrocyte 1 | -1675 | 4.44 | 0.046 | promoter-TSS | DEGs, DAGs | -1.357 | 1.23E-46 | NEG | NEG |
| *ABCA2* | chr9 | 137028017 | Astrocyte 1 | 21 | 2.15 | 0.047 | promoter-TSS | DEGs, DAGs | -1.319 | 1.32E-43 | NEG | NEG |
| *LHX2* | chr9 | 124018110 | Astrocyte 1 | 6750 | 2.50 | 0.049 | intron | DEGs, DAGs | -1.389 | 7.39E-66 | NEG | NEG |
| *ZMIZ1* | chr10 | 79093984 | Astrocyte 1 | 25199 | 5.44 | 0.004 | intron | DEGs, DAGs | -1.210 | 7.98E-52 | NEG | NEG |
| *ZMIZ1* | chr10 | 79282683 | Astrocyte 1 | -2520 | 3.86 | 0.004 | intron | DEGs, DAGs | -1.210 | 7.98E-52 | NEG | NEG |
| *CAMK2G* | chr10 | 73874447 | Astrocyte 1 | -106 | 2.23 | 0.006 | promoter-TSS | DEGs, DAGs | -1.448 | 2.18E-82 | NEG | NEG |
| *VSIR* | chr10 | 71757267 | Astrocyte 1 | 15981 | 2.26 | 0.013 | intron | DEGs, DAGs | -1.236 | 2.17E-39 | NEG | NEG |
| *ZMIZ1* | chr10 | 79273507 | Astrocyte 1 | -11696 | 5.28 | 0.014 | intron | DEGs, DAGs | -1.210 | 7.98E-52 | NEG | NEG |
| *ZMIZ1* | chr10 | 79195360 | Astrocyte 1 | -5985 | 3.00 | 0.018 | intron | DEGs, DAGs | -1.210 | 7.98E-52 | NEG | NEG |
| *ZMIZ1* | chr10 | 79240241 | Astrocyte 1 | 38896 | 4.19 | 0.019 | intron | DEGs, DAGs | -1.210 | 7.98E-52 | NEG | NEG |

| | | | | | | | | | | | | |
|---|---|---|---|---|---|---|---|---|---|---|---|---|
| *EMX2OS* | chr10 | 117551660 | Astrocyte 1 | -6842 | 3.01 | 0.024 | Intergenic | DEGs, DAGs | -1.262 | 1.11E-61 | NEG | NEG |
| *SORBS1* | chr10 | 95435294 | Astrocyte 1 | 5591 | 4.39 | 0.024 | intron | DEGs, DAGs | -1.281 | 2.99E-44 | NEG | NEG |
| *HK1* | chr10 | 69337547 | Astrocyte 1 | 18953 | 51.65 | 0.028 | intron | DEGs, DAGs | -1.215 | 3.79E-51 | NEG | NEG |
| *CHST3* | chr10 | 71975516 | Astrocyte 1 | 11401 | 2.22 | 0.028 | intron | DEGs, DAGs | -1.231 | 3.60E-28 | NEG | NEG |
| *TACC2* | chr10 | 122192068 | Astrocyte 1 | 28474 | 2.98 | 0.028 | intron | DEGs, DAGs | -1.234 | 4.39E-29 | NEG | NEG |
| *CHST3* | chr10 | 71957526 | Astrocyte 1 | -6589 | 6.10 | 0.028 | Intergenic | DEGs, DAGs | -1.231 | 3.60E-28 | NEG | NEG |
| *ZMIZ1* | chr10 | 79248500 | Astrocyte 1 | -36703 | 2.21 | 0.031 | intron | DEGs, DAGs | -1.210 | 7.98E-52 | NEG | NEG |
| *CDH23* | chr10 | 71473854 | Astrocyte 1 | 34272 | 3.68 | 0.032 | intron | DEGs, DAGs | -1.419 | 4.96E-48 | NEG | NEG |
| *ZMIZ1* | chr10 | 79165758 | Astrocyte 1 | -35587 | 2.81 | 0.035 | intron | DEGs, DAGs | -1.210 | 7.98E-52 | NEG | NEG |
| *ZMIZ1* | chr10 | 79254756 | Astrocyte 1 | -30447 | 3.16 | 0.040 | intron | DEGs, DAGs | -1.210 | 7.98E-52 | NEG | NEG |
| *CHST3* | chr10 | 71999269 | Astrocyte 1 | 35154 | 2.19 | 0.041 | intron | DEGs, DAGs | -1.231 | 3.60E-28 | NEG | NEG |
| *CHST3* | chr10 | 71965050 | Astrocyte 1 | 935 | 2.74 | 0.045 | promoter-TSS | DEGs, DAGs | -1.231 | 3.60E-28 | NEG | NEG |
| *SORBS1* | chr10 | 95457525 | Astrocyte 1 | -16613 | 2.61 | 0.045 | intron | DEGs, DAGs | -1.281 | 2.99E-44 | NEG | NEG |
| *FGFR2* | chr10 | 121539701 | Astrocyte 1 | -8637 | 6.31 | 0.047 | intron | DEGs, DAGs | -1.250 | 4.85E-118 | NEG | NEG |
| *MXI1* | chr10 | 110212064 | Astrocyte 1 | 2083 | 3.78 | 0.047 | intron | DEGs, DAGs | -1.250 | 6.79E-114 | NEG | NEG |
| *B4GALNT4* | chr11 | 369176 | Astrocyte 1 | -378 | 4.91 | 0.002 | promoter-TSS | DEGs, DAGs | 1.316 | 7.27E-80 | NEG | NEG |
| *SLC1A2* | chr11 | 35362258 | Astrocyte 1 | -2380 | 11.26 | 0.009 | intron | DEGs, DAGs | 1.587 | 9.94E-114 | NEG | NEG |
| *SLC3A2* | chr11 | 62883502 | Astrocyte 1 | 2027 | 2.45 | 0.011 | intron | DEGs, DAGs | 1.394 | 1.07E-83 | NEG | NEG |
| *NTM* | chr11 | 131373942 | Astrocyte 1 | 3714 | 4.61 | 0.015 | intron | DEGs, DAGs | 1.336 | 7.51E-57 | NEG | NEG |
| *ZBTB16* | chr11 | 114072097 | Astrocyte 1 | 11840 | 2.63 | 0.018 | intron | DEGs, DAGs | -1.313 | 1.27E-83 | NEG | NEG |
| *SLC1A2* | chr11 | 35346145 | Astrocyte 1 | 13655 | 2.92 | 0.026 | intron | DEGs, DAGs | 1.587 | 9.94E-114 | NEG | NEG |
| *ZBTB16* | chr11 | 114064017 | Astrocyte 1 | 3760 | 14.24 | 0.038 | exon | DEGs, DAGs | -1.313 | 1.27E-83 | NEG | NEG |
| *B4GALNT4* | chr11 | 379231 | Astrocyte 1 | 9677 | 18.52 | 0.043 | exon | DEGs, DAGs | 1.316 | 7.27E-80 | NEG | NEG |
| *TMEM132B* | chr12 | 125330854 | Astrocyte 1 | 4488 | 8.49 | 0.000 | intron | DEGs, DAGs | -1.209 | 2.02E-22 | NEG | NEG |
| *TUBA1A* | chr12 | 49188882 | Astrocyte 1 | -66 | 6.44 | 0.000 | promoter-TSS | DEGs, DAGs | -1.271 | 1.05E-52 | NEG | NEG |
| *PITPNM2* | chr12 | 123071756 | Astrocyte 1 | 9327 | 10.51 | 0.002 | intron | DEGs, DAGs | -1.207 | 6.27E-56 | NEG | NEG |
| *PITPNM2* | chr12 | 123043412 | Astrocyte 1 | -9008 | 3.83 | 0.016 | intron | DEGs, DAGs | -1.207 | 6.27E-56 | NEG | NEG |
| *PITPNM2* | chr12 | 123059510 | Astrocyte 1 | 21573 | 4.30 | 0.019 | intron | DEGs, DAGs | -1.207 | 6.27E-56 | NEG | NEG |
| *TMEM132B* | chr12 | 125407786 | Astrocyte 1 | 81420 | 7.08 | 0.025 | intron | DEGs, DAGs | -1.209 | 2.02E-22 | NEG | NEG |
| *PITPNM2* | chr12 | 123052013 | Astrocyte 1 | -17609 | 10.77 | 0.043 | intron | DEGs, DAGs | -1.207 | 6.27E-56 | NEG | NEG |
| *TMEM132B* | chr12 | 125432713 | Astrocyte 1 | 106347 | 160.82 | 0.043 | intron | DEGs, DAGs | -1.209 | 2.02E-22 | NEG | NEG |
| *CAB39L* | chr13 | 49303565 | Astrocyte 1 | 47312 | 3.17 | 0.025 | Intergenic | DEGs, DAGs | -1.468 | 1.22E-135 | NEG | NEG |
| *DCLK1* | chr13 | 36074555 | Astrocyte 1 | 56501 | 4.85 | 0.044 | intron | DEGs, DAGs | -1.201 | 1.79E-32 | NEG | NEG |
| *FOS* | chr14 | 75276950 | Astrocyte 1 | -1574 | 2.31 | 0.014 | promoter-TSS | DEGs, DAGs | -1.547 | 1.82E-60 | NEG | NEG |

| | | | | | | | | | | | | |
|---|---|---|---|---|---|---|---|---|---|---|---|---|
| *FOS* | chr14 | 75277563 | Astrocyte 1 | -961 | 3.28 | 0.024 | promoter-TSS | DEGs, DAGs | -1.547 | 1.82E-60 | NEG | NEG |
| *FOS* | chr14 | 75278122 | Astrocyte 1 | -402 | 2.10 | 0.045 | promoter-TSS | DEGs, DAGs | -1.547 | 1.82E-60 | NEG | NEG |
| *ACSBG1* | chr15 | 78203184 | Astrocyte 1 | 31273 | 2.72 | 0.006 | intron | DEGs, DAGs | -1.241 | 8.93E-79 | NEG | NEG |
| *ACSBG1* | chr15 | 78205751 | Astrocyte 1 | 28706 | 3.69 | 0.046 | intron | DEGs, DAGs | -1.241 | 8.93E-79 | NEG | NEG |
| *ABCA3* | chr16 | 2339875 | Astrocyte 1 | 610 | 4.37 | 0.008 | promoter-TSS | DEGs, DAGs | -1.274 | 1.11E-56 | NEG | NEG |
| *ABCA3* | chr16 | 2340524 | Astrocyte 1 | -28 | 2.27 | 0.012 | promoter-TSS | DEGs, DAGs | -1.274 | 1.11E-56 | NEG | NEG |
| *MT1G* | chr16 | 56667890 | Astrocyte 1 | -75 | 2.33 | 0.014 | promoter-TSS | DEGs, DAGs | -1.417 | 5.14E-36 | NEG | NEG |
| *GSE1* | chr16 | 85628016 | Astrocyte 1 | 14914 | 4.89 | 0.015 | intron | DEGs, DAGs | -1.237 | 3.56E-37 | NEG | NEG |
| *GSE1* | chr16 | 85602009 | Astrocyte 1 | -9150 | 2.86 | 0.027 | Intergenic | DEGs, DAGs | -1.237 | 3.56E-37 | NEG | NEG |
| *ADGRG1* | chr16 | 57647519 | Astrocyte 1 | 8251 | 8.76 | 0.039 | intron | DEGs, DAGs | 1.338 | 1.56E-100 | NEG | NEG |
| *RGS9* | chr17 | 65100567 | Astrocyte 1 | -169 | 7.10 | 0.000 | promoter-TSS | DEGs, DAGs | -1.199 | 1.97E-12 | NEG | NEG |
| *NXN* | chr17 | 979606 | Astrocyte 1 | -86 | 3.00 | 0.000 | promoter-TSS | DEGs, DAGs | -1.299 | 3.52E-32 | NEG | NEG |
| *TAF15* | chr17 | 35809172 | Astrocyte 1 | -33 | 2.89 | 0.002 | promoter-TSS | DEGs, DAGs | -1.245 | 6.12E-59 | NEG | NEG |
| *SLC43A2* | chr17 | 1629928 | Astrocyte 1 | -1342 | 7.24 | 0.002 | promoter-TSS | DEGs, DAGs | -1.326 | 7.93E-20 | NEG | NEG |
| *ATP1B2* | chr17 | 7650630 | Astrocyte 1 | -232 | 2.16 | 0.003 | promoter-TSS | DEGs, DAGs | 1.668 | 8.62E-133 | NEG | NEG |
| *KANSL1* | chr17 | 46193231 | Astrocyte 1 | -81 | 3.37 | 0.003 | promoter-TSS | DEGs, DAGs | 1.265 | 8.43E-68 | NEG | NEG |
| *LINC00511* | chr17 | 72476453 | Astrocyte 1 | -45544 | 2.11 | 0.004 | intron | DEGs, DAGs | -1.229 | 1.58E-41 | NEG | NEG |
| *ATP1B2* | chr17 | 7654017 | Astrocyte 1 | 3155 | 31.74 | 0.005 | intron | DEGs, DAGs | 1.668 | 8.62E-133 | NEG | NEG |
| *GRIN2C* | chr17 | 74833954 | Astrocyte 1 | 25671 | 2.90 | 0.006 | intron | DEGs, DAGs | -1.220 | 1.95E-92 | NEG | NEG |
| *LINC00511* | chr17 | 72502000 | Astrocyte 1 | -71091 | 2.23 | 0.007 | intron | DEGs, DAGs | -1.229 | 1.58E-41 | NEG | NEG |
| *GRIN2C* | chr17 | 74827045 | Astrocyte 1 | 32580 | 10.58 | 0.008 | intron | DEGs, DAGs | -1.220 | 1.95E-92 | NEG | NEG |
| *GFAP* | chr17 | 44916672 | Astrocyte 1 | -383 | 2.03 | 0.008 | promoter-TSS | DEGs, DAGs | -1.218 | 3.89E-63 | NEG | NEG |
| *GRIN2C* | chr17 | 74843349 | Astrocyte 1 | 16276 | 12.91 | 0.010 | TTS | DEGs, DAGs | -1.220 | 1.95E-92 | NEG | NEG |
| *GRIN2C* | chr17 | 74827671 | Astrocyte 1 | 31954 | 3.05 | 0.011 | intron | DEGs, DAGs | -1.220 | 1.95E-92 | NEG | NEG |
| *GFAP* | chr17 | 44919922 | Astrocyte 1 | -3633 | 3.28 | 0.014 | Intergenic | DEGs, DAGs | -1.218 | 3.89E-63 | NEG | NEG |
| *LINC00511* | chr17 | 72574046 | Astrocyte 1 | 18044 | 7.36 | 0.019 | intron | DEGs, DAGs | -1.229 | 1.58E-41 | NEG | NEG |
| *NXN* | chr17 | 979090 | Astrocyte 1 | 430 | 5.98 | 0.019 | promoter-TSS | DEGs, DAGs | -1.299 | 3.52E-32 | NEG | NEG |
| *LINC00511* | chr17 | 72592555 | Astrocyte 1 | -1 | 2.06 | 0.020 | promoter-TSS | DEGs, DAGs | -1.229 | 1.58E-41 | NEG | NEG |
| *KANSL1* | chr17 | 46192162 | Astrocyte 1 | 388 | 2.95 | 0.020 | promoter-TSS | DEGs, DAGs | 1.265 | 8.43E-68 | NEG | NEG |
| *H3F3B* | chr17 | 75779730 | Astrocyte 1 | -200 | 2.47 | 0.022 | promoter-TSS | DEGs, DAGs | -1.209 | 2.25E-49 | NEG | NEG |
| *GRIN2C* | chr17 | 74858730 | Astrocyte 1 | 895 | 3.58 | 0.024 | promoter-TSS | DEGs, DAGs | -1.220 | 1.95E-92 | NEG | NEG |
| *TOM1L2* | chr17 | 17877952 | Astrocyte 1 | -6138 | 6.09 | 0.025 | intron | DEGs, DAGs | -1.408 | 5.09E-74 | NEG | NEG |
| *GRIN2C* | chr17 | 74834761 | Astrocyte 1 | 24864 | 2.88 | 0.026 | intron | DEGs, DAGs | -1.220 | 1.95E-92 | NEG | NEG |

| | | | | | | | | | | | | |
|---|---|---|---|---|---|---|---|---|---|---|---|---|
| *MINK1* | chr17 | 4833053 | Astrocyte 1 | -85 | 2.20 | 0.026 | promoter-TSS | DEGs, DAGs | -1.273 | 2.25E-54 | NEG | NEG |
| *GRIN2C* | chr17 | 74822883 | Astrocyte 1 | 36742 | 2.07 | 0.026 | intron | DEGs, DAGs | -1.220 | 1.95E-92 | NEG | NEG |
| *STARD3* | chr17 | 39636785 | Astrocyte 1 | -30 | 2.66 | 0.026 | promoter-TSS | DEGs, DAGs | -1.204 | 5.71E-37 | NEG | NEG |
| *LINC00511* | chr17 | 72550325 | Astrocyte 1 | 41765 | 16.72 | 0.028 | intron | DEGs, DAGs | -1.229 | 1.58E-41 | NEG | NEG |
| *SLC39A11* | chr17 | 73088450 | Astrocyte 1 | 3981 | 20.35 | 0.031 | exon | DEGs, DAGs | -1.460 | 1.51E-50 | NEG | NEG |
| *AKAP1* | chr17 | 57092633 | Astrocyte 1 | -2961 | 2.02 | 0.032 | exon | DEGs, DAGs | 1.258 | 1.88E-59 | NEG | NEG |
| *LINC00511* | chr17 | 72416246 | Astrocyte 1 | 6530 | 2.01 | 0.034 | intron | DEGs, DAGs | -1.229 | 1.58E-41 | NEG | NEG |
| *LINC00511* | chr17 | 72586997 | Astrocyte 1 | 5093 | 11.82 | 0.038 | intron | DEGs, DAGs | -1.229 | 1.58E-41 | NEG | NEG |
| *SLC43A2* | chr17 | 1628729 | Astrocyte 1 | -143 | 2.50 | 0.043 | promoter-TSS | DEGs, DAGs | -1.326 | 7.93E-20 | NEG | NEG |
| *LINC00511* | chr17 | 72415506 | Astrocyte 1 | 7270 | 3.61 | 0.044 | intron | DEGs, DAGs | -1.229 | 1.58E-41 | NEG | NEG |
| *GRIN2C* | chr17 | 74828196 | Astrocyte 1 | 31429 | 4.82 | 0.044 | intron | DEGs, DAGs | -1.220 | 1.95E-92 | NEG | NEG |
| *PITPNA* | chr17 | 1561744 | Astrocyte 1 | 797 | 4.37 | 0.045 | promoter-TSS | DEGs, DAGs | -1.295 | 7.86E-38 | NEG | NEG |
| *ZBTB7C* | chr18 | 48408812 | Astrocyte 1 | 360 | 3.96 | 0.009 | promoter-TSS | DEGs, DAGs | -1.205 | 1.24E-44 | NEG | NEG |
| *HIF3A* | chr19 | 46296730 | Astrocyte 1 | -66 | 2.62 | 0.000 | promoter-TSS | DEGs, DAGs | 1.412 | 6.22E-116 | NEG | NEG |
| *FTL* | chr19 | 48965056 | Astrocyte 1 | 5 | 2.69 | 0.000 | promoter-TSS | DEGs, DAGs | 1.463 | 2.53E-34 | NEG | NEG |
| *SAFB2* | chr19 | 5621198 | Astrocyte 1 | 1532 | 8.58 | 0.000 | promoter-TSS | DEGs, DAGs | 1.290 | 7.32E-88 | NEG | NEG |
| *APOE* | chr19 | 44905485 | Astrocyte 1 | -19 | 2.57 | 0.001 | promoter-TSS | DEGs, DAGs | 1.556 | 1.24E-129 | NEG | NEG |
| *EML2* | chr19 | 45645399 | Astrocyte 1 | -20 | 3.28 | 0.002 | promoter-TSS | DEGs, DAGs | -1.202 | 1.80E-20 | NEG | NEG |
| *SLC25A42* | chr19 | 19063547 | Astrocyte 1 | -202 | 2.26 | 0.004 | promoter-TSS | DEGs, DAGs | -1.219 | 2.47E-23 | NEG | NEG |
| *DNM2* | chr19 | 10717210 | Astrocyte 1 | -619 | 2.67 | 0.004 | promoter-TSS | DEGs, DAGs | -1.304 | 3.49E-59 | NEG | NEG |
| *SIPA1L3* | chr19 | 37932209 | Astrocyte 1 | 25231 | 6.15 | 0.004 | intron | DEGs, DAGs | -1.326 | 9.22E-75 | NEG | NEG |
| *NWD1* | chr19 | 16719558 | Astrocyte 1 | -172 | 2.75 | 0.005 | promoter-TSS | DEGs, DAGs | -1.307 | 3.93E-133 | NEG | NEG |
| *ADAMTS10* | chr19 | 8609337 | Astrocyte 1 | 1113 | 3.58 | 0.006 | promoter-TSS | DEGs, DAGs | -1.436 | 1.77E-39 | NEG | NEG |
| *CATSPERG* | chr19 | 38335510 | Astrocyte 1 | -15 | 2.41 | 0.007 | promoter-TSS | DEGs, DAGs | -1.204 | 6.46E-28 | NEG | NEG |
| *RFX2* | chr19 | 6056646 | Astrocyte 1 | -7664 | 4.05 | 0.008 | intron | DEGs, DAGs | -1.328 | 3.49E-49 | NEG | NEG |
| *GADD45B* | chr19 | 2474479 | Astrocyte 1 | -1398 | 3.40 | 0.010 | promoter-TSS | DEGs, DAGs | -1.226 | 1.00E-23 | NEG | NEG |
| *MARK4* | chr19 | 45294609 | Astrocyte 1 | 504 | 3.65 | 0.011 | promoter-TSS | DEGs, DAGs | -1.366 | 3.18E-67 | NEG | NEG |
| *GADD45B* | chr19 | 2488825 | Astrocyte 1 | 12927 | 2.23 | 0.011 | Intergenic | DEGs, DAGs | -1.226 | 1.00E-23 | NEG | NEG |
| *CATSPERG* | chr19 | 38336773 | Astrocyte 1 | 1248 | 3.10 | 0.012 | promoter-TSS | DEGs, DAGs | -1.204 | 6.46E-28 | NEG | NEG |
| *SPACA6* | chr19 | 51702232 | Astrocyte 1 | 8986 | 8.47 | 0.019 | intron | DEGs, DAGs | -1.326 | 1.98E-81 | NEG | NEG |
| *JUNB* | chr19 | 12790638 | Astrocyte 1 | -608 | 2.23 | 0.020 | promoter-TSS | DEGs, DAGs | -1.512 | 1.07E-62 | NEG | NEG |
| *APOE* | chr19 | 44903514 | Astrocyte 1 | -1990 | 4.52 | 0.022 | promoter-TSS | DEGs, DAGs | 1.556 | 1.24E-129 | NEG | NEG |
| *JUNB* | chr19 | 12792004 | Astrocyte 1 | 758 | 2.04 | 0.029 | promoter-TSS | DEGs, DAGs | -1.512 | 1.07E-62 | NEG | NEG |

| | | | | | | | | | | | | |
|---|---|---|---|---|---|---|---|---|---|---|---|---|
| *EML2* | chr19 | 45644874 | Astrocyte 1 | 505 | 7.63 | 0.032 | promoter-TSS | DEGs, DAGs | -1.202 | 1.80E-20 | NEG | NEG |
| *FTL* | chr19 | 48970850 | Astrocyte 1 | 5799 | 5.22 | 0.033 | intron | DEGs, DAGs | 1.463 | 2.53E-34 | NEG | NEG |
| *C3* | chr19 | 6719330 | Astrocyte 1 | 1102 | 2.58 | 0.035 | promoter-TSS | DEGs, DAGs | -1.224 | 4.61E-03 | NEG | NEG |
| *INSR* | chr19 | 7197103 | Astrocyte 1 | 96578 | 2.70 | 0.035 | intron | DEGs, DAGs | -1.288 | 9.60E-66 | NEG | NEG |
| *SLC27A1* | chr19 | 17469284 | Astrocyte 1 | -910 | 2.05 | 0.041 | promoter-TSS | DEGs, DAGs | -1.354 | 4.14E-70 | NEG | NEG |
| *HIF3A* | chr19 | 46297672 | Astrocyte 1 | -468 | 4.21 | 0.044 | promoter-TSS | DEGs, DAGs | 1.412 | 6.22E-116 | NEG | NEG |
| *SAFB2* | chr19 | 5622531 | Astrocyte 1 | 199 | 2.44 | 0.046 | promoter-TSS | DEGs, DAGs | 1.290 | 7.32E-88 | NEG | NEG |
| *EYA2* | chr20 | 46894534 | Astrocyte 1 | -86 | 2.45 | 0.008 | promoter-TSS | DEGs, DAGs | -1.296 | 1.28E-29 | NEG | NEG |
| *EYA2* | chr20 | 46895646 | Astrocyte 1 | 1026 | 2.63 | 0.035 | promoter-TSS | DEGs, DAGs | -1.296 | 1.28E-29 | NEG | NEG |
| *RIN2* | chr20 | 19885992 | Astrocyte 1 | -279 | 2.58 | 0.044 | promoter-TSS | DEGs, DAGs | -1.385 | 8.01E-127 | NEG | NEG |
| *HSCB* | chr22 | 28741710 | Astrocyte 1 | -71 | 2.77 | 0.000 | promoter-TSS | DEGs, DAGs | -1.331 | 3.53E-54 | NEG | NEG |
| *PRR5* | chr22 | 44701938 | Astrocyte 1 | -10 | 7.94 | 0.000 | promoter-TSS | DEGs, DAGs | -1.231 | 1.85E-70 | NEG | NEG |
| *TMEM184B* | chr22 | 38270565 | Astrocyte 1 | 1849 | 3.82 | 0.002 | promoter-TSS | DEGs, DAGs | -1.358 | 2.93E-66 | NEG | NEG |
| *MIRLET7BHG* | chr22 | 46081182 | Astrocyte 1 | -4565 | 4.13 | 0.009 | Intergenic | DEGs, DAGs | -1.225 | 1.31E-57 | NEG | NEG |
| *MIRLET7BHG* | chr22 | 46084871 | Astrocyte 1 | -876 | 4.05 | 0.018 | promoter-TSS | DEGs, DAGs | -1.225 | 1.31E-57 | NEG | NEG |
| *TMEM184B* | chr22 | 38272794 | Astrocyte 1 | -10 | 2.43 | 0.019 | promoter-TSS | DEGs, DAGs | -1.358 | 2.93E-66 | NEG | NEG |
| *BCR* | chr22 | 23204653 | Astrocyte 1 | 24538 | 2.70 | 0.020 | intron | DEGs, DAGs | -1.334 | 3.37E-67 | NEG | NEG |
| *CERK* | chr22 | 46737558 | Astrocyte 1 | 453 | 3.79 | 0.021 | promoter-TSS | DEGs, DAGs | -1.220 | 2.67E-27 | NEG | NEG |
| *KIAA1671* | chr22 | 25030575 | Astrocyte 1 | 2851 | 5.20 | 0.026 | intron | DEGs, DAGs | -1.489 | 5.75E-107 | NEG | NEG |
| *TCF20* | chr22 | 42229458 | Astrocyte 1 | -14266 | 13.63 | 0.031 | Intergenic | DEGs, DAGs | -1.233 | 3.20E-41 | NEG | NEG |
| *KIAA1671* | chr22 | 24982638 | Astrocyte 1 | 30158 | 4.13 | 0.034 | intron | DEGs, DAGs | -1.489 | 5.75E-107 | NEG | NEG |
| *KIAA1671* | chr22 | 24965980 | Astrocyte 1 | 13500 | 3.11 | 0.036 | intron | DEGs, DAGs | -1.489 | 5.75E-107 | NEG | NEG |
| *BCR* | chr22 | 23207640 | Astrocyte 1 | 27525 | 14.25 | 0.043 | intron | DEGs, DAGs | -1.334 | 3.37E-67 | NEG | NEG |
| *KIAA1671* | chr22 | 25110164 | Astrocyte 1 | -1428 | 11.65 | 0.043 | promoter-TSS | DEGs, DAGs | -1.489 | 5.75E-107 | NEG | NEG |
| *TMEM184B* | chr22 | 38269021 | Astrocyte 1 | 3393 | 7.77 | 0.043 | intron | DEGs, DAGs | -1.358 | 2.93E-66 | NEG | NEG |
| *KIAA1671* | chr22 | 24976470 | Astrocyte 1 | 23990 | 2.61 | 0.046 | intron | DEGs, DAGs | -1.489 | 5.75E-107 | NEG | NEG |
| *KREMEN1* | chr22 | 29073775 | Astrocyte 1 | 894 | 3.33 | 0.049 | promoter-TSS | DEGs, DAGs | -1.250 | 7.80E-24 | NEG | NEG |
| *BCR* | chr22 | 23203487 | Astrocyte 1 | 23372 | 3.90 | 0.049 | intron | DEGs, DAGs | -1.334 | 3.37E-67 | NEG | NEG |
| *HSPB1* | chr7 | 76314257 | Astrocyte 1 | 11963 | 2.23 | 0.000 | Intergenic | DEGs, DAGs | 1.422 | 1.56E-31 | NEG | NEG |
| *ZMIZ1* | chr10 | 78913741 | Astrocyte 1 | -155044 | 3.05 | 0.000 | Intergenic | DEGs, DAGs | -1.210 | 7.98E-52 | NEG | NEG |
| *CHMP6* | chr17 | 80773465 | Astrocyte 1 | -218126 | 7.44 | 0.000 | Intergenic | DEGs, DAGs | -1.335 | 5.05E-55 | NEG | NEG |
| *ZNF98* | chr19 | 22532306 | Astrocyte 1 | -110210 | 3.32 | 0.000 | Intergenic | DEGs, DAGs | 1.677 | 2.91E-89 | NEG | NEG |
| *SAFB2* | chr19 | 5586055 | Astrocyte 1 | 36675 | 2.91 | 0.000 | Intergenic | DEGs, DAGs | 1.290 | 7.32E-88 | NEG | NEG |

| | | | | | | | | | | | | |
|---|---|---|---|---|---|---|---|---|---|---|---|---|
| *NEK6* | chr9 | 124295915 | Astrocyte 1 | 38202 | 3.95 | 0.000 | intron | DEGs, DAGs | -1.406 | 2.80E-71 | NEG | NEG |
| *FTL* | chr19 | 48965056 | Astrocyte 1 | 5 | 2.69 | 0.000 | promoter-TSS | DEGs, DAGs | 1.463 | 2.53E-34 | NEG | NEG |
| *TUBA1A* | chr12 | 49188882 | Astrocyte 1 | 192 | 6.44 | 0.000 | promoter-TSS | DEGs, DAGs | -1.271 | 1.05E-52 | NEG | NEG |
| *SH3PXD2A* | chr10 | 103727360 | Astrocyte 1 | 127796 | 2.35 | 0.000 | intron | DEGs, DAGs | -1.210 | 2.36E-41 | NEG | NEG |
| *TTC7A* | chr2 | 46940895 | Astrocyte 1 | -141 | 5.83 | 0.000 | promoter-TSS | DEGs, DAGs | -1.221 | 1.84E-61 | NEG | NEG |
| *ZMIZ1* | chr10 | 78178946 | Astrocyte 1 | -889839 | 3.97 | 0.000 | Intergenic | DEGs, DAGs | -1.210 | 7.98E-52 | NEG | NEG |
| *GADD45B* | chr19 | 2546445 | Astrocyte 1 | 70568 | 2.80 | 0.001 | Intergenic | DEGs, DAGs | -1.226 | 1.00E-23 | NEG | NEG |
| *SH3RF3* | chr2 | 109284802 | Astrocyte 1 | 155704 | 6.20 | 0.001 | intron | DEGs, DAGs | -1.237 | 1.77E-60 | NEG | NEG |
| *RXRA* | chr9 | 134163620 | Astrocyte 1 | -162710 | 11.68 | 0.002 | Intergenic | DEGs, DAGs | -1.215 | 5.85E-67 | NEG | NEG |
| *GSE1* | chr16 | 85202667 | Astrocyte 1 | -410299 | 3.14 | 0.002 | intron | DEGs, DAGs | -1.237 | 3.56E-37 | NEG | NEG |
| *TMEM184B* | chr22 | 38270565 | Astrocyte 1 | 2219 | 3.82 | 0.002 | intron | DEGs, DAGs | -1.358 | 2.93E-66 | NEG | NEG |
| *FGFR3* | chr4 | 1797770 | Astrocyte 1 | 4708 | 2.07 | 0.002 | intron | DEGs, DAGs | 1.330 | 1.07E-88 | NEG | NEG |
| *GSE1* | chr16 | 85328300 | Astrocyte 1 | -284666 | 3.41 | 0.002 | intron | DEGs, DAGs | -1.237 | 3.56E-37 | NEG | NEG |
| *SLC43A2* | chr17 | 1629928 | Astrocyte 1 | -1837 | 7.24 | 0.002 | promoter-TSS | DEGs, DAGs | -1.326 | 7.93E-20 | NEG | NEG |
| *NEK6* | chr9 | 124291677 | Astrocyte 1 | 33964 | 4.41 | 0.002 | intron | DEGs, DAGs | -1.406 | 2.80E-71 | NEG | NEG |
| *GLI2* | chr2 | 120758032 | Astrocyte 1 | -34127 | 4.36 | 0.002 | intron | DEGs, DAGs | -1.211 | 5.59E-53 | NEG | NEG |
| *KLF2* | chr19 | 16272869 | Astrocyte 1 | -51698 | 3.98 | 0.003 | Intergenic | DEGs, DAGs | -1.391 | 2.76E-85 | NEG | NEG |
| *ZFP36L1* | chr14 | 68568921 | Astrocyte 1 | 224069 | 2.99 | 0.003 | Intergenic | DEGs, DAGs | -1.316 | 3.72E-156 | NEG | NEG |
| *TMEM184B* | chr22 | 38242950 | Astrocyte 1 | 29834 | 2.60 | 0.003 | intron | DEGs, DAGs | -1.358 | 2.93E-66 | NEG | NEG |
| *KANSL1* | chr17 | 46193231 | Astrocyte 1 | -283 | 3.37 | 0.003 | promoter-TSS | DEGs, DAGs | 1.265 | 8.43E-68 | NEG | NEG |
| *IGSF11* | chr3 | 118991743 | Astrocyte 1 | 42895 | 2.55 | 0.003 | intron | DEGs, DAGs | 1.413 | 2.49E-98 | NEG | NEG |
| *BCR* | chr22 | 23225567 | Astrocyte 1 | 45607 | 3.38 | 0.003 | intron | DEGs, DAGs | -1.334 | 3.37E-67 | NEG | NEG |
| *CHMP6* | chr17 | 80873000 | Astrocyte 1 | -118591 | 4.60 | 0.003 | Intergenic | DEGs, DAGs | -1.335 | 5.05E-55 | NEG | NEG |
| *CAPZB* | chr1 | 19405759 | Astrocyte 1 | 78655 | 5.47 | 0.003 | intron | DEGs, DAGs | -1.254 | 1.72E-60 | NEG | NEG |
| *DNM2* | chr19 | 10717210 | Astrocyte 1 | -671 | 2.67 | 0.004 | promoter-TSS | DEGs, DAGs | -1.304 | 3.49E-59 | NEG | NEG |
| *FGFR3* | chr4 | 1746133 | Astrocyte 1 | -46929 | 3.80 | 0.004 | Intergenic | DEGs, DAGs | 1.330 | 1.07E-88 | NEG | NEG |
| *AJAP1* | chr1 | 4108707 | Astrocyte 1 | -546088 | 6.72 | 0.004 | Intergenic | DEGs, DAGs | -1.255 | 3.35E-59 | NEG | NEG |
| *GADD45B* | chr19 | 2587958 | Astrocyte 1 | 112081 | 3.65 | 0.004 | Intergenic | DEGs, DAGs | -1.226 | 1.00E-23 | NEG | NEG |
| *ELN* | chr7 | 73876904 | Astrocyte 1 | -150635 | 8.21 | 0.004 | Intergenic | DEGs, DAGs | -1.204 | 2.59E-10 | NEG | NEG |
| *GTF2IRD1* | chr7 | 74430833 | Astrocyte 1 | -22887 | 2.76 | 0.005 | Intergenic | DEGs, DAGs | -1.348 | 5.20E-58 | NEG | NEG |
| *KREMEN1* | chr22 | 28976867 | Astrocyte 1 | -96001 | 7.60 | 0.005 | Intergenic | DEGs, DAGs | -1.250 | 7.80E-24 | NEG | NEG |
| *NFASC* | chr1 | 204685192 | Astrocyte 1 | -143212 | 3.38 | 0.005 | Intergenic | DEGs, DAGs | -1.293 | 9.92E-29 | NEG | NEG |
| *SLC22A23* | chr6 | 3456974 | Astrocyte 1 | -665 | 2.34 | 0.005 | promoter-TSS | DEGs, DAGs | -1.391 | 1.05E-91 | NEG | NEG |
| *ZMIZ1* | chr10 | 78248774 | Astrocyte 1 | -820011 | 4.75 | 0.005 | Intergenic | DEGs, DAGs | -1.210 | 7.98E-52 | NEG | NEG |

| | | | | | | | | | | | | |
|---|---|---|---|---|---|---|---|---|---|---|---|---|
| *CHMP6* | chr17 | 80835931 | Astrocyte 1 | -155660 | 4.35 | 0.006 | Intergenic | DEGs, DAGs | -1.335 | 5.05E-55 | NEG | NEG |
| *GADD45B* | chr19 | 2579327 | Astrocyte 1 | 103450 | 3.19 | 0.006 | Intergenic | DEGs, DAGs | -1.226 | 1.00E-23 | NEG | NEG |
| *SIPA1L3* | chr19 | 37980467 | Astrocyte 1 | 73489 | 6.44 | 0.006 | intron | DEGs, DAGs | -1.326 | 9.22E-75 | NEG | NEG |
| *ADAMTS10* | chr19 | 8609337 | Astrocyte 1 | -1322 | 3.58 | 0.006 | promoter-TSS | DEGs, DAGs | -1.436 | 1.77E-39 | NEG | NEG |
| *GRIN2C* | chr17 | 74833954 | Astrocyte 1 | 25671 | 2.90 | 0.006 | Intergenic | DEGs, DAGs | -1.220 | 1.95E-92 | NEG | NEG |
| *CHMP6* | chr17 | 80826981 | Astrocyte 1 | -164610 | 5.76 | 0.006 | Intergenic | DEGs, DAGs | -1.335 | 5.05E-55 | NEG | NEG |
| *SDC4* | chr20 | 45344966 | Astrocyte 1 | 3208 | 11.49 | 0.006 | intron | DEGs, DAGs | 1.396 | 1.82E-125 | NEG | NEG |
| *SAFB2* | chr19 | 5586638 | Astrocyte 1 | 36092 | 9.50 | 0.006 | exon | DEGs, DAGs | 1.290 | 7.32E-88 | NEG | NEG |
| *SAFB2* | chr19 | 5586638 | Astrocyte 1 | 36092 | 9.50 | 0.006 | Intergenic | DEGs, DAGs | 1.290 | 7.32E-88 | NEG | NEG |
| *CHMP6* | chr17 | 80948636 | Astrocyte 1 | -42955 | 11.46 | 0.007 | Intergenic | DEGs, DAGs | -1.335 | 5.05E-55 | NEG | NEG |
| *KANSL1* | chr17 | 46193828 | Astrocyte 1 | -880 | 2.49 | 0.007 | promoter-TSS | DEGs, DAGs | 1.265 | 8.43E-68 | NEG | NEG |
| *EYA2* | chr20 | 46894534 | Astrocyte 1 | 160 | 2.45 | 0.008 | promoter-TSS | DEGs, DAGs | -1.296 | 1.28E-29 | NEG | NEG |
| *LINGO1* | chr15 | 77589839 | Astrocyte 1 | 42401 | 4.75 | 0.008 | Intergenic | DEGs, DAGs | -1.338 | 2.43E-31 | NEG | NEG |
| *SNED1* | chr2 | 240968988 | Astrocyte 1 | -29600 | 3.45 | 0.008 | Intergenic | DEGs, DAGs | -1.922 | 8.41E-75 | NEG | NEG |
| *FBXO2* | chr1 | 11648759 | Astrocyte 1 | 5673 | 4.12 | 0.008 | exon | DEGs, DAGs | -1.338 | 1.24E-53 | NEG | NEG |
| *FBXO2* | chr1 | 11648759 | Astrocyte 1 | 5673 | 4.12 | 0.008 | intron | DEGs, DAGs | -1.338 | 1.24E-53 | NEG | NEG |
| *GRIN2C* | chr17 | 74827045 | Astrocyte 1 | 32580 | 10.58 | 0.008 | Intergenic | DEGs, DAGs | -1.220 | 1.95E-92 | NEG | NEG |
| *GFAP* | chr17 | 44916672 | Astrocyte 1 | -1436 | 2.03 | 0.008 | promoter-TSS | DEGs, DAGs | -1.218 | 3.89E-63 | NEG | NEG |
| *SLC1A2* | chr11 | 35362258 | Astrocyte 1 | 56741 | 11.26 | 0.009 | intron | DEGs, DAGs | 1.587 | 9.94E-114 | NEG | NEG |
| *TTC7A* | chr2 | 46955039 | Astrocyte 1 | 14003 | 3.80 | 0.009 | intron | DEGs, DAGs | -1.221 | 1.84E-61 | NEG | NEG |
| *CHST11* | chr12 | 104668932 | Astrocyte 1 | 212209 | 2.01 | 0.009 | intron | DEGs, DAGs | -1.280 | 4.85E-42 | NEG | NEG |
| *SH3PXD2A* | chr10 | 103884592 | Astrocyte 1 | -29436 | 10.31 | 0.009 | Intergenic | DEGs, DAGs | -1.210 | 2.36E-41 | NEG | NEG |
| *ZBTB7C* | chr18 | 48408812 | Astrocyte 1 | -243496 | 3.96 | 0.009 | intron | DEGs, DAGs | -1.205 | 1.24E-44 | NEG | NEG |
| *ZBTB7C* | chr18 | 48408812 | Astrocyte 1 | -243496 | 3.96 | 0.009 | exon | DEGs, DAGs | -1.205 | 1.24E-44 | NEG | NEG |
| *NDRG1* | chr8 | 133453259 | Astrocyte 1 | -155923 | 14.05 | 0.009 | Intergenic | DEGs, DAGs | -1.254 | 2.15E-40 | NEG | NEG |
| *NEK6* | chr9 | 124188229 | Astrocyte 1 | -69484 | 5.03 | 0.010 | Intergenic | DEGs, DAGs | -1.406 | 2.80E-71 | NEG | NEG |
| *NEK6* | chr9 | 124042056 | Astrocyte 1 | -215657 | 2.88 | 0.010 | Intergenic | DEGs, DAGs | -1.406 | 2.80E-71 | NEG | NEG |
| *GRIN2C* | chr17 | 74843349 | Astrocyte 1 | 16276 | 12.91 | 0.010 | exon | DEGs, DAGs | -1.220 | 1.95E-92 | NEG | NEG |
| *GRIN2C* | chr17 | 74843349 | Astrocyte 1 | 16276 | 12.91 | 0.010 | intron | DEGs, DAGs | -1.220 | 1.95E-92 | NEG | NEG |
| *ZMIZ1* | chr10 | 78777381 | Astrocyte 1 | -291404 | 3.08 | 0.010 | Intergenic | DEGs, DAGs | -1.210 | 7.98E-52 | NEG | NEG |
| *GSN* | chr9 | 121317720 | Astrocyte 1 | 18177 | 3.38 | 0.011 | exon | DEGs, DAGs | -1.420 | 3.57E-112 | NEG | NEG |
| *TXNIP* | chr1 | 145985936 | Astrocyte 1 | 10414 | 3.06 | 0.011 | Intergenic | DEGs, DAGs | -1.238 | 4.28E-40 | NEG | NEG |
| *KREMEN1* | chr22 | 29011940 | Astrocyte 1 | -60928 | 2.17 | 0.011 | Intergenic | DEGs, DAGs | -1.250 | 7.80E-24 | NEG | NEG |
| *GRIN2C* | chr17 | 74827671 | Astrocyte 1 | 31954 | 3.05 | 0.011 | Intergenic | DEGs, DAGs | -1.220 | 1.95E-92 | NEG | NEG |

| | | | | | | | | | | | | |
|---|---|---|---|---|---|---|---|---|---|---|---|---|
| *SLC3A2* | chr11 | 62883502 | Astrocyte 1 | 27641 | 2.45 | 0.011 | intron | DEGs, DAGs | 1.394 | 1.07E-83 | NEG | NEG |
| *SLC3A2* | chr11 | 62883502 | Astrocyte 1 | 27641 | 2.45 | 0.011 | exon | DEGs, DAGs | 1.394 | 1.07E-83 | NEG | NEG |
| *ATP2B4* | chr1 | 203508765 | Astrocyte 1 | -117546 | 3.12 | 0.011 | Intergenic | DEGs, DAGs | -1.739 | 4.86E-90 | NEG | NEG |
| *GPC5* | chr13 | 92133332 | Astrocyte 1 | 734975 | 2.25 | 0.011 | intron | DEGs, DAGs | 1.491 | 1.00E-120 | NEG | NEG |
| *GADD45B* | chr19 | 2488825 | Astrocyte 1 | 12948 | 2.23 | 0.011 | Intergenic | DEGs, DAGs | -1.226 | 1.00E-23 | NEG | NEG |
| *ANGPTL4* | chr19 | 8343054 | Astrocyte 1 | -20847 | 3.02 | 0.011 | Intergenic | DEGs, DAGs | -1.485 | 1.12E-44 | NEG | NEG |
| *ABCA3* | chr16 | 2340524 | Astrocyte 1 | -39 | 2.27 | 0.012 | promoter-TSS | DEGs, DAGs | -1.274 | 1.11E-56 | NEG | NEG |
| *BCR* | chr22 | 23281747 | Astrocyte 1 | 101787 | 2.12 | 0.012 | intron | DEGs, DAGs | -1.334 | 3.37E-67 | NEG | NEG |
| *ZMIZ1* | chr10 | 78433213 | Astrocyte 1 | -635572 | 6.33 | 0.013 | Intergenic | DEGs, DAGs | -1.210 | 7.98E-52 | NEG | NEG |
| *SLC39A11* | chr17 | 72943707 | Astrocyte 1 | 148731 | 11.31 | 0.013 | Intergenic | DEGs, DAGs | -1.460 | 1.51E-50 | NEG | NEG |
| *SLC39A11* | chr17 | 72943707 | Astrocyte 1 | 148731 | 11.31 | 0.013 | intron | DEGs, DAGs | -1.460 | 1.51E-50 | NEG | NEG |
| *GADD45B* | chr19 | 2587381 | Astrocyte 1 | 111504 | 7.53 | 0.013 | Intergenic | DEGs, DAGs | -1.226 | 1.00E-23 | NEG | NEG |
| *ZMIZ1* | chr10 | 78656702 | Astrocyte 1 | -412083 | 2.03 | 0.013 | Intergenic | DEGs, DAGs | -1.210 | 7.98E-52 | NEG | NEG |
| *ZMIZ1* | chr10 | 78331353 | Astrocyte 1 | -737432 | 3.11 | 0.013 | Intergenic | DEGs, DAGs | -1.210 | 7.98E-52 | NEG | NEG |
| *CCDC3* | chr10 | 12822606 | Astrocyte 1 | 178841 | 5.66 | 0.013 | Intergenic | DEGs, DAGs | -1.268 | 3.04E-28 | NEG | NEG |
| *ZMIZ1* | chr10 | 78407522 | Astrocyte 1 | -661263 | 13.05 | 0.013 | Intergenic | DEGs, DAGs | -1.210 | 7.98E-52 | NEG | NEG |
| *MGRN1* | chr16 | 4525142 | Astrocyte 1 | -99398 | 2.29 | 0.013 | Intergenic | DEGs, DAGs | -1.201 | 4.68E-53 | NEG | NEG |
| *ZMIZ1* | chr10 | 78178411 | Astrocyte 1 | -890374 | 3.94 | 0.014 | Intergenic | DEGs, DAGs | -1.210 | 7.98E-52 | NEG | NEG |
| *GNG7* | chr19 | 2598884 | Astrocyte 1 | 103575 | 3.61 | 0.014 | intron | DEGs, DAGs | -1.468 | 1.72E-60 | NEG | NEG |
| *CHMP6* | chr17 | 80781730 | Astrocyte 1 | -209861 | 4.83 | 0.014 | Intergenic | DEGs, DAGs | -1.335 | 5.05E-55 | NEG | NEG |
| *GFAP* | chr17 | 44919922 | Astrocyte 1 | -4686 | 3.28 | 0.014 | Intergenic | DEGs, DAGs | -1.218 | 3.89E-63 | NEG | NEG |
| *RNF220* | chr1 | 44423504 | Astrocyte 1 | 18560 | 2.75 | 0.014 | intron | DEGs, DAGs | -1.365 | 1.25E-53 | NEG | NEG |
| *GSE1* | chr16 | 85628016 | Astrocyte 1 | 15050 | 4.89 | 0.015 | intron | DEGs, DAGs | -1.237 | 3.56E-37 | NEG | NEG |
| *AGAP1* | chr2 | 235638486 | Astrocyte 1 | 144629 | 12.91 | 0.015 | intron | DEGs, DAGs | -1.270 | 3.22E-66 | NEG | NEG |
| *ITGB5* | chr3 | 124860600 | Astrocyte 1 | 26447 | 16.57 | 0.015 | intron | DEGs, DAGs | -1.229 | 3.35E-32 | NEG | NEG |
| *NWD1* | chr19 | 16714709 | Astrocyte 1 | -5021 | 3.40 | 0.015 | Intergenic | DEGs, DAGs | -1.307 | 3.93E-133 | NEG | NEG |
| *MAP7* | chr6 | 136761480 | Astrocyte 1 | -235517 | 3.26 | 0.015 | Intergenic | DEGs, DAGs | -1.263 | 2.75E-120 | NEG | NEG |
| *ERBIN* | chr5 | 65816658 | Astrocyte 1 | -109865 | 13.08 | 0.015 | Intergenic | DEGs, DAGs | -1.199 | 2.02E-91 | NEG | NEG |
| *NTM* | chr11 | 131373942 | Astrocyte 1 | -537270 | 4.61 | 0.015 | intron | DEGs, DAGs | 1.336 | 7.51E-57 | NEG | NEG |
| *KREMEN1* | chr22 | 28963370 | Astrocyte 1 | -109498 | 7.00 | 0.015 | Intergenic | DEGs, DAGs | -1.250 | 7.80E-24 | NEG | NEG |
| *HSPA1A* | chr6 | 31814460 | Astrocyte 1 | -754 | 3.36 | 0.015 | promoter-TSS | DEGs, DAGs | 1.636 | 2.51E-30 | NEG | NEG |
| *CNTN2* | chr1 | 205061544 | Astrocyte 1 | 18670 | 2.35 | 0.016 | intron | DEGs, DAGs | -1.204 | 5.98E-09 | NEG | NEG |
| *CNTN2* | chr1 | 205061544 | Astrocyte 1 | 18670 | 2.35 | 0.016 | exon | DEGs, DAGs | -1.204 | 5.98E-09 | NEG | NEG |
| *SLC43A2* | chr17 | 1643467 | Astrocyte 1 | -15376 | 17.08 | 0.016 | Intergenic | DEGs, DAGs | -1.326 | 7.93E-20 | NEG | NEG |

| | | | | | | | | | | | | |
|---|---|---|---|---|---|---|---|---|---|---|---|---|
| *GADD45B* | chr19 | 2547033 | Astrocyte 1 | 71156 | 8.83 | 0.017 | Intergenic | DEGs, DAGs | -1.226 | 1.00E-23 | NEG | NEG |
| *MXI1* | chr10 | 110141878 | Astrocyte 1 | -65477 | 3.31 | 0.018 | Intergenic | DEGs, DAGs | -1.250 | 6.79E-114 | NEG | NEG |
| *AGAP1* | chr2 | 235807289 | Astrocyte 1 | 313432 | 3.98 | 0.018 | exon | DEGs, DAGs | -1.270 | 3.22E-66 | NEG | NEG |
| *AGAP1* | chr2 | 235807289 | Astrocyte 1 | 313432 | 3.98 | 0.018 | intron | DEGs, DAGs | -1.270 | 3.22E-66 | NEG | NEG |
| *NDRG1* | chr8 | 133449429 | Astrocyte 1 | -152093 | 3.45 | 0.018 | Intergenic | DEGs, DAGs | -1.254 | 2.15E-40 | NEG | NEG |
| *SPACA6* | chr19 | 51702232 | Astrocyte 1 | 8986 | 8.47 | 0.019 | exon | DEGs, DAGs | -1.326 | 1.98E-81 | NEG | NEG |
| *CHST11* | chr12 | 104565507 | Astrocyte 1 | 108784 | 3.28 | 0.019 | intron | DEGs, DAGs | -1.280 | 4.85E-42 | NEG | NEG |
| *ATOH8* | chr2 | 85796462 | Astrocyte 1 | 42818 | 2.70 | 0.019 | Intergenic | DEGs, DAGs | -1.321 | 3.33E-24 | NEG | NEG |
| *ZMIZ1* | chr10 | 78331871 | Astrocyte 1 | -736914 | 4.39 | 0.019 | Intergenic | DEGs, DAGs | -1.210 | 7.98E-52 | NEG | NEG |
| *SFXN5* | chr2 | 73085572 | Astrocyte 1 | -13986 | 3.74 | 0.019 | Intergenic | DEGs, DAGs | -1.236 | 1.99E-25 | NEG | NEG |
| *DAAM2* | chr6 | 39812302 | Astrocyte 1 | 19534 | 3.27 | 0.019 | intron | DEGs, DAGs | -1.239 | 1.80E-56 | NEG | NEG |
| *ZMIZ1* | chr10 | 78316306 | Astrocyte 1 | -752479 | 3.44 | 0.020 | Intergenic | DEGs, DAGs | -1.210 | 7.98E-52 | NEG | NEG |
| *PURA* | chr5 | 140108994 | Astrocyte 1 | -4879 | 4.45 | 0.020 | intron | DEGs, DAGs | 1.251 | 6.96E-61 | NEG | NEG |
| *ZMIZ1* | chr10 | 78139301 | Astrocyte 1 | -929484 | 2.53 | 0.020 | Intergenic | DEGs, DAGs | -1.210 | 7.98E-52 | NEG | NEG |
| *ZMIZ1* | chr10 | 79067785 | Astrocyte 1 | -1000 | 2.05 | 0.020 | promoter-TSS | DEGs, DAGs | -1.210 | 7.98E-52 | NEG | NEG |
| *KANSL1* | chr17 | 46192162 | Astrocyte 1 | 786 | 2.95 | 0.020 | promoter-TSS | DEGs, DAGs | 1.265 | 8.43E-68 | NEG | NEG |
| *BCR* | chr22 | 23204653 | Astrocyte 1 | 24693 | 2.70 | 0.020 | intron | DEGs, DAGs | -1.334 | 3.37E-67 | NEG | NEG |
| *FRMD4B* | chr3 | 69686932 | Astrocyte 1 | -301109 | -34.81 | 0.020 | Intergenic | DEGs, DAGs | -1.330 | 9.96E-13 | NEG | NEG |
| *RFX4* | chr12 | 106428327 | Astrocyte 1 | -172560 | 2.58 | 0.020 | Intergenic | DEGs, DAGs | -1.217 | 2.70E-58 | NEG | NEG |
| *DCAKD* | chr17 | 45110952 | Astrocyte 1 | -50093 | 2.48 | 0.020 | Intergenic | DEGs, DAGs | -1.244 | 4.92E-39 | NEG | NEG |
| *CHMP6* | chr17 | 80941598 | Astrocyte 1 | -49993 | 12.88 | 0.020 | Intergenic | DEGs, DAGs | -1.335 | 5.05E-55 | NEG | NEG |
| *ATOH8* | chr2 | 85811305 | Astrocyte 1 | 57661 | 10.90 | 0.020 | Intergenic | DEGs, DAGs | -1.321 | 3.33E-24 | NEG | NEG |
| *NRXN1* | chr2 | 50651690 | Astrocyte 1 | 380459 | 2.12 | 0.020 | intron | DEGs, DAGs | 1.383 | 1.48E-83 | NEG | NEG |
| *UHRF2* | chr9 | 6354041 | Astrocyte 1 | -59032 | 3.66 | 0.021 | Intergenic | DEGs, DAGs | -1.300 | 2.52E-47 | NEG | NEG |
| *GNA14* | chr9 | 77515229 | Astrocyte 1 | 132828 | 2.97 | 0.022 | intron | DEGs, DAGs | -1.421 | 1.11E-90 | NEG | NEG |
| *STK24* | chr13 | 98672982 | Astrocyte 1 | -151234 | 2.25 | 0.022 | Intergenic | DEGs, DAGs | -1.224 | 1.41E-45 | NEG | NEG |
| *FOXN3* | chr14 | 89503483 | Astrocyte 1 | 115397 | 30.05 | 0.023 | intron | DEGs, DAGs | -1.226 | 9.17E-35 | NEG | NEG |
| *RGS9* | chr17 | 65078569 | Astrocyte 1 | -58655 | 2.49 | 0.023 | Intergenic | DEGs, DAGs | -1.199 | 1.97E-12 | NEG | NEG |
| *CHMP6* | chr17 | 80577459 | Astrocyte 1 | -414132 | 14.27 | 0.023 | Intergenic | DEGs, DAGs | -1.335 | 5.05E-55 | NEG | NEG |
| *FAM13C* | chr10 | 59622277 | Astrocyte 1 | -259977 | 2.73 | 0.023 | Intergenic | DEGs, DAGs | -1.383 | 1.95E-68 | NEG | NEG |
| *GADD45B* | chr19 | 2578435 | Astrocyte 1 | 102558 | 5.74 | 0.024 | Intergenic | DEGs, DAGs | -1.226 | 1.00E-23 | NEG | NEG |
| *RHOT1* | chr17 | 32103407 | Astrocyte 1 | -38809 | 7.70 | 0.024 | Intergenic | DEGs, DAGs | -1.219 | 7.94E-61 | NEG | NEG |
| *NEK6* | chr9 | 124110256 | Astrocyte 1 | -147457 | 4.71 | 0.024 | Intergenic | DEGs, DAGs | -1.406 | 2.80E-71 | NEG | NEG |
| *KREMEN1* | chr22 | 29040623 | Astrocyte 1 | -32245 | 4.67 | 0.024 | Intergenic | DEGs, DAGs | -1.250 | 7.80E-24 | NEG | NEG |

| | | | | | | | | | | | | |
|---|---|---|---|---|---|---|---|---|---|---|---|---|
| *SREBF1* | chr17 | 17877952 | Astrocyte 1 | -41215 | 6.09 | 0.025 | Intergenic | DEGs, DAGs | 1.273 | 7.14E-76 | NEG | NEG |
| *MACF1* | chr1 | 39082823 | Astrocyte 1 | -1094 | 6.76 | 0.025 | promoter-TSS | DEGs, DAGs | 1.266 | 2.76E-75 | NEG | NEG |
| *GNG7* | chr19 | 2611132 | Astrocyte 1 | 91327 | 4.77 | 0.025 | intron | DEGs, DAGs | -1.468 | 1.72E-60 | NEG | NEG |
| *GRIN2C* | chr17 | 74834761 | Astrocyte 1 | 24864 | 2.88 | 0.026 | Intergenic | DEGs, DAGs | -1.220 | 1.95E-92 | NEG | NEG |
| *SLC39A11* | chr17 | 73156148 | Astrocyte 1 | -63710 | 3.24 | 0.026 | Intergenic | DEGs, DAGs | -1.460 | 1.51E-50 | NEG | NEG |
| *TMEM132C* | chr12 | 128383556 | Astrocyte 1 | 116403 | 2.82 | 0.026 | intron | DEGs, DAGs | -1.260 | 1.57E-17 | NEG | NEG |
| *IRS2* | chr13 | 109779081 | Astrocyte 1 | 7237 | 5.65 | 0.026 | intron | DEGs, DAGs | -1.271 | 1.91E-58 | NEG | NEG |
| *SLC1A2* | chr11 | 35346145 | Astrocyte 1 | 72854 | 2.92 | 0.026 | intron | DEGs, DAGs | 1.587 | 9.94E-114 | NEG | NEG |
| *HSPB1* | chr7 | 76309982 | Astrocyte 1 | 7688 | 3.13 | 0.026 | Intergenic | DEGs, DAGs | 1.422 | 1.56E-31 | NEG | NEG |
| *TMEM132C* | chr12 | 128408720 | Astrocyte 1 | 141567 | 3.72 | 0.026 | intron | DEGs, DAGs | -1.260 | 1.57E-17 | NEG | NEG |
| *FYN* | chr6 | 112034176 | Astrocyte 1 | -160974 | 2.77 | 0.026 | Intergenic | DEGs, DAGs | -1.395 | 1.56E-94 | NEG | NEG |
| *ADRA1A* | chr8 | 27084256 | Astrocyte 1 | -219101 | 4.18 | 0.026 | Intergenic | DEGs, DAGs | 1.276 | 9.55E-62 | NEG | NEG |
| *GRIN2C* | chr17 | 74822883 | Astrocyte 1 | 36742 | 2.07 | 0.026 | Intergenic | DEGs, DAGs | -1.220 | 1.95E-92 | NEG | NEG |
| *DAAM2* | chr6 | 39811753 | Astrocyte 1 | 18985 | 3.94 | 0.026 | intron | DEGs, DAGs | -1.239 | 1.80E-56 | NEG | NEG |
| *SH3RF3* | chr2 | 109195159 | Astrocyte 1 | 66061 | 8.86 | 0.027 | intron | DEGs, DAGs | -1.237 | 1.77E-60 | NEG | NEG |
| *ZMIZ1* | chr10 | 78846646 | Astrocyte 1 | -222139 | 2.57 | 0.027 | Intergenic | DEGs, DAGs | -1.210 | 7.98E-52 | NEG | NEG |
| *ZMIZ1* | chr10 | 78215637 | Astrocyte 1 | -853148 | 3.92 | 0.027 | Intergenic | DEGs, DAGs | -1.210 | 7.98E-52 | NEG | NEG |
| *CDH23* | chr10 | 71476633 | Astrocyte 1 | 79921 | 3.52 | 0.027 | intron | DEGs, DAGs | -1.419 | 4.96E-48 | NEG | NEG |
| *KREMEN1* | chr22 | 28952106 | Astrocyte 1 | -120762 | 2.11 | 0.027 | Intergenic | DEGs, DAGs | -1.250 | 7.80E-24 | NEG | NEG |
| *ZMIZ1* | chr10 | 78780617 | Astrocyte 1 | -288168 | 2.73 | 0.027 | Intergenic | DEGs, DAGs | -1.210 | 7.98E-52 | NEG | NEG |
| *CACNA2D3* | chr3 | 54146596 | Astrocyte 1 | 24180 | 3.17 | 0.027 | intron | DEGs, DAGs | 1.523 | 8.67E-65 | NEG | NEG |
| *HK1* | chr10 | 69337547 | Astrocyte 1 | 67797 | 51.65 | 0.028 | intron | DEGs, DAGs | -1.215 | 3.79E-51 | NEG | NEG |
| *HK1* | chr10 | 69337547 | Astrocyte 1 | 67797 | 51.65 | 0.028 | exon | DEGs, DAGs | -1.215 | 3.79E-51 | NEG | NEG |
| *DGKG* | chr3 | 186413695 | Astrocyte 1 | -51708 | 2.38 | 0.028 | Intergenic | DEGs, DAGs | 1.329 | 3.97E-63 | NEG | NEG |
| *GNA14* | chr9 | 77852102 | Astrocyte 1 | -204045 | 2.16 | 0.028 | Intergenic | DEGs, DAGs | -1.421 | 1.11E-90 | NEG | NEG |
| *ZMIZ1* | chr10 | 78247711 | Astrocyte 1 | -821074 | 9.02 | 0.028 | Intergenic | DEGs, DAGs | -1.210 | 7.98E-52 | NEG | NEG |
| *CAPZB* | chr1 | 19437795 | Astrocyte 1 | 46619 | 4.39 | 0.028 | intron | DEGs, DAGs | -1.254 | 1.72E-60 | NEG | NEG |
| *PIK3C2B* | chr1 | 204494520 | Astrocyte 1 | -4346 | 2.07 | 0.029 | exon | DEGs, DAGs | -1.203 | 1.30E-27 | NEG | NEG |
| *HSPB1* | chr7 | 76305500 | Astrocyte 1 | 3206 | 3.73 | 0.031 | Intergenic | DEGs, DAGs | 1.422 | 1.56E-31 | NEG | NEG |
| *ZMIZ1* | chr10 | 78282206 | Astrocyte 1 | -786579 | 15.92 | 0.031 | Intergenic | DEGs, DAGs | -1.210 | 7.98E-52 | NEG | NEG |
| *GSE1* | chr16 | 85317054 | Astrocyte 1 | -295912 | 17.07 | 0.031 | intron | DEGs, DAGs | -1.237 | 3.56E-37 | NEG | NEG |
| *SLC39A11* | chr17 | 73088450 | Astrocyte 1 | 3988 | 20.35 | 0.031 | Intergenic | DEGs, DAGs | -1.460 | 1.51E-50 | NEG | NEG |
| *SLC39A11* | chr17 | 73088450 | Astrocyte 1 | 3988 | 20.35 | 0.031 | intron | DEGs, DAGs | -1.460 | 1.51E-50 | NEG | NEG |
| *CHMP6* | chr17 | 80677263 | Astrocyte 1 | -314328 | 13.35 | 0.031 | Intergenic | DEGs, DAGs | -1.335 | 5.05E-55 | NEG | NEG |

| | | | | | | | | | | | | |
|---|---|---|---|---|---|---|---|---|---|---|---|---|
| *FAM171B* | chr2 | 186734877 | Astrocyte 1 | 41156 | 13.95 | 0.031 | intron | DEGs, DAGs | 1.410 | 4.15E-80 | NEG | NEG |
| *TCF20* | chr22 | 42229458 | Astrocyte 1 | -14266 | 13.63 | 0.031 | intron | DEGs, DAGs | -1.233 | 3.20E-41 | NEG | NEG |
| *GTF2IRD1* | chr7 | 74379882 | Astrocyte 1 | -73838 | 2.34 | 0.031 | Intergenic | DEGs, DAGs | -1.348 | 5.20E-58 | NEG | NEG |
| *GNG7* | chr19 | 2633229 | Astrocyte 1 | 69230 | 9.36 | 0.031 | intron | DEGs, DAGs | -1.468 | 1.72E-60 | NEG | NEG |
| *CAPZB* | chr1 | 19429577 | Astrocyte 1 | 54837 | 5.88 | 0.032 | intron | DEGs, DAGs | -1.254 | 1.72E-60 | NEG | NEG |
| *MGRN1* | chr16 | 4537677 | Astrocyte 1 | -86863 | 4.90 | 0.032 | Intergenic | DEGs, DAGs | -1.201 | 4.68E-53 | NEG | NEG |
| *CDH23* | chr10 | 71473854 | Astrocyte 1 | 77142 | 3.68 | 0.032 | intron | DEGs, DAGs | -1.419 | 4.96E-48 | NEG | NEG |
| *FTL* | chr19 | 48970850 | Astrocyte 1 | 5799 | 5.22 | 0.033 | Intergenic | DEGs, DAGs | 1.463 | 2.53E-34 | NEG | NEG |
| *KCNN3* | chr1 | 154911854 | Astrocyte 1 | -41826 | 4.08 | 0.033 | Intergenic | DEGs, DAGs | -1.256 | 1.59E-33 | NEG | NEG |
| *ZFP36L1* | chr14 | 68735751 | Astrocyte 1 | 57239 | 4.00 | 0.033 | Intergenic | DEGs, DAGs | -1.316 | 3.72E-156 | NEG | NEG |
| *KREMEN1* | chr22 | 28919095 | Astrocyte 1 | -153773 | 3.46 | 0.033 | Intergenic | DEGs, DAGs | -1.250 | 7.80E-24 | NEG | NEG |
| *ENO1* | chr1 | 8878552 | Astrocyte 1 | -112 | 2.15 | 0.033 | promoter-TSS | DEGs, DAGs | -1.265 | 2.20E-77 | NEG | NEG |
| *LRIG1* | chr3 | 66481842 | Astrocyte 1 | 18840 | 2.20 | 0.033 | intron | DEGs, DAGs | -1.224 | 1.39E-118 | NEG | NEG |
| *CLMN* | chr14 | 95336826 | Astrocyte 1 | -17170 | 3.57 | 0.034 | Intergenic | DEGs, DAGs | -1.387 | 6.27E-53 | NEG | NEG |
| *HSPA1A* | chr6 | 31813950 | Astrocyte 1 | -1264 | 5.66 | 0.034 | promoter-TSS | DEGs, DAGs | 1.636 | 2.51E-30 | NEG | NEG |
| *ATOH8* | chr2 | 85780662 | Astrocyte 1 | 27018 | 2.75 | 0.034 | intron | DEGs, DAGs | -1.321 | 3.33E-24 | NEG | NEG |
| *ATOH8* | chr2 | 85780662 | Astrocyte 1 | 27018 | 2.75 | 0.034 | exon | DEGs, DAGs | -1.321 | 3.33E-24 | NEG | NEG |
| *OAF* | chr11 | 120169720 | Astrocyte 1 | -40797 | 2.42 | 0.034 | Intergenic | DEGs, DAGs | -1.447 | 7.94E-70 | NEG | NEG |
| *NEK6* | chr9 | 124272809 | Astrocyte 1 | 15096 | 4.12 | 0.034 | intron | DEGs, DAGs | -1.406 | 2.80E-71 | NEG | NEG |
| *EGLN3* | chr14 | 33475607 | Astrocyte 1 | 475224 | 3.29 | 0.035 | Intergenic | DEGs, DAGs | -2.155 | 6.05E-189 | NEG | NEG |
| *EYA2* | chr20 | 46895646 | Astrocyte 1 | 1272 | 2.63 | 0.035 | promoter-TSS | DEGs, DAGs | -1.296 | 1.28E-29 | NEG | NEG |
| *TMEM184B* | chr22 | 38310037 | Astrocyte 1 | -37253 | 2.47 | 0.035 | Intergenic | DEGs, DAGs | -1.358 | 2.93E-66 | NEG | NEG |
| *TMEM184B* | chr22 | 38246591 | Astrocyte 1 | 26193 | 2.26 | 0.035 | intron | DEGs, DAGs | -1.358 | 2.93E-66 | NEG | NEG |
| *INSR* | chr19 | 7197103 | Astrocyte 1 | 96681 | 2.70 | 0.035 | intron | DEGs, DAGs | -1.288 | 9.60E-66 | NEG | NEG |
| *ERCC6L2* | chr9 | 96027262 | Astrocyte 1 | 151811 | -2.95 | 0.035 | intron | DEGs, DAGs | 1.299 | 2.89E-83 | NEG | NEG |
| *FGFR3* | chr4 | 1808860 | Astrocyte 1 | 15798 | 3.51 | 0.036 | exon | DEGs, DAGs | 1.330 | 1.07E-88 | NEG | NEG |
| *FGFR3* | chr4 | 1808860 | Astrocyte 1 | 15798 | 3.51 | 0.036 | Intergenic | DEGs, DAGs | 1.330 | 1.07E-88 | NEG | NEG |
| *SAFB2* | chr19 | 5539297 | Astrocyte 1 | 83433 | 3.13 | 0.037 | Intergenic | DEGs, DAGs | 1.290 | 7.32E-88 | NEG | NEG |
| *TCF20* | chr22 | 42296442 | Astrocyte 1 | -81250 | 2.38 | 0.038 | Intergenic | DEGs, DAGs | -1.233 | 3.20E-41 | NEG | NEG |
| *TCF20* | chr22 | 42296442 | Astrocyte 1 | -81250 | 2.38 | 0.038 | intron | DEGs, DAGs | -1.233 | 3.20E-41 | NEG | NEG |
| *PLEKHA7* | chr11 | 16925096 | Astrocyte 1 | 89066 | 2.03 | 0.038 | intron | DEGs, DAGs | -1.282 | 2.59E-61 | NEG | NEG |
| *RGS9* | chr17 | 65067030 | Astrocyte 1 | -70194 | 8.61 | 0.038 | Intergenic | DEGs, DAGs | -1.199 | 1.97E-12 | NEG | NEG |
| *RXRA* | chr9 | 134316329 | Astrocyte 1 | -10001 | 3.43 | 0.039 | Intergenic | DEGs, DAGs | -1.215 | 5.85E-67 | NEG | NEG |
| *ZMIZ1* | chr10 | 78255967 | Astrocyte 1 | -812818 | 2.41 | 0.040 | Intergenic | DEGs, DAGs | -1.210 | 7.98E-52 | NEG | NEG |

| | | | | | | | | | | | | |
|---|---|---|---|---|---|---|---|---|---|---|---|---|
| *IGSF11* | chr3 | 118972903 | Astrocyte 1 | 61735 | 2.00 | 0.041 | intron | DEGs, DAGs | 1.413 | 2.49E-98 | NEG | NEG |
| *NEK6* | chr9 | 124013712 | Astrocyte 1 | -244001 | 4.45 | 0.042 | Intergenic | DEGs, DAGs | -1.406 | 2.80E-71 | NEG | NEG |
| *SH3GL1* | chr19 | 4402187 | Astrocyte 1 | -1890 | 2.20 | 0.043 | promoter-TSS | DEGs, DAGs | -1.211 | 4.82E-28 | NEG | NEG |
| *SLC43A2* | chr17 | 1628729 | Astrocyte 1 | -638 | 2.50 | 0.043 | promoter-TSS | DEGs, DAGs | -1.326 | 7.93E-20 | NEG | NEG |
| *CDH23* | chr10 | 71543729 | Astrocyte 1 | 147017 | 27.94 | 0.043 | intron | DEGs, DAGs | -1.419 | 4.96E-48 | NEG | NEG |
| *TMEM132B* | chr12 | 125069512 | Astrocyte 1 | -256854 | 13.64 | 0.043 | Intergenic | DEGs, DAGs | -1.209 | 2.02E-22 | NEG | NEG |
| *GSE1* | chr16 | 85194915 | Astrocyte 1 | -418051 | 11.04 | 0.043 | intron | DEGs, DAGs | -1.237 | 3.56E-37 | NEG | NEG |
| *GNG7* | chr19 | 2676051 | Astrocyte 1 | 26408 | 11.81 | 0.043 | intron | DEGs, DAGs | -1.468 | 1.72E-60 | NEG | NEG |
| *GNG7* | chr19 | 2676911 | Astrocyte 1 | 25548 | 12.50 | 0.043 | intron | DEGs, DAGs | -1.468 | 1.72E-60 | NEG | NEG |
| *SIPA1L3* | chr19 | 37979751 | Astrocyte 1 | 72773 | 17.69 | 0.043 | intron | DEGs, DAGs | -1.326 | 9.22E-75 | NEG | NEG |
| *BCR* | chr22 | 23207640 | Astrocyte 1 | 27680 | 14.25 | 0.043 | intron | DEGs, DAGs | -1.334 | 3.37E-67 | NEG | NEG |
| *TMEM184B* | chr22 | 38269021 | Astrocyte 1 | 3763 | 7.77 | 0.043 | intron | DEGs, DAGs | -1.358 | 2.93E-66 | NEG | NEG |
| *ITGB5* | chr3 | 124822077 | Astrocyte 1 | 64970 | 17.74 | 0.043 | intron | DEGs, DAGs | -1.229 | 3.35E-32 | NEG | NEG |
| *SCARA3* | chr8 | 27633557 | Astrocyte 1 | -374 | 2.51 | 0.043 | promoter-TSS | DEGs, DAGs | -1.510 | 1.88E-117 | NEG | NEG |
| *NEK6* | chr9 | 124262539 | Astrocyte 1 | 4826 | 8.37 | 0.043 | intron | DEGs, DAGs | -1.406 | 2.80E-71 | NEG | NEG |
| *NEK6* | chr9 | 124259376 | Astrocyte 1 | 1663 | 2.27 | 0.043 | promoter-TSS | DEGs, DAGs | -1.406 | 2.80E-71 | NEG | NEG |
| *SLC7A11* | chr4 | 138890849 | Astrocyte 1 | -648750 | -7.32 | 0.043 | Intergenic | DEGs, DAGs | 1.384 | 1.16E-13 | NEG | NEG |
| *HIF3A* | chr19 | 46297672 | Astrocyte 1 | 876 | 4.21 | 0.044 | promoter-TSS | DEGs, DAGs | 1.412 | 6.22E-116 | NEG | NEG |
| *BCR* | chr22 | 23301599 | Astrocyte 1 | 121639 | 15.53 | 0.044 | intron | DEGs, DAGs | -1.334 | 3.37E-67 | NEG | NEG |
| *KREMEN1* | chr22 | 28897941 | Astrocyte 1 | -174927 | 6.37 | 0.044 | Intergenic | DEGs, DAGs | -1.250 | 7.80E-24 | NEG | NEG |
| *KLHL32* | chr6 | 97048443 | Astrocyte 1 | 123964 | -7.01 | 0.044 | intron | DEGs, DAGs | -1.198 | 8.80E-79 | NEG | NEG |
| *CERS4* | chr19 | 8251133 | Astrocyte 1 | 42030 | 2.46 | 0.044 | exon | DEGs, DAGs | -1.272 | 3.82E-60 | NEG | NEG |
| *CERS4* | chr19 | 8251133 | Astrocyte 1 | 42030 | 2.46 | 0.044 | intron | DEGs, DAGs | -1.272 | 3.82E-60 | NEG | NEG |
| *FGFR2* | chr10 | 121406751 | Astrocyte 1 | 191402 | 4.37 | 0.044 | Intergenic | DEGs, DAGs | -1.250 | 4.85E-118 | NEG | NEG |
| *GRIN2C* | chr17 | 74828196 | Astrocyte 1 | 31429 | 4.82 | 0.044 | Intergenic | DEGs, DAGs | -1.220 | 1.95E-92 | NEG | NEG |
| *GADD45B* | chr19 | 2495219 | Astrocyte 1 | 19342 | 5.27 | 0.045 | Intergenic | DEGs, DAGs | -1.226 | 1.00E-23 | NEG | NEG |
| *SLC3A2* | chr11 | 62855700 | Astrocyte 1 | -161 | 2.63 | 0.045 | promoter-TSS | DEGs, DAGs | 1.394 | 1.07E-83 | NEG | NEG |
| *GADD45B* | chr19 | 2543535 | Astrocyte 1 | 67658 | 2.63 | 0.045 | Intergenic | DEGs, DAGs | -1.226 | 1.00E-23 | NEG | NEG |
| *PITPNA* | chr17 | 1561744 | Astrocyte 1 | 822 | 4.37 | 0.045 | promoter-TSS | DEGs, DAGs | -1.295 | 7.86E-38 | NEG | NEG |
| *ACSBG1* | chr15 | 78193923 | Astrocyte 1 | 40534 | 3.92 | 0.045 | intron | DEGs, DAGs | -1.241 | 8.93E-79 | NEG | NEG |
| *FGFR3* | chr4 | 1788491 | Astrocyte 1 | -4571 | 6.51 | 0.045 | Intergenic | DEGs, DAGs | 1.330 | 1.07E-88 | NEG | NEG |
| *ZMIZ1* | chr10 | 78294644 | Astrocyte 1 | -774141 | 4.95 | 0.045 | Intergenic | DEGs, DAGs | -1.210 | 7.98E-52 | NEG | NEG |
| *LINGO1* | chr15 | 77541603 | Astrocyte 1 | 90637 | 2.04 | 0.045 | Intergenic | DEGs, DAGs | -1.338 | 2.43E-31 | NEG | NEG |
| *TAF15* | chr17 | 35804695 | Astrocyte 1 | -4510 | 2.12 | 0.045 | Intergenic | DEGs, DAGs | -1.245 | 6.12E-59 | NEG | NEG |

| | | | | | | | | | | | | |
|---|---|---|---|---|---|---|---|---|---|---|---|---|
| *CLEC16A* | chr16 | 11054227 | Astrocyte 1 | 109989 | 5.17 | 0.046 | intron | DEGs, DAGs | -1.228 | 4.44E-87 | NEG | NEG |
| *ACOT11* | chr1 | 54579410 | Astrocyte 1 | 31432 | 2.23 | 0.046 | intron | DEGs, DAGs | -1.244 | 1.85E-35 | NEG | NEG |
| *ZMIZ1* | chr10 | 78434148 | Astrocyte 1 | -634637 | 4.14 | 0.047 | Intergenic | DEGs, DAGs | -1.210 | 7.98E-52 | NEG | NEG |
| *FGFR2* | chr10 | 121539701 | Astrocyte 1 | 58452 | 6.31 | 0.047 | intron | DEGs, DAGs | -1.250 | 4.85E-118 | NEG | NEG |
| *ABCA2* | chr9 | 137028017 | Astrocyte 1 | -77 | 2.15 | 0.047 | promoter-TSS | DEGs, DAGs | -1.319 | 1.32E-43 | NEG | NEG |
| *MXI1* | chr10 | 110212064 | Astrocyte 1 | 4709 | 3.78 | 0.047 | intron | DEGs, DAGs | -1.250 | 6.79E-114 | NEG | NEG |
| *VEGFA* | chr6 | 43770739 | Astrocyte 1 | 282 | 2.39 | 0.048 | promoter-TSS | DEGs, DAGs | 1.284 | 1.68E-130 | NEG | NEG |
| *KREMEN1* | chr22 | 29073775 | Astrocyte 1 | 907 | 3.33 | 0.049 | promoter-TSS | DEGs, DAGs | -1.250 | 7.80E-24 | NEG | NEG |
| *NEK6* | chr9 | 124018110 | Astrocyte 1 | -239603 | 2.50 | 0.049 | Intergenic | DEGs, DAGs | -1.406 | 2.80E-71 | NEG | NEG |
| *BCR* | chr22 | 23203487 | Astrocyte 1 | 23527 | 3.90 | 0.049 | intron | DEGs, DAGs | -1.334 | 3.37E-67 | NEG | NEG |
| *SPACA6* | chr19 | 51690593 | Astrocyte 1 | -2653 | 2.45 | 0.050 | intron | DEGs, DAGs | -1.326 | 1.98E-81 | NEG | NEG |
| *RGS9* | chr17 | 65100567 | Astrocyte 1 | -36657 | 7.10 | 0.000 | Intergenic | DEGs, DAGs | -1.199 | 1.97E-12 | NEG | NEG |
| *CASTOR2* | chr7 | 75046320 | Astrocyte 1 | 81752 | 4.93 | 0.000 | Intergenic | DEGs, DAGs | -1.462 | 1.02E-49 | NEG | NEG |
| *SH3GL1* | chr19 | 4369832 | Astrocyte 1 | 30465 | 8.69 | 0.001 | intron | DEGs, DAGs | -1.211 | 4.82E-28 | NEG | NEG |
| *GNG7* | chr19 | 2546445 | Astrocyte 1 | 156014 | 2.80 | 0.001 | intron | DEGs, DAGs | -1.468 | 1.72E-60 | NEG | NEG |
| *GRIN2C* | chr17 | 74791406 | Astrocyte 1 | 68219 | 4.19 | 0.001 | Intergenic | DEGs, DAGs | -1.220 | 1.95E-92 | NEG | NEG |
| *ZNF98* | chr19 | 22515372 | Astrocyte 1 | -93276 | 9.78 | 0.001 | intron | DEGs, DAGs | 1.677 | 2.91E-89 | NEG | NEG |
| *ERGIC1* | chr5 | 172867295 | Astrocyte 1 | 33270 | 4.48 | 0.001 | intron | DEGs, DAGs | -1.227 | 6.40E-52 | NEG | NEG |
| *FYN* | chr6 | 111658301 | Astrocyte 1 | 214901 | 4.53 | 0.001 | Intergenic | DEGs, DAGs | -1.395 | 1.56E-94 | NEG | NEG |
| *RXRA* | chr9 | 134370038 | Astrocyte 1 | 43708 | 5.79 | 0.001 | intron | DEGs, DAGs | -1.215 | 5.85E-67 | NEG | NEG |
| *FBXO2* | chr1 | 11499891 | Astrocyte 1 | 154541 | 3.10 | 0.001 | Intergenic | DEGs, DAGs | -1.338 | 1.24E-53 | NEG | NEG |
| *FBXO2* | chr1 | 11499260 | Astrocyte 1 | 155172 | 9.94 | 0.002 | Intergenic | DEGs, DAGs | -1.338 | 1.24E-53 | NEG | NEG |
| *MACF1* | chr1 | 39272026 | Astrocyte 1 | 188109 | 5.30 | 0.002 | intron | DEGs, DAGs | 1.266 | 2.76E-75 | NEG | NEG |
| *SDK2* | chr17 | 73341992 | Astrocyte 1 | 301847 | 3.84 | 0.002 | intron | DEGs, DAGs | -1.245 | 4.12E-68 | NEG | NEG |
| *PITPNM2* | chr12 | 123071756 | Astrocyte 1 | 38422 | 10.51 | 0.002 | intron | DEGs, DAGs | -1.207 | 6.27E-56 | NEG | NEG |
| *EYA2* | chr20 | 47319864 | Astrocyte 1 | 425490 | 3.64 | 0.002 | Intergenic | DEGs, DAGs | -1.296 | 1.28E-29 | NEG | NEG |
| *ID3* | chr1 | 23539750 | Astrocyte 1 | 19794 | 3.17 | 0.003 | Intergenic | DEGs, DAGs | -1.839 | 6.61E-109 | NEG | NEG |
| *INO80D* | chr2 | 205728877 | Astrocyte 1 | 357055 | 10.57 | 0.003 | Intergenic | DEGs, DAGs | -1.908 | 2.02E-63 | NEG | NEG |
| *ASPH* | chr8 | 61471355 | Astrocyte 1 | 242954 | 2.95 | 0.003 | Intergenic | DEGs, DAGs | 1.448 | 3.50E-100 | NEG | NEG |
| *SIPA1L3* | chr19 | 38093561 | Astrocyte 1 | 186583 | 19.01 | 0.003 | intron | DEGs, DAGs | -1.326 | 9.22E-75 | NEG | NEG |
| *SLC6A11* | chr3 | 10738800 | Astrocyte 1 | -77150 | 4.09 | 0.003 | Intergenic | DEGs, DAGs | 1.387 | 1.41E-117 | NEG | NEG |
| *RASSF4* | chr10 | 44976405 | Astrocyte 1 | 16863 | -8.72 | 0.003 | intron | DEGs, DAGs | 1.262 | 1.68E-66 | NEG | NEG |
| *CHST3* | chr10 | 72044348 | Astrocyte 1 | 80233 | 2.24 | 0.004 | Intergenic | DEGs, DAGs | -1.231 | 3.60E-28 | NEG | NEG |
| *KCNN3* | chr1 | 154866993 | Astrocyte 1 | 3035 | 2.97 | 0.004 | intron | DEGs, DAGs | -1.256 | 1.59E-33 | NEG | NEG |

| | | | | | | | | | | | | |
|---|---|---|---|---|---|---|---|---|---|---|---|---|
| *SDK2* | chr17 | 73365850 | Astrocyte 1 | 277989 | 8.68 | 0.004 | intron | DEGs, DAGs | -1.245 | 4.12E-68 | NEG | NEG |
| *SLC39A11* | chr17 | 72476453 | Astrocyte 1 | 615985 | 2.11 | 0.004 | Intergenic | DEGs, DAGs | -1.460 | 1.51E-50 | NEG | NEG |
| *GNG7* | chr19 | 2587958 | Astrocyte 1 | 114501 | 3.65 | 0.004 | intron | DEGs, DAGs | -1.468 | 1.72E-60 | NEG | NEG |
| *ZMIZ1* | chr10 | 79282683 | Astrocyte 1 | 213898 | 3.86 | 0.004 | intron | DEGs, DAGs | -1.210 | 7.98E-52 | NEG | NEG |
| *CLIP2* | chr7 | 74430833 | Astrocyte 1 | 141608 | 2.76 | 0.005 | Intergenic | DEGs, DAGs | -1.528 | 1.83E-58 | NEG | NEG |
| *RXRA* | chr9 | 134361775 | Astrocyte 1 | 35445 | 6.58 | 0.006 | intron | DEGs, DAGs | -1.215 | 5.85E-67 | NEG | NEG |
| *EFHD1* | chr2 | 232659199 | Astrocyte 1 | 26221 | 3.34 | 0.006 | intron | DEGs, DAGs | -1.315 | 1.40E-66 | NEG | NEG |
| *GNG7* | chr19 | 2579327 | Astrocyte 1 | 123132 | 3.19 | 0.006 | intron | DEGs, DAGs | -1.468 | 1.72E-60 | NEG | NEG |
| *RFX2* | chr19 | 6058656 | Astrocyte 1 | 51628 | 9.56 | 0.006 | intron | DEGs, DAGs | -1.328 | 3.49E-49 | NEG | NEG |
| *CTBP2* | chr10 | 125060769 | Astrocyte 1 | -33129 | 2.46 | 0.006 | intron | DEGs, DAGs | -1.249 | 4.58E-57 | NEG | NEG |
| *FGF1* | chr5 | 142678528 | Astrocyte 1 | 19292 | 2.09 | 0.006 | intron | DEGs, DAGs | -1.299 | 1.17E-52 | NEG | NEG |
| *SDK2* | chr17 | 73401920 | Astrocyte 1 | 241919 | 2.60 | 0.006 | intron | DEGs, DAGs | -1.245 | 4.12E-68 | NEG | NEG |
| *SIPA1L3* | chr19 | 38090537 | Astrocyte 1 | 183559 | 4.52 | 0.006 | intron | DEGs, DAGs | -1.326 | 9.22E-75 | NEG | NEG |
| *DNM2* | chr19 | 10796614 | Astrocyte 1 | 78733 | 3.27 | 0.006 | intron | DEGs, DAGs | -1.304 | 3.49E-59 | NEG | NEG |
| *GSE1* | chr16 | 85537893 | Astrocyte 1 | -75073 | 7.69 | 0.007 | intron | DEGs, DAGs | -1.237 | 3.56E-37 | NEG | NEG |
| *SLC39A11* | chr17 | 72502000 | Astrocyte 1 | 590438 | 2.23 | 0.007 | Intergenic | DEGs, DAGs | -1.460 | 1.51E-50 | NEG | NEG |
| *RXRA* | chr9 | 134434494 | Astrocyte 1 | 108164 | 2.97 | 0.008 | intron | DEGs, DAGs | -1.215 | 5.85E-67 | NEG | NEG |
| *ENO1* | chr1 | 8889183 | Astrocyte 1 | -10743 | 8.24 | 0.008 | Intergenic | DEGs, DAGs | -1.265 | 2.20E-77 | NEG | NEG |
| *GMPR* | chr6 | 16541319 | Astrocyte 1 | 302989 | 9.65 | 0.008 | Intergenic | DEGs, DAGs | -1.411 | 1.22E-66 | NEG | NEG |
| *MACF1* | chr1 | 39204515 | Astrocyte 1 | 120598 | 17.26 | 0.008 | intron | DEGs, DAGs | 1.266 | 2.76E-75 | NEG | NEG |
| *MACF1* | chr1 | 39204515 | Astrocyte 1 | 120598 | 17.26 | 0.008 | exon | DEGs, DAGs | 1.266 | 2.76E-75 | NEG | NEG |
| *RFX2* | chr19 | 6056646 | Astrocyte 1 | 53638 | 4.05 | 0.008 | intron | DEGs, DAGs | -1.328 | 3.49E-49 | NEG | NEG |
| *KLF15* | chr3 | 126277893 | Astrocyte 1 | 79299 | 7.95 | 0.008 | Intergenic | DEGs, DAGs | -1.209 | 1.08E-55 | NEG | NEG |
| *ITGB5* | chr3 | 124772506 | Astrocyte 1 | 114541 | 3.39 | 0.008 | intron | DEGs, DAGs | -1.229 | 3.35E-32 | NEG | NEG |
| *CD44* | chr11 | 35362258 | Astrocyte 1 | 223327 | 11.26 | 0.009 | Intergenic | DEGs, DAGs | -1.315 | 2.26E-16 | NEG | NEG |
| *CORO2B* | chr15 | 68700480 | Astrocyte 1 | 121761 | 4.44 | 0.009 | intron | DEGs, DAGs | -1.243 | 4.68E-56 | NEG | NEG |
| *SFXN5* | chr2 | 72973546 | Astrocyte 1 | 98040 | 10.14 | 0.009 | intron | DEGs, DAGs | -1.236 | 1.99E-25 | NEG | NEG |
| *KYAT1* | chr9 | 128881709 | Astrocyte 1 | 116 | 2.31 | 0.010 | promoter-TSS | DEGs, DAGs | -1.251 | 8.37E-32 | NEG | NEG |
| *CUX1* | chr7 | 101733828 | Astrocyte 1 | -81933 | 3.04 | 0.010 | Intergenic | DEGs, DAGs | -1.271 | 8.56E-65 | NEG | NEG |
| *LHX2* | chr9 | 124188229 | Astrocyte 1 | 176869 | 5.03 | 0.010 | Intergenic | DEGs, DAGs | -1.389 | 7.39E-66 | NEG | NEG |
| *LHX2* | chr9 | 124042056 | Astrocyte 1 | 30696 | 2.88 | 0.010 | Intergenic | DEGs, DAGs | -1.389 | 7.39E-66 | NEG | NEG |
| *COL27A1* | chr9 | 114216024 | Astrocyte 1 | 60714 | 7.95 | 0.010 | intron | DEGs, DAGs | -1.357 | 1.23E-46 | NEG | NEG |
| *ZBTB16* | chr11 | 114228702 | Astrocyte 1 | 169359 | 4.39 | 0.011 | intron | DEGs, DAGs | -1.313 | 1.27E-83 | NEG | NEG |
| *ATP1A2* | chr1 | 160122920 | Astrocyte 1 | 7397 | 5.84 | 0.011 | intron | DEGs, DAGs | 1.507 | 3.27E-138 | NEG | NEG |

| | | | | | | | | | | | | |
|---|---|---|---|---|---|---|---|---|---|---|---|---|
| *SNED1* | chr2 | 241079055 | Astrocyte 1 | 80467 | 12.49 | 0.011 | intron | DEGs, DAGs | -1.922 | 8.41E-75 | NEG | NEG |
| *MTRNR2L8* | chr11 | 10478614 | Astrocyte 1 | 30325 | 2.59 | 0.011 | Intergenic | DEGs, DAGs | 1.356 | 1.03E-60 | NEG | NEG |
| *HHIPL1* | chr14 | 99675725 | Astrocyte 1 | 30865 | 12.96 | 0.011 | exon | DEGs, DAGs | -1.226 | 2.56E-50 | NEG | NEG |
| *HHIPL1* | chr14 | 99675725 | Astrocyte 1 | 30865 | 12.96 | 0.011 | intron | DEGs, DAGs | -1.226 | 2.56E-50 | NEG | NEG |
| *MRAS* | chr3 | 138357470 | Astrocyte 1 | 10023 | 2.51 | 0.011 | intron | DEGs, DAGs | -1.451 | 3.05E-64 | NEG | NEG |
| *MARK4* | chr19 | 45294609 | Astrocyte 1 | 43601 | 3.65 | 0.011 | intron | DEGs, DAGs | -1.366 | 3.18E-67 | NEG | NEG |
| *DAPK1* | chr9 | 87613730 | Astrocyte 1 | 116099 | 3.29 | 0.011 | intron | DEGs, DAGs | -1.259 | 1.60E-62 | NEG | NEG |
| *KCNN3* | chr1 | 154776183 | Astrocyte 1 | 93845 | 3.84 | 0.011 | intron | DEGs, DAGs | -1.256 | 1.59E-33 | NEG | NEG |
| *GNG7* | chr19 | 2488825 | Astrocyte 1 | 213634 | 2.23 | 0.011 | Intergenic | DEGs, DAGs | -1.468 | 1.72E-60 | NEG | NEG |
| *CHST3* | chr10 | 72078770 | Astrocyte 1 | 114655 | 13.57 | 0.012 | Intergenic | DEGs, DAGs | -1.231 | 3.60E-28 | NEG | NEG |
| *GTF2IRD1* | chr7 | 74542465 | Astrocyte 1 | 88745 | 9.78 | 0.012 | intron | DEGs, DAGs | -1.348 | 5.20E-58 | NEG | NEG |
| *TEAD1* | chr11 | 12941548 | Astrocyte 1 | 267207 | 6.43 | 0.013 | exon | DEGs, DAGs | -1.229 | 5.49E-75 | NEG | NEG |
| *TEAD1* | chr11 | 12941548 | Astrocyte 1 | 267207 | 6.43 | 0.013 | intron | DEGs, DAGs | -1.229 | 5.49E-75 | NEG | NEG |
| *GNG7* | chr19 | 2587381 | Astrocyte 1 | 115078 | 7.53 | 0.013 | intron | DEGs, DAGs | -1.468 | 1.72E-60 | NEG | NEG |
| *SH3PXD2B* | chr5 | 172452765 | Astrocyte 1 | 1508 | 5.01 | 0.013 | promoter-TSS | DEGs, DAGs | -1.286 | 4.62E-47 | NEG | NEG |
| *EFHD1* | chr2 | 232669038 | Astrocyte 1 | 36060 | 10.89 | 0.013 | intron | DEGs, DAGs | -1.315 | 1.40E-66 | NEG | NEG |
| *GADD45B* | chr19 | 2598884 | Astrocyte 1 | 123007 | 3.61 | 0.014 | Intergenic | DEGs, DAGs | -1.226 | 1.00E-23 | NEG | NEG |
| *KLF2* | chr19 | 16457812 | Astrocyte 1 | 133245 | 6.83 | 0.014 | Intergenic | DEGs, DAGs | -1.391 | 2.76E-85 | NEG | NEG |
| *PRICKLE2* | chr3 | 64071990 | Astrocyte 1 | 153241 | 2.06 | 0.014 | Intergenic | DEGs, DAGs | -1.274 | 3.76E-44 | NEG | NEG |
| *TMEM184B* | chr22 | 38214584 | Astrocyte 1 | 58200 | 5.03 | 0.014 | Intergenic | DEGs, DAGs | -1.358 | 2.93E-66 | NEG | NEG |
| *ZFP36L1* | chr14 | 68248369 | Astrocyte 1 | 544621 | 2.14 | 0.014 | Intergenic | DEGs, DAGs | -1.316 | 3.72E-156 | NEG | NEG |
| *ZMIZ1* | chr10 | 79273507 | Astrocyte 1 | 204722 | 5.28 | 0.014 | intron | DEGs, DAGs | -1.210 | 7.98E-52 | NEG | NEG |
| *TMEM108* | chr3 | 133041158 | Astrocyte 1 | 3017 | 2.48 | 0.014 | intron | DEGs, DAGs | 1.294 | 1.26E-60 | NEG | NEG |
| *CHMP6* | chr17 | 81025180 | Astrocyte 1 | 33589 | 5.33 | 0.015 | Intergenic | DEGs, DAGs | -1.335 | 5.05E-55 | NEG | NEG |
| *KCNN3* | chr1 | 154751500 | Astrocyte 1 | 118528 | 2.33 | 0.015 | intron | DEGs, DAGs | -1.256 | 1.59E-33 | NEG | NEG |
| *GSE1* | chr16 | 85370720 | Astrocyte 1 | -242246 | 2.65 | 0.015 | intron | DEGs, DAGs | -1.237 | 3.56E-37 | NEG | NEG |
| *COL5A3* | chr19 | 9953946 | Astrocyte 1 | 56275 | 2.13 | 0.015 | Intergenic | DEGs, DAGs | 1.581 | 7.92E-160 | NEG | NEG |
| *CACNA1C* | chr12 | 2396468 | Astrocyte 1 | 343155 | 3.48 | 0.015 | intron | DEGs, DAGs | -1.396 | 3.63E-93 | NEG | NEG |
| *APBB2* | chr4 | 40845172 | Astrocyte 1 | 369110 | 2.07 | 0.015 | intron | DEGs, DAGs | -1.215 | 4.72E-59 | NEG | NEG |
| *PITPNM2* | chr12 | 123043412 | Astrocyte 1 | 66766 | 3.83 | 0.016 | intron | DEGs, DAGs | -1.207 | 6.27E-56 | NEG | NEG |
| *GSN* | chr9 | 121289239 | Astrocyte 1 | -10304 | 3.43 | 0.017 | intron | DEGs, DAGs | -1.420 | 3.57E-112 | NEG | NEG |
| *DYSF* | chr2 | 72144696 | Astrocyte 1 | 678244 | 10.07 | 0.017 | Intergenic | DEGs, DAGs | -1.253 | 6.88E-21 | NEG | NEG |
| *GNG7* | chr19 | 2547033 | Astrocyte 1 | 155426 | 8.83 | 0.017 | intron | DEGs, DAGs | -1.468 | 1.72E-60 | NEG | NEG |
| *DNMT3A* | chr2 | 25305730 | Astrocyte 1 | 36610 | 2.58 | 0.017 | intron | DEGs, DAGs | -1.415 | 3.81E-61 | NEG | NEG |

| | | | | | | | | | | | | |
|---|---|---|---|---|---|---|---|---|---|---|---|---|
| *PRKCE* | chr2 | 45874186 | Astrocyte 1 | 222662 | 4.77 | 0.018 | intron | DEGs, DAGs | -1.219 | 4.54E-49 | NEG | NEG |
| *MACF1* | chr1 | 39142705 | Astrocyte 1 | 58788 | 2.32 | 0.018 | intron | DEGs, DAGs | 1.266 | 2.76E-75 | NEG | NEG |
| *CACNA1C* | chr12 | 2163467 | Astrocyte 1 | 110154 | 3.89 | 0.018 | intron | DEGs, DAGs | -1.396 | 3.63E-93 | NEG | NEG |
| *ELN* | chr7 | 74067778 | Astrocyte 1 | 40239 | 3.10 | 0.018 | intron | DEGs, DAGs | -1.204 | 2.59E-10 | NEG | NEG |
| *SSBP3* | chr1 | 54275171 | Astrocyte 1 | 130998 | 3.15 | 0.018 | intron | DEGs, DAGs | -1.204 | 8.16E-47 | NEG | NEG |
| *ZMIZ1* | chr10 | 79195360 | Astrocyte 1 | 126575 | 3.00 | 0.018 | intron | DEGs, DAGs | -1.210 | 7.98E-52 | NEG | NEG |
| *ZBTB16* | chr11 | 114072097 | Astrocyte 1 | 12754 | 2.63 | 0.018 | intron | DEGs, DAGs | -1.313 | 1.27E-83 | NEG | NEG |
| *FGF1* | chr5 | 142611006 | Astrocyte 1 | 86814 | 2.21 | 0.018 | intron | DEGs, DAGs | -1.299 | 1.17E-52 | NEG | NEG |
| *SLC39A11* | chr17 | 72574046 | Astrocyte 1 | 518392 | 7.36 | 0.019 | Intergenic | DEGs, DAGs | -1.460 | 1.51E-50 | NEG | NEG |
| *ZMIZ1* | chr10 | 79240241 | Astrocyte 1 | 171456 | 4.19 | 0.019 | intron | DEGs, DAGs | -1.210 | 7.98E-52 | NEG | NEG |
| *PITPNM2* | chr12 | 123059510 | Astrocyte 1 | 50668 | 4.30 | 0.019 | intron | DEGs, DAGs | -1.207 | 6.27E-56 | NEG | NEG |
| *SLC39A11* | chr17 | 72592555 | Astrocyte 1 | 499883 | 2.06 | 0.020 | Intergenic | DEGs, DAGs | -1.460 | 1.51E-50 | NEG | NEG |
| *FBXO2* | chr1 | 11479998 | Astrocyte 1 | 174434 | 3.30 | 0.020 | Intergenic | DEGs, DAGs | -1.338 | 1.24E-53 | NEG | NEG |
| *ZNF710* | chr15 | 90082235 | Astrocyte 1 | 81093 | 2.72 | 0.020 | Intergenic | DEGs, DAGs | -1.266 | 5.14E-76 | NEG | NEG |
| *TMEM184B* | chr22 | 38213772 | Astrocyte 1 | 59012 | 9.97 | 0.020 | Intergenic | DEGs, DAGs | -1.358 | 2.93E-66 | NEG | NEG |
| *H3F3B* | chr17 | 75779730 | Astrocyte 1 | -201 | 2.47 | 0.022 | promoter-TSS | DEGs, DAGs | -1.209 | 2.25E-49 | NEG | NEG |
| *NEK6* | chr9 | 124343043 | Astrocyte 1 | 85330 | 3.04 | 0.022 | intron | DEGs, DAGs | -1.406 | 2.80E-71 | NEG | NEG |
| *COL5A3* | chr19 | 9967005 | Astrocyte 1 | 43216 | 9.93 | 0.023 | intron | DEGs, DAGs | 1.581 | 7.92E-160 | NEG | NEG |
| *CACNA2D3* | chr3 | 54870252 | Astrocyte 1 | 747836 | 12.71 | 0.023 | intron | DEGs, DAGs | 1.523 | 8.67E-65 | NEG | NEG |
| *SLC16A9* | chr10 | 59622277 | Astrocyte 1 | 87552 | 2.73 | 0.023 | Intergenic | DEGs, DAGs | -1.209 | 7.10E-39 | NEG | NEG |
| *KLF2* | chr19 | 16460392 | Astrocyte 1 | 135825 | 3.23 | 0.024 | Intergenic | DEGs, DAGs | -1.391 | 2.76E-85 | NEG | NEG |
| *GNG7* | chr19 | 2578435 | Astrocyte 1 | 124024 | 5.74 | 0.024 | intron | DEGs, DAGs | -1.468 | 1.72E-60 | NEG | NEG |
| *RXRA* | chr9 | 134354442 | Astrocyte 1 | 28112 | 6.28 | 0.024 | intron | DEGs, DAGs | -1.215 | 5.85E-67 | NEG | NEG |
| *LHX2* | chr9 | 124110256 | Astrocyte 1 | 98896 | 4.71 | 0.024 | Intergenic | DEGs, DAGs | -1.389 | 7.39E-66 | NEG | NEG |
| *SDK2* | chr17 | 73356668 | Astrocyte 1 | 287171 | 2.96 | 0.024 | intron | DEGs, DAGs | -1.245 | 4.12E-68 | NEG | NEG |
| *EYA2* | chr20 | 47317139 | Astrocyte 1 | 422765 | 5.79 | 0.024 | Intergenic | DEGs, DAGs | -1.296 | 1.28E-29 | NEG | NEG |
| *KLF15* | chr3 | 126208618 | Astrocyte 1 | 148574 | 4.36 | 0.024 | Intergenic | DEGs, DAGs | -1.209 | 1.08E-55 | NEG | NEG |
| *RXRA* | chr9 | 134385086 | Astrocyte 1 | 58756 | 2.58 | 0.024 | intron | DEGs, DAGs | -1.215 | 5.85E-67 | NEG | NEG |
| *FBXO2* | chr1 | 11652863 | Astrocyte 1 | 1569 | 4.68 | 0.024 | promoter-TSS | DEGs, DAGs | -1.338 | 1.24E-53 | NEG | NEG |
| *SORBS1* | chr10 | 95435294 | Astrocyte 1 | 125834 | 4.39 | 0.024 | intron | DEGs, DAGs | -1.281 | 2.99E-44 | NEG | NEG |
| *TOM1L2* | chr17 | 17877952 | Astrocyte 1 | 94195 | 6.09 | 0.025 | intron | DEGs, DAGs | -1.408 | 5.09E-74 | NEG | NEG |
| *CAB39L* | chr13 | 49303565 | Astrocyte 1 | 109936 | 3.17 | 0.025 | Intergenic | DEGs, DAGs | -1.468 | 1.22E-135 | NEG | NEG |
| *GADD45B* | chr19 | 2611132 | Astrocyte 1 | 135255 | 4.77 | 0.025 | Intergenic | DEGs, DAGs | -1.226 | 1.00E-23 | NEG | NEG |
| *RFX2* | chr19 | 6087797 | Astrocyte 1 | 22487 | 6.24 | 0.025 | intron | DEGs, DAGs | -1.328 | 3.49E-49 | NEG | NEG |

| | | | | | | | | | | | | |
|---|---|---|---|---|---|---|---|---|---|---|---|---|
| *MINK1* | chr17 | 4881079 | Astrocyte 1 | 47941 | 10.88 | 0.025 | intron | DEGs, DAGs | -1.273 | 2.25E-54 | NEG | NEG |
| *CUX1* | chr7 | 101875029 | Astrocyte 1 | 59268 | 5.03 | 0.026 | intron | DEGs, DAGs | -1.271 | 8.56E-65 | NEG | NEG |
| *CSRP1* | chr1 | 201494211 | Astrocyte 1 | 14995 | 4.37 | 0.026 | intron | DEGs, DAGs | -1.335 | 4.90E-71 | NEG | NEG |
| *COL27A1* | chr9 | 114250612 | Astrocyte 1 | 95302 | 7.70 | 0.026 | intron | DEGs, DAGs | -1.357 | 1.23E-46 | NEG | NEG |
| *CD44* | chr11 | 35346145 | Astrocyte 1 | 207214 | 2.92 | 0.026 | Intergenic | DEGs, DAGs | -1.315 | 2.26E-16 | NEG | NEG |
| *STX8* | chr17 | 9292103 | Astrocyte 1 | 283883 | 3.45 | 0.026 | intron | DEGs, DAGs | -1.215 | 3.59E-63 | NEG | NEG |
| *GSE1* | chr16 | 85602009 | Astrocyte 1 | -10957 | 2.86 | 0.027 | intron | DEGs, DAGs | -1.237 | 3.56E-37 | NEG | NEG |
| *MAN1C1* | chr1 | 25652130 | Astrocyte 1 | 34912 | 2.52 | 0.027 | intron | DEGs, DAGs | -1.418 | 8.03E-37 | NEG | NEG |
| *SLC39A11* | chr17 | 72550325 | Astrocyte 1 | 542113 | 16.72 | 0.028 | Intergenic | DEGs, DAGs | -1.460 | 1.51E-50 | NEG | NEG |
| *SH3RF3* | chr2 | 109386318 | Astrocyte 1 | 257220 | 2.20 | 0.028 | intron | DEGs, DAGs | -1.237 | 1.77E-60 | NEG | NEG |
| *PPM1F* | chr22 | 21937164 | Astrocyte 1 | 15423 | 3.99 | 0.028 | intron | DEGs, DAGs | -1.205 | 1.27E-39 | NEG | NEG |
| *PPM1F* | chr22 | 21937164 | Astrocyte 1 | 15423 | 3.99 | 0.028 | exon | DEGs, DAGs | -1.205 | 1.27E-39 | NEG | NEG |
| *HIF3A* | chr19 | 46316266 | Astrocyte 1 | 19470 | 8.78 | 0.028 | intron | DEGs, DAGs | 1.412 | 6.22E-116 | NEG | NEG |
| *MRAS* | chr3 | 138359412 | Astrocyte 1 | 11965 | 2.23 | 0.028 | intron | DEGs, DAGs | -1.451 | 3.05E-64 | NEG | NEG |
| *C11orf49* | chr11 | 47154666 | Astrocyte 1 | 218138 | 3.59 | 0.028 | intron | DEGs, DAGs | -1.253 | 8.97E-81 | NEG | NEG |
| *C11orf49* | chr11 | 47154666 | Astrocyte 1 | 218138 | 3.59 | 0.028 | exon | DEGs, DAGs | -1.253 | 8.97E-81 | NEG | NEG |
| *TACC2* | chr10 | 122192068 | Astrocyte 1 | 203124 | 2.98 | 0.028 | intron | DEGs, DAGs | -1.234 | 4.39E-29 | NEG | NEG |
| *TACC2* | chr10 | 122192068 | Astrocyte 1 | 203124 | 2.98 | 0.028 | exon | DEGs, DAGs | -1.234 | 4.39E-29 | NEG | NEG |
| *MARK4* | chr19 | 45309908 | Astrocyte 1 | 58900 | 5.62 | 0.028 | Intergenic | DEGs, DAGs | -1.366 | 3.18E-67 | NEG | NEG |
| *PLCH1* | chr3 | 155356545 | Astrocyte 1 | 319244 | 2.23 | 0.029 | Intergenic | DEGs, DAGs | -1.212 | 1.87E-36 | NEG | NEG |
| *ZNF710* | chr15 | 90087654 | Astrocyte 1 | 86512 | 4.27 | 0.029 | Intergenic | DEGs, DAGs | -1.266 | 5.14E-76 | NEG | NEG |
| *SLC39A11* | chr17 | 72122617 | Astrocyte 1 | 969821 | 3.15 | 0.030 | Intergenic | DEGs, DAGs | -1.460 | 1.51E-50 | NEG | NEG |
| *ZSWIM6* | chr5 | 61292836 | Astrocyte 1 | -39187 | 2.68 | 0.030 | Intergenic | DEGs, DAGs | 1.296 | 5.38E-64 | NEG | NEG |
| *NEK6* | chr9 | 124392971 | Astrocyte 1 | 135258 | 3.99 | 0.030 | Intergenic | DEGs, DAGs | -1.406 | 2.80E-71 | NEG | NEG |
| *ALDH7A1* | chr5 | 126450892 | Astrocyte 1 | 144276 | 3.19 | 0.030 | Intergenic | DEGs, DAGs | -1.237 | 5.40E-60 | NEG | NEG |
| *ZMIZ1* | chr10 | 79248500 | Astrocyte 1 | 179715 | 2.21 | 0.031 | intron | DEGs, DAGs | -1.210 | 7.98E-52 | NEG | NEG |
| *VEGFA* | chr6 | 43972399 | Astrocyte 1 | 201942 | 3.25 | 0.031 | Intergenic | DEGs, DAGs | 1.284 | 1.68E-130 | NEG | NEG |
| *C11orf49* | chr11 | 47065909 | Astrocyte 1 | 129381 | 13.47 | 0.031 | intron | DEGs, DAGs | -1.253 | 8.97E-81 | NEG | NEG |
| *VEGFA* | chr6 | 43973354 | Astrocyte 1 | 202897 | 10.63 | 0.031 | Intergenic | DEGs, DAGs | 1.284 | 1.68E-130 | NEG | NEG |
| *CLIP2* | chr7 | 74379882 | Astrocyte 1 | 90657 | 2.34 | 0.031 | intron | DEGs, DAGs | -1.528 | 1.83E-58 | NEG | NEG |
| *CLIP2* | chr7 | 74379882 | Astrocyte 1 | 90657 | 2.34 | 0.031 | exon | DEGs, DAGs | -1.528 | 1.83E-58 | NEG | NEG |
| *NEK6* | chr9 | 124364062 | Astrocyte 1 | 106349 | 14.33 | 0.031 | Intergenic | DEGs, DAGs | -1.406 | 2.80E-71 | NEG | NEG |
| *AKAP1* | chr17 | 57092633 | Astrocyte 1 | 7166 | 2.02 | 0.032 | intron | DEGs, DAGs | 1.258 | 1.88E-59 | NEG | NEG |
| *CDK14* | chr7 | 91194752 | Astrocyte 1 | 598505 | 2.22 | 0.032 | intron | DEGs, DAGs | 1.253 | 7.33E-63 | NEG | NEG |

| | | | | | | | | | | | | |
|---|---|---|---|---|---|---|---|---|---|---|---|---|
| *MINK1* | chr17 | 4881915 | Astrocyte 1 | 48777 | 2.88 | 0.032 | intron | DEGs, DAGs | -1.273 | 2.25E-54 | NEG | NEG |
| *COBL* | chr7 | 52088557 | Astrocyte 1 | -771989 | 2.61 | 0.033 | Intergenic | DEGs, DAGs | -1.220 | 1.19E-60 | NEG | NEG |
| *ADRA1A* | chr8 | 26863491 | Astrocyte 1 | 1664 | 6.36 | 0.033 | promoter-TSS | DEGs, DAGs | 1.276 | 9.55E-62 | NEG | NEG |
| *GSE1* | chr16 | 85365958 | Astrocyte 1 | -247008 | 3.57 | 0.033 | intron | DEGs, DAGs | -1.237 | 3.56E-37 | NEG | NEG |
| *EYA2* | chr20 | 47319297 | Astrocyte 1 | 424923 | 2.22 | 0.033 | Intergenic | DEGs, DAGs | -1.296 | 1.28E-29 | NEG | NEG |
| *SLC39A11* | chr17 | 72416246 | Astrocyte 1 | 676192 | 2.01 | 0.034 | Intergenic | DEGs, DAGs | -1.460 | 1.51E-50 | NEG | NEG |
| *MMD2* | chr7 | 4915848 | Astrocyte 1 | 43114 | 4.50 | 0.034 | intron | DEGs, DAGs | 1.266 | 7.90E-70 | NEG | NEG |
| *GRIN2C* | chr17 | 74800687 | Astrocyte 1 | 58938 | 8.66 | 0.034 | Intergenic | DEGs, DAGs | -1.220 | 1.95E-92 | NEG | NEG |
| *SH3D19* | chr4 | 151251552 | Astrocyte 1 | -25294 | 2.40 | 0.034 | intron | DEGs, DAGs | -1.236 | 2.97E-19 | NEG | NEG |
| *KLF15* | chr3 | 126213000 | Astrocyte 1 | 144192 | 2.00 | 0.035 | Intergenic | DEGs, DAGs | -1.209 | 1.08E-55 | NEG | NEG |
| *KLF15* | chr3 | 126341652 | Astrocyte 1 | 15540 | 2.70 | 0.035 | Intergenic | DEGs, DAGs | -1.209 | 1.08E-55 | NEG | NEG |
| *ZMIZ1* | chr10 | 79165758 | Astrocyte 1 | 96973 | 2.81 | 0.035 | intron | DEGs, DAGs | -1.210 | 7.98E-52 | NEG | NEG |
| *DNMT3A* | chr2 | 25256938 | Astrocyte 1 | 85402 | 3.25 | 0.035 | intron | DEGs, DAGs | -1.415 | 3.81E-61 | NEG | NEG |
| *CUX1* | chr7 | 101900122 | Astrocyte 1 | 84361 | 3.22 | 0.035 | intron | DEGs, DAGs | -1.271 | 8.56E-65 | NEG | NEG |
| *SDK1* | chr7 | 3892689 | Astrocyte 1 | 591491 | -2.74 | 0.035 | intron | DEGs, DAGs | -1.255 | 4.71E-22 | NEG | NEG |
| *LHX2* | chr9 | 124223400 | Astrocyte 1 | 212040 | 3.32 | 0.037 | Intergenic | DEGs, DAGs | -1.389 | 7.39E-66 | NEG | NEG |
| *CTBP2* | chr10 | 125038600 | Astrocyte 1 | -10960 | 13.75 | 0.038 | exon | DEGs, DAGs | -1.249 | 4.58E-57 | NEG | NEG |
| *CTBP2* | chr10 | 125038600 | Astrocyte 1 | -10960 | 13.75 | 0.038 | intron | DEGs, DAGs | -1.249 | 4.58E-57 | NEG | NEG |
| *SLC39A11* | chr17 | 72586997 | Astrocyte 1 | 505441 | 11.82 | 0.038 | Intergenic | DEGs, DAGs | -1.460 | 1.51E-50 | NEG | NEG |
| *ZBTB16* | chr11 | 114064017 | Astrocyte 1 | 4674 | 14.24 | 0.038 | exon | DEGs, DAGs | -1.313 | 1.27E-83 | NEG | NEG |
| *ZBTB16* | chr11 | 114064017 | Astrocyte 1 | 4674 | 14.24 | 0.038 | intron | DEGs, DAGs | -1.313 | 1.27E-83 | NEG | NEG |
| *CHMP6* | chr17 | 81023714 | Astrocyte 1 | 32123 | 8.61 | 0.038 | Intergenic | DEGs, DAGs | -1.335 | 5.05E-55 | NEG | NEG |
| *ADGRG1* | chr16 | 57647519 | Astrocyte 1 | 27771 | 8.76 | 0.039 | intron | DEGs, DAGs | 1.338 | 1.56E-100 | NEG | NEG |
| *MERTK* | chr2 | 111986364 | Astrocyte 1 | 88135 | 7.02 | 0.039 | intron | DEGs, DAGs | -1.216 | 5.10E-54 | NEG | NEG |
| *ZBTB16* | chr11 | 114182371 | Astrocyte 1 | 123028 | 7.07 | 0.039 | intron | DEGs, DAGs | -1.313 | 1.27E-83 | NEG | NEG |
| *PIK3C2B* | chr1 | 204473218 | Astrocyte 1 | 16956 | 5.63 | 0.039 | intron | DEGs, DAGs | -1.203 | 1.30E-27 | NEG | NEG |
| *CHST11* | chr12 | 104709874 | Astrocyte 1 | 253151 | 14.13 | 0.039 | intron | DEGs, DAGs | -1.280 | 4.85E-42 | NEG | NEG |
| *ZMIZ1* | chr10 | 79254756 | Astrocyte 1 | 185971 | 3.16 | 0.040 | intron | DEGs, DAGs | -1.210 | 7.98E-52 | NEG | NEG |
| *CTBP2* | chr10 | 125048018 | Astrocyte 1 | -20378 | 3.19 | 0.041 | intron | DEGs, DAGs | -1.249 | 4.58E-57 | NEG | NEG |
| *TMSB10* | chr2 | 84926919 | Astrocyte 1 | 21544 | 2.29 | 0.041 | Intergenic | DEGs, DAGs | -1.263 | 1.03E-15 | NEG | NEG |
| *LHX2* | chr9 | 124013712 | Astrocyte 1 | 2352 | 4.45 | 0.042 | intron | DEGs, DAGs | -1.389 | 7.39E-66 | NEG | NEG |
| *SORBS1* | chr10 | 95335799 | Astrocyte 1 | 225329 | 26.94 | 0.043 | intron | DEGs, DAGs | -1.281 | 2.99E-44 | NEG | NEG |
| *B4GALNT4* | chr11 | 379231 | Astrocyte 1 | 9677 | 18.52 | 0.043 | intron | DEGs, DAGs | 1.316 | 7.27E-80 | NEG | NEG |
| *C11orf49* | chr11 | 47163045 | Astrocyte 1 | 226517 | 13.86 | 0.043 | intron | DEGs, DAGs | -1.253 | 8.97E-81 | NEG | NEG |

| | | | | | | | | | | | | |
|---|---|---|---|---|---|---|---|---|---|---|---|---|
| *PITPNM2* | chr12 | 123052013 | Astrocyte 1 | 58165 | 10.77 | 0.043 | intron | DEGs, DAGs | -1.207 | 6.27E-56 | NEG | NEG |
| *GADD45B* | chr19 | 2676051 | Astrocyte 1 | 200174 | 11.81 | 0.043 | Intergenic | DEGs, DAGs | -1.226 | 1.00E-23 | NEG | NEG |
| *GADD45B* | chr19 | 2676911 | Astrocyte 1 | 201034 | 12.50 | 0.043 | Intergenic | DEGs, DAGs | -1.226 | 1.00E-23 | NEG | NEG |
| *DYSF* | chr2 | 71871517 | Astrocyte 1 | 405065 | 15.47 | 0.043 | Intergenic | DEGs, DAGs | -1.253 | 6.88E-21 | NEG | NEG |
| *GALNT13* | chr2 | 154374952 | Astrocyte 1 | 430704 | 12.61 | 0.043 | intron | DEGs, DAGs | -1.206 | 4.82E-06 | NEG | NEG |
| *DST* | chr6 | 56552089 | Astrocyte 1 | 299577 | 22.51 | 0.043 | exon | DEGs, DAGs | 1.296 | 1.70E-82 | NEG | NEG |
| *DST* | chr6 | 56552089 | Astrocyte 1 | 299577 | 22.51 | 0.043 | intron | DEGs, DAGs | 1.296 | 1.70E-82 | NEG | NEG |
| *MAP7* | chr6 | 136306435 | Astrocyte 1 | 219528 | 4.04 | 0.043 | Intergenic | DEGs, DAGs | -1.263 | 2.75E-120 | NEG | NEG |
| *RAPGEF5* | chr7 | 22313647 | Astrocyte 1 | 43017 | -26.12 | 0.043 | intron | DEGs, DAGs | -1.280 | 1.75E-19 | NEG | NEG |
| *CAMK2B* | chr7 | 44219158 | Astrocyte 1 | 106090 | 14.15 | 0.043 | intron | DEGs, DAGs | 1.304 | 8.54E-91 | NEG | NEG |
| *CAMK2B* | chr7 | 44219158 | Astrocyte 1 | 106090 | 14.15 | 0.043 | exon | DEGs, DAGs | 1.304 | 8.54E-91 | NEG | NEG |
| *DAPK1* | chr9 | 87656415 | Astrocyte 1 | 158784 | 7.83 | 0.043 | intron | DEGs, DAGs | -1.259 | 1.60E-62 | NEG | NEG |
| *KREMEN1* | chr22 | 29154345 | Astrocyte 1 | 81477 | 5.86 | 0.043 | intron | DEGs, DAGs | -1.250 | 7.80E-24 | NEG | NEG |
| *KREMEN1* | chr22 | 29154345 | Astrocyte 1 | 81477 | 5.86 | 0.043 | exon | DEGs, DAGs | -1.250 | 7.80E-24 | NEG | NEG |
| *CERK* | chr22 | 46636901 | Astrocyte 1 | 101110 | 3.83 | 0.043 | Intergenic | DEGs, DAGs | -1.220 | 2.67E-27 | NEG | NEG |
| *GSE1* | chr16 | 85395601 | Astrocyte 1 | -217365 | 6.47 | 0.044 | intron | DEGs, DAGs | -1.237 | 3.56E-37 | NEG | NEG |
| *SIPA1L3* | chr19 | 38095623 | Astrocyte 1 | 188645 | 9.38 | 0.044 | intron | DEGs, DAGs | -1.326 | 9.22E-75 | NEG | NEG |
| *KAZN* | chr1 | 14930548 | Astrocyte 1 | 332094 | 10.33 | 0.044 | intron | DEGs, DAGs | -1.370 | 3.41E-43 | NEG | NEG |
| *KLF6* | chr10 | 3538892 | Astrocyte 1 | 246133 | -11.66 | 0.044 | Intergenic | DEGs, DAGs | -1.210 | 3.25E-17 | NEG | NEG |
| *SLC39A11* | chr17 | 72415506 | Astrocyte 1 | 676932 | 3.61 | 0.044 | Intergenic | DEGs, DAGs | -1.460 | 1.51E-50 | NEG | NEG |
| *RGS3* | chr9 | 113566609 | Astrocyte 1 | 122128 | 2.59 | 0.044 | intron | DEGs, DAGs | -1.240 | 4.11E-53 | NEG | NEG |
| *GNG7* | chr19 | 2495219 | Astrocyte 1 | 207240 | 5.27 | 0.045 | Intergenic | DEGs, DAGs | -1.468 | 1.72E-60 | NEG | NEG |
| *RAPGEF3* | chr12 | 47736721 | Astrocyte 1 | 21674 | 2.02 | 0.045 | exon | DEGs, DAGs | 1.288 | 3.27E-113 | NEG | NEG |
| *RAPGEF3* | chr12 | 47736721 | Astrocyte 1 | 21674 | 2.02 | 0.045 | intron | DEGs, DAGs | 1.288 | 3.27E-113 | NEG | NEG |
| *GNG7* | chr19 | 2543535 | Astrocyte 1 | 158924 | 2.63 | 0.045 | intron | DEGs, DAGs | -1.468 | 1.72E-60 | NEG | NEG |
| *RXRA* | chr9 | 134484264 | Astrocyte 1 | 157934 | 2.35 | 0.045 | Intergenic | DEGs, DAGs | -1.215 | 5.85E-67 | NEG | NEG |
| *SORBS1* | chr10 | 95457525 | Astrocyte 1 | 103603 | 2.61 | 0.045 | intron | DEGs, DAGs | -1.281 | 2.99E-44 | NEG | NEG |
| *AK5* | chr1 | 77489988 | Astrocyte 1 | 208187 | 2.58 | 0.045 | intron | DEGs, DAGs | -1.296 | 2.34E-26 | NEG | NEG |
| *CKB* | chr14 | 103393914 | Astrocyte 1 | 128947 | 6.56 | 0.046 | Intergenic | DEGs, DAGs | 1.492 | 5.93E-140 | NEG | NEG |
| *TRNAU1AP* | chr1 | 28582208 | Astrocyte 1 | 29373 | 5.13 | 0.046 | Intergenic | DEGs, DAGs | -1.203 | 6.07E-35 | NEG | NEG |
| *GSE1* | chr16 | 85656617 | Astrocyte 1 | 43651 | 2.29 | 0.046 | intron | DEGs, DAGs | -1.237 | 3.56E-37 | NEG | NEG |
| *GSE1* | chr16 | 85656617 | Astrocyte 1 | 43651 | 2.29 | 0.046 | exon | DEGs, DAGs | -1.237 | 3.56E-37 | NEG | NEG |
| *AJAP1* | chr1 | 4408980 | Astrocyte 1 | -245815 | 3.91 | 0.046 | Intergenic | DEGs, DAGs | -1.255 | 3.35E-59 | NEG | NEG |
| *ADCY9* | chr16 | 3939012 | Astrocyte 1 | 176923 | 4.81 | 0.046 | Intergenic | DEGs, DAGs | -1.217 | 2.34E-48 | NEG | NEG |

| | | | | | | | | | | | | |
|---|---|---|---|---|---|---|---|---|---|---|---|---|
| *TEAD1* | chr11 | 12845625 | Astrocyte 1 | 171284 | 3.75 | 0.046 | intron | DEGs, DAGs | -1.229 | 5.49E-75 | NEG | NEG |
| *RGS9* | chr17 | 65516258 | Astrocyte 1 | 379034 | 4.18 | 0.046 | Intergenic | DEGs, DAGs | -1.199 | 1.97E-12 | NEG | NEG |
| *EYA2* | chr20 | 47318494 | Astrocyte 1 | 424120 | 2.51 | 0.046 | Intergenic | DEGs, DAGs | -1.296 | 1.28E-29 | NEG | NEG |
| *CUX1* | chr7 | 102260788 | Astrocyte 1 | 445027 | 2.02 | 0.046 | intron | DEGs, DAGs | -1.271 | 8.56E-65 | NEG | NEG |
| *CORO2B* | chr15 | 68775671 | Astrocyte 1 | 196952 | 2.86 | 0.046 | Intergenic | DEGs, DAGs | -1.243 | 4.68E-56 | NEG | NEG |
| *AJAP1* | chr1 | 4512609 | Astrocyte 1 | -142186 | 3.74 | 0.046 | Intergenic | DEGs, DAGs | -1.255 | 3.35E-59 | NEG | NEG |
| *OAF* | chr11 | 120187185 | Astrocyte 1 | -23332 | 2.67 | 0.046 | Intergenic | DEGs, DAGs | -1.447 | 7.94E-70 | NEG | NEG |
| *DPYSL5* | chr2 | 26961671 | Astrocyte 1 | 113820 | 3.65 | 0.047 | Intergenic | DEGs, DAGs | -1.289 | 1.46E-24 | NEG | NEG |
| *ADARB2* | chr10 | 1110001 | Astrocyte 1 | 627225 | 2.19 | 0.047 | Intergenic | DEGs, DAGs | -1.244 | 4.07E-37 | NEG | NEG |
| *LHX2* | chr9 | 124018110 | Astrocyte 1 | 6750 | 2.50 | 0.049 | intron | DEGs, DAGs | -1.389 | 7.39E-66 | NEG | NEG |
| *FGFR3* | chr4 | 1775533 | Astrocyte 1 | -17510 | 3.30 | 0.000 | Distal enhancer | DEGs, DAGs | 1.330 | 1.07E-88 | FGFR3/GH04J001757 | eQTLs,eRNA_co-expression |
| *FGFR3* | chr4 | 1775533 | Astrocyte 1 | -17510 | 3.30 | 0.000 | Distal enhancer | DEGs, DAGs | 1.330 | 1.07E-88 | FGFR3/GH04J001764 | eRNA_co-expression |
| *FGFR3* | chr4 | 1775533 | Astrocyte 1 | -17510 | 3.30 | 0.000 | Distal enhancer | DEGs, DAGs | 1.330 | 1.07E-88 | FGFR3/GH04J001775 | eQTLs |
| *FGFR3* | chr4 | 1775533 | Astrocyte 1 | -17510 | 3.30 | 0.000 | Distal enhancer | DEGs, DAGs | 1.330 | 1.07E-88 | FGFR3/GH04J001784 | eQTLs |
| *FGFR3* | chr4 | 1775533 | Astrocyte 1 | -17510 | 3.30 | 0.000 | Distal enhancer | DEGs, DAGs | 1.330 | 1.07E-88 | FGFR3/GH04J001792 | eQTLs |
| *FGFR3* | chr4 | 1784490 | Astrocyte 1 | -8553 | 2.39 | 0.019 | Distal enhancer | DEGs, DAGs | 1.330 | 1.07E-88 | FGFR3/GH04J001757 | eQTLs,eRNA_co-expression |
| *FGFR3* | chr4 | 1784490 | Astrocyte 1 | -8553 | 2.39 | 0.019 | Distal enhancer | DEGs, DAGs | 1.330 | 1.07E-88 | FGFR3/GH04J001764 | eRNA_co-expression |
| *FGFR3* | chr4 | 1784490 | Astrocyte 1 | -8553 | 2.39 | 0.019 | Distal enhancer | DEGs, DAGs | 1.330 | 1.07E-88 | FGFR3/GH04J001775 | eQTLs |
| *FGFR3* | chr4 | 1784490 | Astrocyte 1 | -8553 | 2.39 | 0.019 | Distal enhancer | DEGs, DAGs | 1.330 | 1.07E-88 | FGFR3/GH04J001784 | eQTLs |
| *FGFR3* | chr4 | 1784490 | Astrocyte 1 | -8553 | 2.39 | 0.019 | Distal enhancer | DEGs, DAGs | 1.330 | 1.07E-88 | FGFR3/GH04J001792 | eQTLs |
| *FGFR3* | chr4 | 1763318 | Astrocyte 1 | -29725 | 2.55 | 0.036 | Distal enhancer | DEGs, DAGs | 1.330 | 1.07E-88 | FGFR3/GH04J001757 | eQTLs,eRNA_co-expression |
| *FGFR3* | chr4 | 1763318 | Astrocyte 1 | -29725 | 2.55 | 0.036 | Distal enhancer | DEGs, DAGs | 1.330 | 1.07E-88 | FGFR3/GH04J001764 | eRNA_co-expression |
| *FGFR3* | chr4 | 1763318 | Astrocyte 1 | -29725 | 2.55 | 0.036 | Distal enhancer | DEGs, DAGs | 1.330 | 1.07E-88 | FGFR3/GH04J001775 | eQTLs |
| *FGFR3* | chr4 | 1763318 | Astrocyte 1 | -29725 | 2.55 | 0.036 | Distal enhancer | DEGs, DAGs | 1.330 | 1.07E-88 | FGFR3/GH04J001784 | eQTLs |
| *FGFR3* | chr4 | 1763318 | Astrocyte 1 | -29725 | 2.55 | 0.036 | Distal enhancer | DEGs, DAGs | 1.330 | 1.07E-88 | FGFR3/GH04J001792 | eQTLs |
| *ADCY2* | chr5 | 7392333 | Astrocyte 1 | -3625 | 3.79 | 0.016 | Distal enhancer | DEGs, DAGs | -1.409 | 1.92E-74 | ADCY2/GH05J007391 | eQTLs |
| *ADCY2* | chr5 | 7392333 | Astrocyte 1 | -3625 | 3.79 | 0.016 | Distal enhancer | DEGs, DAGs | -1.409 | 1.92E-74 | ADCY2/GH05J007393 | eQTLs,eRNA_co-expression |
| *ADCY2* | chr5 | 7392333 | Astrocyte 1 | -3625 | 3.79 | 0.016 | Distal enhancer | DEGs, DAGs | -1.409 | 1.92E-74 | ADCY2/GH05J007802 | C-HiC |
| *ADCY2* | chr5 | 7392333 | Astrocyte 1 | -3625 | 3.79 | 0.016 | Distal enhancer | DEGs, DAGs | -1.409 | 1.92E-74 | ADCY2/GH05J007421 | eQTLs,C-HiC |
| *ADCY2* | chr5 | 7392333 | Astrocyte 1 | -3625 | 3.79 | 0.016 | Distal enhancer | DEGs, DAGs | -1.409 | 1.92E-74 | ADCY2/GH05J007419 | C-HiC |
| *HSPA1A* | chr6 | 31821171 | Astrocyte 1 | 5878 | 4.96 | 0.044 | Distal enhancer | DEGs, DAGs | 1.636 | 2.51E-30 | HSPA1A/GH06J031733 | C-HiC,eRNA_co-expression |
| *HSPA1A* | chr6 | 31821171 | Astrocyte 1 | 5878 | 4.96 | 0.044 | Distal enhancer | DEGs, DAGs | 1.636 | 2.51E-30 | HSPA1A/GH06J031813 | eRNA_co-expression |
| *COX19* | chr7 | 977740 | Astrocyte 1 | -2391 | 2.13 | 0.003 | Distal enhancer | DEGs, DAGs | -1.302 | 2.99E-47 | COX19/GH07J000906 | eRNA_co-expression |

| | | | | | | | | | | | | |
|---|---|---|---|---|---|---|---|---|---|---|---|---|
| *COX19* | chr7 | 977740 | Astrocyte 1 | -2391 | 2.13 | 0.003 | Distal enhancer | DEGs, DAGs | -1.302 | 2.99E-47 | COX19/GH07J000921 | eRNA_co-expression |
| *COX19* | chr7 | 977740 | Astrocyte 1 | -2391 | 2.13 | 0.003 | Distal enhancer | DEGs, DAGs | -1.302 | 2.99E-47 | COX19/GH07J000962 | eQTLs |
| *COX19* | chr7 | 977740 | Astrocyte 1 | -2391 | 2.13 | 0.003 | Distal enhancer | DEGs, DAGs | -1.302 | 2.99E-47 | COX19/GH07J000971 | eQTLs |
| *COX19* | chr7 | 977740 | Astrocyte 1 | -2391 | 2.13 | 0.003 | Distal enhancer | DEGs, DAGs | -1.302 | 2.99E-47 | COX19/GH07J000981 | eQTLs |
| *COX19* | chr7 | 977740 | Astrocyte 1 | -2391 | 2.13 | 0.003 | Distal enhancer | DEGs, DAGs | -1.302 | 2.99E-47 | COX19/GH07J001058 | eQTLs,eRNA_co-expression |
| *COX19* | chr7 | 977740 | Astrocyte 1 | -2391 | 2.13 | 0.003 | Distal enhancer | DEGs, DAGs | -1.302 | 2.99E-47 | COX19/GH07J001022 | eQTLs,eRNA_co-expression |
| *PTPRD* | chr9 | 10612526 | Astrocyte 1 | -53 | 4.86 | 0.015 | Distal enhancer | DEGs, DAGs | -1.300 | 1.20E-68 | PTPRD/GH09J010612 | eQTLs |
| *NEK6* | chr9 | 124223400 | Astrocyte 1 | -33956 | 3.32 | 0.037 | Distal enhancer | DEGs, DAGs | -1.406 | 2.80E-71 | NEK6/GH09J124207 | eQTLs |
| *NEK6* | chr9 | 124223400 | Astrocyte 1 | -33956 | 3.32 | 0.037 | Distal enhancer | DEGs, DAGs | -1.406 | 2.80E-71 | NEK6/GH09J124210 | eQTLs |
| *NEK6* | chr9 | 124223400 | Astrocyte 1 | -33956 | 3.32 | 0.037 | Distal enhancer | DEGs, DAGs | -1.406 | 2.80E-71 | NEK6/GH09J124212 | eQTLs |
| *NEK6* | chr9 | 124223400 | Astrocyte 1 | -33956 | 3.32 | 0.037 | Distal enhancer | DEGs, DAGs | -1.406 | 2.80E-71 | NEK6/GH09J124213 | eQTLs |
| *NEK6* | chr9 | 124223400 | Astrocyte 1 | -33956 | 3.32 | 0.037 | Distal enhancer | DEGs, DAGs | -1.406 | 2.80E-71 | NEK6/GH09J124215 | eQTLs,eRNA_co-expression |
| *NEK6* | chr9 | 124223400 | Astrocyte 1 | -33956 | 3.32 | 0.037 | Distal enhancer | DEGs, DAGs | -1.406 | 2.80E-71 | NEK6/GH09J124220 | eQTLs,C-HiC |
| *NEK6* | chr9 | 124223400 | Astrocyte 1 | -33956 | 3.32 | 0.037 | Distal enhancer | DEGs, DAGs | -1.406 | 2.80E-71 | NEK6/GH09J124238 | eQTLs,eRNA_co-expression |
| *NEK6* | chr9 | 124223400 | Astrocyte 1 | -33956 | 3.32 | 0.037 | Distal enhancer | DEGs, DAGs | -1.406 | 2.80E-71 | NEK6/GH09J124242 | eQTLs |
| *NEK6* | chr9 | 124223400 | Astrocyte 1 | -33956 | 3.32 | 0.037 | Distal enhancer | DEGs, DAGs | -1.406 | 2.80E-71 | NEK6/GH09J124255 | eQTLs |
| *NEK6* | chr9 | 124223400 | Astrocyte 1 | -33956 | 3.32 | 0.037 | Distal enhancer | DEGs, DAGs | -1.406 | 2.80E-71 | NEK6/GH09J124256 | eQTLs |
| *NEK6* | chr9 | 124223400 | Astrocyte 1 | -33956 | 3.32 | 0.037 | Distal enhancer | DEGs, DAGs | -1.406 | 2.80E-71 | NEK6/GH09J124360 | eQTLs |
| *NEK6* | chr9 | 124223400 | Astrocyte 1 | -33956 | 3.32 | 0.037 | Distal enhancer | DEGs, DAGs | -1.406 | 2.80E-71 | NEK6/GH09J124340 | eQTLs |
| *NEK6* | chr9 | 124223400 | Astrocyte 1 | -33956 | 3.32 | 0.037 | Distal enhancer | DEGs, DAGs | -1.406 | 2.80E-71 | NEK6/GH09J124308 | eQTLs |
| *NEK6* | chr9 | 124223400 | Astrocyte 1 | -33956 | 3.32 | 0.037 | Distal enhancer | DEGs, DAGs | -1.406 | 2.80E-71 | NEK6/GH09J124305 | eQTLs |
| *NEK6* | chr9 | 124223400 | Astrocyte 1 | -33956 | 3.32 | 0.037 | Distal enhancer | DEGs, DAGs | -1.406 | 2.80E-71 | NEK6/GH09J124303 | eQTLs |
| *NEK6* | chr9 | 124223400 | Astrocyte 1 | -33956 | 3.32 | 0.037 | Distal enhancer | DEGs, DAGs | -1.406 | 2.80E-71 | NEK6/GH09J124277 | eQTLs,eRNA_co-expression |
| *NEK6* | chr9 | 124223400 | Astrocyte 1 | -33956 | 3.32 | 0.037 | Distal enhancer | DEGs, DAGs | -1.406 | 2.80E-71 | NEK6/GH09J124270 | eQTLs |
| *CAMK2G* | chr10 | 73886004 | Astrocyte 1 | -11663 | 3.33 | 0.019 | Distal enhancer | DEGs, DAGs | -1.448 | 2.18E-82 | CAMK2G/GH10J073809 | eQTLs,C-HiC |
| *CAMK2G* | chr10 | 73886004 | Astrocyte 1 | -11663 | 3.33 | 0.019 | Distal enhancer | DEGs, DAGs | -1.448 | 2.18E-82 | CAMK2G/GH10J073819 | C-HiC |
| *CAMK2G* | chr10 | 73886004 | Astrocyte 1 | -11663 | 3.33 | 0.019 | Distal enhancer | DEGs, DAGs | -1.448 | 2.18E-82 | CAMK2G/GH10J073836 | eQTLs,C-HiC |
| *CAMK2G* | chr10 | 73886004 | Astrocyte 1 | -11663 | 3.33 | 0.019 | Distal enhancer | DEGs, DAGs | -1.448 | 2.18E-82 | CAMK2G/GH10J073838 | C-HiC |
| *CAMK2G* | chr10 | 73886004 | Astrocyte 1 | -11663 | 3.33 | 0.019 | Distal enhancer | DEGs, DAGs | -1.448 | 2.18E-82 | CAMK2G/GH10J073841 | eQTLs |
| *CAMK2G* | chr10 | 73886004 | Astrocyte 1 | -11663 | 3.33 | 0.019 | Distal enhancer | DEGs, DAGs | -1.448 | 2.18E-82 | CAMK2G/GH10J073848 | C-HiC |
| *CAMK2G* | chr10 | 73886004 | Astrocyte 1 | -11663 | 3.33 | 0.019 | Distal enhancer | DEGs, DAGs | -1.448 | 2.18E-82 | CAMK2G/GH10J073850 | eQTLs,C-HiC |
| *CAMK2G* | chr10 | 73886004 | Astrocyte 1 | -11663 | 3.33 | 0.019 | Distal enhancer | DEGs, DAGs | -1.448 | 2.18E-82 | CAMK2G/GH10J073863 | eQTLs,eRNA_co-expression |

| | | | | | | | | | | | | |
|---|---|---|---|---|---|---|---|---|---|---|---|---|
| *CAMK2G* | chr10 | 73886004 | Astrocyte 1 | -11663 | 3.33 | 0.019 | Distal enhancer | DEGs, DAGs | -1.448 | 2.18E-82 | CAMK2G/GH10J073869 | eQTLs |
| *CAMK2G* | chr10 | 73886004 | Astrocyte 1 | -11663 | 3.33 | 0.019 | Distal enhancer | DEGs, DAGs | -1.448 | 2.18E-82 | CAMK2G/GH10J073886 | eQTLs |
| *SLC3A2* | chr11 | 62889982 | Astrocyte 1 | 8507 | 2.51 | 0.012 | Distal enhancer | DEGs, DAGs | 1.394 | 1.07E-83 | SLC3A2/GH11J062852 | eRNA_co-expression |
| *SLC3A2* | chr11 | 62889982 | Astrocyte 1 | 8507 | 2.51 | 0.012 | Distal enhancer | DEGs, DAGs | 1.394 | 1.07E-83 | SLC3A2/GH11J062887 | C-HiC |
| *SLC3A2* | chr11 | 62889982 | Astrocyte 1 | 8507 | 2.51 | 0.012 | Distal enhancer | DEGs, DAGs | 1.394 | 1.07E-83 | SLC3A2/GH11J062878 | C-HiC |
| *P2RX7* | chr12 | 121128846 | Astrocyte 1 | -3732 | 3.40 | 0.000 | Distal enhancer | DEGs, DAGs | -1.296 | 5.84E-27 | P2RX7/GH12J121059 | C-HiC,eRNA_co-expression |
| *P2RX7* | chr12 | 121128846 | Astrocyte 1 | -3732 | 3.40 | 0.000 | Distal enhancer | DEGs, DAGs | -1.296 | 5.84E-27 | P2RX7/GH12J121127 | eRNA_co-expression |
| *P2RX7* | chr12 | 121128846 | Astrocyte 1 | -3732 | 3.40 | 0.000 | Distal enhancer | DEGs, DAGs | -1.296 | 5.84E-27 | P2RX7/GH12J121132 | eRNA_co-expression |
| *P2RX7* | chr12 | 121128846 | Astrocyte 1 | -3732 | 3.40 | 0.000 | Distal enhancer | DEGs, DAGs | -1.296 | 5.84E-27 | P2RX7/GH12J121196 | C-HiC,eRNA_co-expression |
| *P2RX7* | chr12 | 121128846 | Astrocyte 1 | -3732 | 3.40 | 0.000 | Distal enhancer | DEGs, DAGs | -1.296 | 5.84E-27 | P2RX7/GH12J121185 | C-HiC |
| *P2RX7* | chr12 | 121128846 | Astrocyte 1 | -3732 | 3.40 | 0.000 | Distal enhancer | DEGs, DAGs | -1.296 | 5.84E-27 | P2RX7/GH12J121184 | C-HiC |
| *P2RX7* | chr12 | 121128846 | Astrocyte 1 | -3732 | 3.40 | 0.000 | Distal enhancer | DEGs, DAGs | -1.296 | 5.84E-27 | P2RX7/GH12J121165 | C-HiC |
| *P2RX7* | chr12 | 121128846 | Astrocyte 1 | -3732 | 3.40 | 0.000 | Distal enhancer | DEGs, DAGs | -1.296 | 5.84E-27 | P2RX7/GH12J121159 | C-HiC |
| *P2RX7* | chr12 | 121128846 | Astrocyte 1 | -3732 | 3.40 | 0.000 | Distal enhancer | DEGs, DAGs | -1.296 | 5.84E-27 | P2RX7/GH12J121143 | eQTLs,eRNA_co-expression |
| *CKB* | chr14 | 103518486 | Astrocyte 1 | 4375 | 4.54 | 0.004 | Distal enhancer | DEGs, DAGs | 1.492 | 5.93E-140 | CKB/GH14J103332 | eQTLs,TF_co-expression |
| *CKB* | chr14 | 103518486 | Astrocyte 1 | 4375 | 4.54 | 0.004 | Distal enhancer | DEGs, DAGs | 1.492 | 5.93E-140 | CKB/GH14J103517 | eQTLs |
| *CKB* | chr14 | 103518486 | Astrocyte 1 | 4375 | 4.54 | 0.004 | Distal enhancer | DEGs, DAGs | 1.492 | 5.93E-140 | CKB/GH14J103549 | eQTLs,eRNA_co-expression |
| *ANKDD1A* | chr15 | 64893927 | Astrocyte 1 | -17725 | 2.82 | 0.005 | Distal enhancer | DEGs, DAGs | 1.614 | 2.33E-151 | ANKDD1A/GH15J064840 | eQTLs,C-HiC |
| *ANKDD1A* | chr15 | 64893927 | Astrocyte 1 | -17725 | 2.82 | 0.005 | Distal enhancer | DEGs, DAGs | 1.614 | 2.33E-151 | ANKDD1A/GH15J064891 | eQTLs |
| *ANKDD1A* | chr15 | 64893927 | Astrocyte 1 | -17725 | 2.82 | 0.005 | Distal enhancer | DEGs, DAGs | 1.614 | 2.33E-151 | ANKDD1A/GH15J064903 | eQTLs |
| *STARD3* | chr17 | 39632547 | Astrocyte 1 | -4268 | 2.49 | 0.000 | Distal enhancer | DEGs, DAGs | -1.204 | 5.71E-37 | STARD3/GH17J039449 | eQTLs,TF_co-expression |
| *STARD3* | chr17 | 39632547 | Astrocyte 1 | -4268 | 2.49 | 0.000 | Distal enhancer | DEGs, DAGs | -1.204 | 5.71E-37 | STARD3/GH17J039460 | eQTLs,TF_co-expression |
| *STARD3* | chr17 | 39632547 | Astrocyte 1 | -4268 | 2.49 | 0.000 | Distal enhancer | DEGs, DAGs | -1.204 | 5.71E-37 | STARD3/GH17J039626 | eQTLs |
| *STARD3* | chr17 | 39632547 | Astrocyte 1 | -4268 | 2.49 | 0.000 | Distal enhancer | DEGs, DAGs | -1.204 | 5.71E-37 | STARD3/GH17J039631 | eQTLs |
| *STARD3* | chr17 | 39632547 | Astrocyte 1 | -4268 | 2.49 | 0.000 | Distal enhancer | DEGs, DAGs | -1.204 | 5.71E-37 | STARD3/GH17J039635 | eQTLs |
| *STARD3* | chr17 | 39632547 | Astrocyte 1 | -4268 | 2.49 | 0.000 | Distal enhancer | DEGs, DAGs | -1.204 | 5.71E-37 | STARD3/GH17J039658 | TF_co-expression |
| *STARD3* | chr17 | 39632547 | Astrocyte 1 | -4268 | 2.49 | 0.000 | Distal enhancer | DEGs, DAGs | -1.204 | 5.71E-37 | STARD3/GH17J039651 | eQTLs |
| *STARD3* | chr17 | 39632547 | Astrocyte 1 | -4268 | 2.49 | 0.000 | Distal enhancer | DEGs, DAGs | -1.204 | 5.71E-37 | STARD3/GH17J039649 | TF_co-expression |
| *STARD3* | chr17 | 39632547 | Astrocyte 1 | -4268 | 2.49 | 0.000 | Distal enhancer | DEGs, DAGs | -1.204 | 5.71E-37 | STARD3/GH17J039644 | eQTLs |
| *STARD3* | chr17 | 39632547 | Astrocyte 1 | -4268 | 2.49 | 0.000 | Distal enhancer | DEGs, DAGs | -1.204 | 5.71E-37 | STARD3/GH17J039641 | TF_co-expression |
| *STARD3* | chr17 | 39632547 | Astrocyte 1 | -4268 | 2.49 | 0.000 | Distal enhancer | DEGs, DAGs | -1.204 | 5.71E-37 | STARD3/GH17J039639 | eQTLs |
| *STARD3* | chr17 | 39633093 | Astrocyte 1 | -3722 | 3.68 | 0.005 | Distal enhancer | DEGs, DAGs | -1.204 | 5.71E-37 | STARD3/GH17J039449 | eQTLs,TF_co-expression |

| | | | | | | | | | | | | |
|---|---|---|---|---|---|---|---|---|---|---|---|---|
| *STARD3* | chr17 | 39633093 | Astrocyte 1 | -3722 | 3.68 | 0.005 | Distal enhancer | DEGs, DAGs | -1.204 | 5.71E-37 | STARD3/GH17J039460 | eQTLs,TF_co-expression |
| *STARD3* | chr17 | 39633093 | Astrocyte 1 | -3722 | 3.68 | 0.005 | Distal enhancer | DEGs, DAGs | -1.204 | 5.71E-37 | STARD3/GH17J039626 | eQTLs |
| *STARD3* | chr17 | 39633093 | Astrocyte 1 | -3722 | 3.68 | 0.005 | Distal enhancer | DEGs, DAGs | -1.204 | 5.71E-37 | STARD3/GH17J039631 | eQTLs |
| *STARD3* | chr17 | 39633093 | Astrocyte 1 | -3722 | 3.68 | 0.005 | Distal enhancer | DEGs, DAGs | -1.204 | 5.71E-37 | STARD3/GH17J039635 | eQTLs |
| *STARD3* | chr17 | 39633093 | Astrocyte 1 | -3722 | 3.68 | 0.005 | Distal enhancer | DEGs, DAGs | -1.204 | 5.71E-37 | STARD3/GH17J039658 | TF_co-expression |
| *STARD3* | chr17 | 39633093 | Astrocyte 1 | -3722 | 3.68 | 0.005 | Distal enhancer | DEGs, DAGs | -1.204 | 5.71E-37 | STARD3/GH17J039651 | eQTLs |
| *STARD3* | chr17 | 39633093 | Astrocyte 1 | -3722 | 3.68 | 0.005 | Distal enhancer | DEGs, DAGs | -1.204 | 5.71E-37 | STARD3/GH17J039649 | TF_co-expression |
| *STARD3* | chr17 | 39633093 | Astrocyte 1 | -3722 | 3.68 | 0.005 | Distal enhancer | DEGs, DAGs | -1.204 | 5.71E-37 | STARD3/GH17J039644 | eQTLs |
| *STARD3* | chr17 | 39633093 | Astrocyte 1 | -3722 | 3.68 | 0.005 | Distal enhancer | DEGs, DAGs | -1.204 | 5.71E-37 | STARD3/GH17J039641 | TF_co-expression |
| *STARD3* | chr17 | 39633093 | Astrocyte 1 | -3722 | 3.68 | 0.005 | Distal enhancer | DEGs, DAGs | -1.204 | 5.71E-37 | STARD3/GH17J039639 | eQTLs |
| *STARD3* | chr17 | 39634309 | Astrocyte 1 | -2506 | 21.18 | 0.006 | Distal enhancer | DEGs, DAGs | -1.204 | 5.71E-37 | STARD3/GH17J039449 | eQTLs,TF_co-expression |
| *STARD3* | chr17 | 39634309 | Astrocyte 1 | -2506 | 21.18 | 0.006 | Distal enhancer | DEGs, DAGs | -1.204 | 5.71E-37 | STARD3/GH17J039460 | eQTLs,TF_co-expression |
| *STARD3* | chr17 | 39634309 | Astrocyte 1 | -2506 | 21.18 | 0.006 | Distal enhancer | DEGs, DAGs | -1.204 | 5.71E-37 | STARD3/GH17J039626 | eQTLs |
| *STARD3* | chr17 | 39634309 | Astrocyte 1 | -2506 | 21.18 | 0.006 | Distal enhancer | DEGs, DAGs | -1.204 | 5.71E-37 | STARD3/GH17J039631 | eQTLs |
| *STARD3* | chr17 | 39634309 | Astrocyte 1 | -2506 | 21.18 | 0.006 | Distal enhancer | DEGs, DAGs | -1.204 | 5.71E-37 | STARD3/GH17J039635 | eQTLs |
| *STARD3* | chr17 | 39634309 | Astrocyte 1 | -2506 | 21.18 | 0.006 | Distal enhancer | DEGs, DAGs | -1.204 | 5.71E-37 | STARD3/GH17J039658 | TF_co-expression |
| *STARD3* | chr17 | 39634309 | Astrocyte 1 | -2506 | 21.18 | 0.006 | Distal enhancer | DEGs, DAGs | -1.204 | 5.71E-37 | STARD3/GH17J039651 | eQTLs |
| *STARD3* | chr17 | 39634309 | Astrocyte 1 | -2506 | 21.18 | 0.006 | Distal enhancer | DEGs, DAGs | -1.204 | 5.71E-37 | STARD3/GH17J039649 | TF_co-expression |
| *STARD3* | chr17 | 39634309 | Astrocyte 1 | -2506 | 21.18 | 0.006 | Distal enhancer | DEGs, DAGs | -1.204 | 5.71E-37 | STARD3/GH17J039644 | eQTLs |
| *STARD3* | chr17 | 39634309 | Astrocyte 1 | -2506 | 21.18 | 0.006 | Distal enhancer | DEGs, DAGs | -1.204 | 5.71E-37 | STARD3/GH17J039641 | TF_co-expression |
| *STARD3* | chr17 | 39634309 | Astrocyte 1 | -2506 | 21.18 | 0.006 | Distal enhancer | DEGs, DAGs | -1.204 | 5.71E-37 | STARD3/GH17J039639 | eQTLs |
| *GRIN2C* | chr17 | 74840331 | Astrocyte 1 | 19294 | 5.00 | 0.015 | Distal enhancer | DEGs, DAGs | -1.220 | 1.95E-92 | GRIN2C/GH17J074807 | C-HiC,eRNA_co-expression |
| *GRIN2C* | chr17 | 74840331 | Astrocyte 1 | 19294 | 5.00 | 0.015 | Distal enhancer | DEGs, DAGs | -1.220 | 1.95E-92 | GRIN2C/GH17J074813 | C-HiC,eRNA_co-expression |
| *GRIN2C* | chr17 | 74840331 | Astrocyte 1 | 19294 | 5.00 | 0.015 | Distal enhancer | DEGs, DAGs | -1.220 | 1.95E-92 | GRIN2C/GH17J074834 | eQTLs,C-HiC |
| *GRIN2C* | chr17 | 74840331 | Astrocyte 1 | 19294 | 5.00 | 0.015 | Distal enhancer | DEGs, DAGs | -1.220 | 1.95E-92 | GRIN2C/GH17J074840 | eQTLs |
| *GRIN2C* | chr17 | 74840331 | Astrocyte 1 | 19294 | 5.00 | 0.015 | Distal enhancer | DEGs, DAGs | -1.220 | 1.95E-92 | GRIN2C/GH17J074851 | eQTLs |
| *CHMP6* | chr17 | 80988896 | Astrocyte 1 | -2695 | 3.91 | 0.020 | Distal enhancer | DEGs, DAGs | -1.335 | 5.05E-55 | CHMP6/GH17J080889 | eQTLs,TF_co-expression |
| *CHMP6* | chr17 | 80988896 | Astrocyte 1 | -2695 | 3.91 | 0.020 | Distal enhancer | DEGs, DAGs | -1.335 | 5.05E-55 | CHMP6/GH17J080958 | eQTLs,C-HiC |
| *CHMP6* | chr17 | 80988896 | Astrocyte 1 | -2695 | 3.91 | 0.020 | Distal enhancer | DEGs, DAGs | -1.335 | 5.05E-55 | CHMP6/GH17J080968 | eQTLs,C-HiC |
| *CHMP6* | chr17 | 80988896 | Astrocyte 1 | -2695 | 3.91 | 0.020 | Distal enhancer | DEGs, DAGs | -1.335 | 5.05E-55 | CHMP6/GH17J080980 | eQTLs,TF_co-expression |
| *CHMP6* | chr17 | 80988896 | Astrocyte 1 | -2695 | 3.91 | 0.020 | Distal enhancer | DEGs, DAGs | -1.335 | 5.05E-55 | CHMP6/GH17J080986 | eQTLs |
| *CHMP6* | chr17 | 80988896 | Astrocyte 1 | -2695 | 3.91 | 0.020 | Distal enhancer | DEGs, DAGs | -1.335 | 5.05E-55 | CHMP6/GH17J080988 | eQTLs |

| | | | | | | | | | | | | |
|---|---|---|---|---|---|---|---|---|---|---|---|---|
| *CHMP6* | chr17 | 80988896 | Astrocyte 1 | -2695 | 3.91 | 0.020 | Distal enhancer | DEGs, DAGs | -1.335 | 5.05E-55 | CHMP6/GH17J080991 | eQTLs,TF_co-expression |
| *CHMP6* | chr17 | 80988896 | Astrocyte 1 | -2695 | 3.91 | 0.020 | Distal enhancer | DEGs, DAGs | -1.335 | 5.05E-55 | CHMP6/GH17J081017 | eQTLs,TF_co-expression |
| *FTL* | chr19 | 48967449 | Astrocyte 1 | 2398 | 6.32 | 0.033 | Distal enhancer | DEGs, DAGs | 1.463 | 2.53E-34 | FTL/GH19J048958 | eRNA_co-expression |
| *FTL* | chr19 | 48968008 | Astrocyte 1 | 2957 | 5.54 | 0.035 | Distal enhancer | DEGs, DAGs | 1.463 | 2.53E-34 | FTL/GH19J048958 | eRNA_co-expression |
| *ZFP36* | chr19 | 39403546 | Astrocyte 1 | -3017 | 2.30 | 0.046 | Distal enhancer | DEGs, DAGs | -1.425 | 9.33E-50 | ZFP36/GH19J039397 | eRNA_co-expression,TF_co-expression |
| *ZFP36* | chr19 | 39403546 | Astrocyte 1 | -3017 | 2.30 | 0.046 | Distal enhancer | DEGs, DAGs | -1.425 | 9.33E-50 | ZFP36/GH19J039552 | C-HiC,eRNA_co-expression |
| *ERCC6L2* | chr9 | 95515744 | Astrocyte 1 | -359707 | 12.09 | 0.000 | Distal enhancer | DEGs, DAGs | 1.299 | 2.89E-83 | ERCC6L2/GH09J095503 | eRNA_co-expression,TF_co-expression |
| *ERCC6L2* | chr9 | 95515744 | Astrocyte 1 | -359707 | 12.09 | 0.000 | Distal enhancer | DEGs, DAGs | 1.299 | 2.89E-83 | ERCC6L2/GH09J095874 | eQTLs,TF_co-expression |
| *ERCC6L2* | chr9 | 95515744 | Astrocyte 1 | -359707 | 12.09 | 0.000 | Distal enhancer | DEGs, DAGs | 1.299 | 2.89E-83 | ERCC6L2/GH09J096072 | TF_co-expression |
| *ERCC6L2* | chr9 | 95515744 | Astrocyte 1 | -359707 | 12.09 | 0.000 | Distal enhancer | DEGs, DAGs | 1.299 | 2.89E-83 | ERCC6L2/GH09J096061 | eQTLs |
| *ERCC6L2* | chr9 | 95515744 | Astrocyte 1 | -359707 | 12.09 | 0.000 | Distal enhancer | DEGs, DAGs | 1.299 | 2.89E-83 | ERCC6L2/GH09J096055 | eQTLs |
| *ERCC6L2* | chr9 | 95515744 | Astrocyte 1 | -359707 | 12.09 | 0.000 | Distal enhancer | DEGs, DAGs | 1.299 | 2.89E-83 | ERCC6L2/GH09J096047 | eQTLs |
| *ERCC6L2* | chr9 | 95515744 | Astrocyte 1 | -359707 | 12.09 | 0.000 | Distal enhancer | DEGs, DAGs | 1.299 | 2.89E-83 | ERCC6L2/GH09J096040 | eQTLs |
| *ERCC6L2* | chr9 | 95515744 | Astrocyte 1 | -359707 | 12.09 | 0.000 | Distal enhancer | DEGs, DAGs | 1.299 | 2.89E-83 | ERCC6L2/GH09J096030 | eQTLs |
| *ERCC6L2* | chr9 | 95515744 | Astrocyte 1 | -359707 | 12.09 | 0.000 | Distal enhancer | DEGs, DAGs | 1.299 | 2.89E-83 | ERCC6L2/GH09J096025 | eQTLs,eRNA_co-expression |
| *ERCC6L2* | chr9 | 95515744 | Astrocyte 1 | -359707 | 12.09 | 0.000 | Distal enhancer | DEGs, DAGs | 1.299 | 2.89E-83 | ERCC6L2/GH09J096020 | eQTLs |
| *ERCC6L2* | chr9 | 95515744 | Astrocyte 1 | -359707 | 12.09 | 0.000 | Distal enhancer | DEGs, DAGs | 1.299 | 2.89E-83 | ERCC6L2/GH09J096015 | eQTLs |
| *ERCC6L2* | chr9 | 95515744 | Astrocyte 1 | -359707 | 12.09 | 0.000 | Distal enhancer | DEGs, DAGs | 1.299 | 2.89E-83 | ERCC6L2/GH09J096008 | eQTLs,eRNA_co-expression |
| *ERCC6L2* | chr9 | 95515744 | Astrocyte 1 | -359707 | 12.09 | 0.000 | Distal enhancer | DEGs, DAGs | 1.299 | 2.89E-83 | ERCC6L2/GH09J096006 | eQTLs |
| *ERCC6L2* | chr9 | 95515744 | Astrocyte 1 | -359707 | 12.09 | 0.000 | Distal enhancer | DEGs, DAGs | 1.299 | 2.89E-83 | ERCC6L2/GH09J095980 | eQTLs |
| *ERCC6L2* | chr9 | 95515744 | Astrocyte 1 | -359707 | 12.09 | 0.000 | Distal enhancer | DEGs, DAGs | 1.299 | 2.89E-83 | ERCC6L2/GH09J095887 | eQTLs |
| *ERCC6L2* | chr9 | 95515744 | Astrocyte 1 | -359707 | 12.09 | 0.000 | Distal enhancer | DEGs, DAGs | 1.299 | 2.89E-83 | ERCC6L2/GH09J095879 | eQTLs,eRNA_co-expression |
| *FGFR3* | chr4 | 1775533 | Astrocyte 1 | -17529 | 3.30 | 0.000 | Distal enhancer | DEGs, DAGs | 1.330 | 1.07E-88 | FGFR3/GH04J001757 | eQTLs,eRNA_co-expression |
| *FGFR3* | chr4 | 1775533 | Astrocyte 1 | -17529 | 3.30 | 0.000 | Distal enhancer | DEGs, DAGs | 1.330 | 1.07E-88 | FGFR3/GH04J001764 | eRNA_co-expression |
| *FGFR3* | chr4 | 1775533 | Astrocyte 1 | -17529 | 3.30 | 0.000 | Distal enhancer | DEGs, DAGs | 1.330 | 1.07E-88 | FGFR3/GH04J001775 | eQTLs |
| *FGFR3* | chr4 | 1775533 | Astrocyte 1 | -17529 | 3.30 | 0.000 | Distal enhancer | DEGs, DAGs | 1.330 | 1.07E-88 | FGFR3/GH04J001784 | eQTLs |
| *FGFR3* | chr4 | 1775533 | Astrocyte 1 | -17529 | 3.30 | 0.000 | Distal enhancer | DEGs, DAGs | 1.330 | 1.07E-88 | FGFR3/GH04J001792 | eQTLs |
| *JUNB* | chr19 | 12782256 | Astrocyte 1 | -8990 | 3.96 | 0.004 | Distal enhancer | DEGs, DAGs | -1.512 | 1.07E-62 | JUNB/GH19J012781 | eRNA_co-expression |
| *JUNB* | chr19 | 12782256 | Astrocyte 1 | -8990 | 3.96 | 0.004 | Distal enhancer | DEGs, DAGs | -1.512 | 1.07E-62 | JUNB/GH19J013160 | eQTLs,eRNA_co-expression |
| *JUNB* | chr19 | 12782256 | Astrocyte 1 | -8990 | 3.96 | 0.004 | Distal enhancer | DEGs, DAGs | -1.512 | 1.07E-62 | JUNB/GH19J013148 | eQTLs,eRNA_co-expression |
| *DCAKD* | chr17 | 45093607 | Astrocyte 1 | -32748 | 3.52 | 0.007 | Distal enhancer | DEGs, DAGs | -1.244 | 4.92E-39 | DCAKD/GH17J045020 | eQTLs,C-HiC |
| *DCAKD* | chr17 | 45093607 | Astrocyte 1 | -32748 | 3.52 | 0.007 | Distal enhancer | DEGs, DAGs | -1.244 | 4.92E-39 | DCAKD/GH17J045029 | eQTLs,C-HiC |

| | | | | | | | | | | | | |
|---|---|---|---|---|---|---|---|---|---|---|---|---|
| *DCAKD* | chr17 | 45093607 | Astrocyte 1 | -32748 | 3.52 | 0.007 | Distal enhancer | DEGs, DAGs | -1.244 | 4.92E-39 | DCAKD/GH17J045038 | eQTLs |
| *DCAKD* | chr17 | 45093607 | Astrocyte 1 | -32748 | 3.52 | 0.007 | Distal enhancer | DEGs, DAGs | -1.244 | 4.92E-39 | DCAKD/GH17J045042 | eQTLs |
| *DCAKD* | chr17 | 45093607 | Astrocyte 1 | -32748 | 3.52 | 0.007 | Distal enhancer | DEGs, DAGs | -1.244 | 4.92E-39 | DCAKD/GH17J045049 | eQTLs |
| *DCAKD* | chr17 | 45093607 | Astrocyte 1 | -32748 | 3.52 | 0.007 | Distal enhancer | DEGs, DAGs | -1.244 | 4.92E-39 | DCAKD/GH17J045059 | eQTLs |
| *DCAKD* | chr17 | 45093607 | Astrocyte 1 | -32748 | 3.52 | 0.007 | Distal enhancer | DEGs, DAGs | -1.244 | 4.92E-39 | DCAKD/GH17J045168 | eQTLs,eRNA_co-expression |
| *DCAKD* | chr17 | 45093607 | Astrocyte 1 | -32748 | 3.52 | 0.007 | Distal enhancer | DEGs, DAGs | -1.244 | 4.92E-39 | DCAKD/GH17J045143 | eQTLs,TF_co-expression |
| *DCAKD* | chr17 | 45093607 | Astrocyte 1 | -32748 | 3.52 | 0.007 | Distal enhancer | DEGs, DAGs | -1.244 | 4.92E-39 | DCAKD/GH17J045089 | eQTLs,C-HiC |
| *TCF20* | chr22 | 42282712 | Astrocyte 1 | -67520 | 9.55 | 0.007 | Distal enhancer | DEGs, DAGs | -1.233 | 3.20E-41 | TCF20/GH22J042176 | C-HiC,eRNA_co-expression |
| *TCF20* | chr22 | 42282712 | Astrocyte 1 | -67520 | 9.55 | 0.007 | Distal enhancer | DEGs, DAGs | -1.233 | 3.20E-41 | TCF20/GH22J042268 | C-HiC |
| *TCF20* | chr22 | 42282712 | Astrocyte 1 | -67520 | 9.55 | 0.007 | Distal enhancer | DEGs, DAGs | -1.233 | 3.20E-41 | TCF20/GH22J042271 | C-HiC |
| *TCF20* | chr22 | 42282712 | Astrocyte 1 | -67520 | 9.55 | 0.007 | Distal enhancer | DEGs, DAGs | -1.233 | 3.20E-41 | TCF20/GH22J042273 | C-HiC |
| *TCF20* | chr22 | 42282712 | Astrocyte 1 | -67520 | 9.55 | 0.007 | Distal enhancer | DEGs, DAGs | -1.233 | 3.20E-41 | TCF20/GH22J042282 | C-HiC |
| *TCF20* | chr22 | 42282712 | Astrocyte 1 | -67520 | 9.55 | 0.007 | Distal enhancer | DEGs, DAGs | -1.233 | 3.20E-41 | TCF20/GH22J042298 | TF_co-expression |
| *TCF20* | chr22 | 42282712 | Astrocyte 1 | -67520 | 9.55 | 0.007 | Distal enhancer | DEGs, DAGs | -1.233 | 3.20E-41 | TCF20/GH22J042320 | C-HiC |
| *TCF20* | chr22 | 42282712 | Astrocyte 1 | -67520 | 9.55 | 0.007 | Distal enhancer | DEGs, DAGs | -1.233 | 3.20E-41 | TCF20/GH22J042444 | C-HiC,eRNA_co-expression |
| *TCF20* | chr22 | 42282712 | Astrocyte 1 | -67520 | 9.55 | 0.007 | Distal enhancer | DEGs, DAGs | -1.233 | 3.20E-41 | TCF20/GH22J042436 | C-HiC,eRNA_co-expression |
| *TCF20* | chr22 | 42282712 | Astrocyte 1 | -67520 | 9.55 | 0.007 | Distal enhancer | DEGs, DAGs | -1.233 | 3.20E-41 | TCF20/GH22J042364 | C-HiC,eRNA_co-expression |
| *CHMP6* | chr17 | 80959524 | Astrocyte 1 | -32067 | 2.26 | 0.008 | Distal enhancer | DEGs, DAGs | -1.335 | 5.05E-55 | CHMP6/GH17J080889 | eQTLs,TF_co-expression |
| *CHMP6* | chr17 | 80959524 | Astrocyte 1 | -32067 | 2.26 | 0.008 | Distal enhancer | DEGs, DAGs | -1.335 | 5.05E-55 | CHMP6/GH17J080958 | eQTLs,C-HiC |
| *CHMP6* | chr17 | 80959524 | Astrocyte 1 | -32067 | 2.26 | 0.008 | Distal enhancer | DEGs, DAGs | -1.335 | 5.05E-55 | CHMP6/GH17J080968 | eQTLs,C-HiC |
| *CHMP6* | chr17 | 80959524 | Astrocyte 1 | -32067 | 2.26 | 0.008 | Distal enhancer | DEGs, DAGs | -1.335 | 5.05E-55 | CHMP6/GH17J080980 | eQTLs,TF_co-expression |
| *CHMP6* | chr17 | 80959524 | Astrocyte 1 | -32067 | 2.26 | 0.008 | Distal enhancer | DEGs, DAGs | -1.335 | 5.05E-55 | CHMP6/GH17J080986 | eQTLs |
| *CHMP6* | chr17 | 80959524 | Astrocyte 1 | -32067 | 2.26 | 0.008 | Distal enhancer | DEGs, DAGs | -1.335 | 5.05E-55 | CHMP6/GH17J080988 | eQTLs |
| *CHMP6* | chr17 | 80959524 | Astrocyte 1 | -32067 | 2.26 | 0.008 | Distal enhancer | DEGs, DAGs | -1.335 | 5.05E-55 | CHMP6/GH17J080991 | eQTLs,TF_co-expression |
| *CHMP6* | chr17 | 80959524 | Astrocyte 1 | -32067 | 2.26 | 0.008 | Distal enhancer | DEGs, DAGs | -1.335 | 5.05E-55 | CHMP6/GH17J081017 | eQTLs,TF_co-expression |
| *HIF1A* | chr14 | 61574030 | Astrocyte 1 | -123342 | -5.19 | 0.008 | Distal enhancer | DEGs, DAGs | 1.296 | 4.92E-82 | HIF1A/GH14J061465 | C-HiC,eRNA_co-expression |
| *HIF1A* | chr14 | 61574030 | Astrocyte 1 | -123342 | -5.19 | 0.008 | Distal enhancer | DEGs, DAGs | 1.296 | 4.92E-82 | HIF1A/GH14J061468 | C-HiC,eRNA_co-expression |
| *HIF1A* | chr14 | 61574030 | Astrocyte 1 | -123342 | -5.19 | 0.008 | Distal enhancer | DEGs, DAGs | 1.296 | 4.92E-82 | HIF1A/GH14J061489 | C-HiC,eRNA_co-expression |
| *HIF1A* | chr14 | 61574030 | Astrocyte 1 | -123342 | -5.19 | 0.008 | Distal enhancer | DEGs, DAGs | 1.296 | 4.92E-82 | HIF1A/GH14J061510 | C-HiC,eRNA_co-expression |
| *HIF1A* | chr14 | 61574030 | Astrocyte 1 | -123342 | -5.19 | 0.008 | Distal enhancer | DEGs, DAGs | 1.296 | 4.92E-82 | HIF1A/GH14J061518 | C-HiC,eRNA_co-expression |
| *HIF1A* | chr14 | 61574030 | Astrocyte 1 | -123342 | -5.19 | 0.008 | Distal enhancer | DEGs, DAGs | 1.296 | 4.92E-82 | HIF1A/GH14J061530 | eQTLs,C-HiC |
| *HIF1A* | chr14 | 61574030 | Astrocyte 1 | -123342 | -5.19 | 0.008 | Distal enhancer | DEGs, DAGs | 1.296 | 4.92E-82 | HIF1A/GH14J061551 | C-HiC,eRNA_co-expression |

| | | | | | | | | | | | | |
|---|---|---|---|---|---|---|---|---|---|---|---|---|
| *HIF1A* | chr14 | 61574030 | Astrocyte 1 | -123342 | -5.19 | 0.008 | Distal enhancer | DEGs, DAGs | 1.296 | 4.92E-82 | HIF1A/GH14J061580 | C-HiC,eRNA_co-expression |
| *HIF1A* | chr14 | 61574030 | Astrocyte 1 | -123342 | -5.19 | 0.008 | Distal enhancer | DEGs, DAGs | 1.296 | 4.92E-82 | HIF1A/GH14J061593 | C-HiC,eRNA_co-expression |
| *HIF1A* | chr14 | 61574030 | Astrocyte 1 | -123342 | -5.19 | 0.008 | Distal enhancer | DEGs, DAGs | 1.296 | 4.92E-82 | HIF1A/GH14J061599 | C-HiC,eRNA_co-expression |
| *HIF1A* | chr14 | 61574030 | Astrocyte 1 | -123342 | -5.19 | 0.008 | Distal enhancer | DEGs, DAGs | 1.296 | 4.92E-82 | HIF1A/GH14J061601 | C-HiC,eRNA_co-expression |
| *HIF1A* | chr14 | 61574030 | Astrocyte 1 | -123342 | -5.19 | 0.008 | Distal enhancer | DEGs, DAGs | 1.296 | 4.92E-82 | HIF1A/GH14J061619 | eQTLs,C-HiC |
| *HIF1A* | chr14 | 61574030 | Astrocyte 1 | -123342 | -5.19 | 0.008 | Distal enhancer | DEGs, DAGs | 1.296 | 4.92E-82 | HIF1A/GH14J061641 | eQTLs,eRNA_co-expression |
| *HIF1A* | chr14 | 61574030 | Astrocyte 1 | -123342 | -5.19 | 0.008 | Distal enhancer | DEGs, DAGs | 1.296 | 4.92E-82 | HIF1A/GH14J061660 | C-HiC,eRNA_co-expression |
| *HIF1A* | chr14 | 61574030 | Astrocyte 1 | -123342 | -5.19 | 0.008 | Distal enhancer | DEGs, DAGs | 1.296 | 4.92E-82 | HIF1A/GH14J061747 | eRNA_co-expression |
| *HIF1A* | chr14 | 61574030 | Astrocyte 1 | -123342 | -5.19 | 0.008 | Distal enhancer | DEGs, DAGs | 1.296 | 4.92E-82 | HIF1A/GH14J061717 | C-HiC |
| *LRMDA* | chr10 | 75409408 | Astrocyte 1 | -21795 | 18.77 | 0.009 | Distal enhancer | DEGs, DAGs | 1.707 | 2.56E-125 | LRMDA/GH10J075394 | eQTLs,C-HiC,eRNA_co-expression |
| *LRMDA* | chr10 | 75409408 | Astrocyte 1 | -21795 | 18.77 | 0.009 | Distal enhancer | DEGs, DAGs | 1.707 | 2.56E-125 | LRMDA/GH10J075427 | eQTLs |
| *LRMDA* | chr10 | 75409408 | Astrocyte 1 | -21795 | 18.77 | 0.009 | Distal enhancer | DEGs, DAGs | 1.707 | 2.56E-125 | LRMDA/GH10J076428 | eQTLs |
| *LRMDA* | chr10 | 75409408 | Astrocyte 1 | -21795 | 18.77 | 0.009 | Distal enhancer | DEGs, DAGs | 1.707 | 2.56E-125 | LRMDA/GH10J076396 | eQTLs |
| *LRMDA* | chr10 | 75409408 | Astrocyte 1 | -21795 | 18.77 | 0.009 | Distal enhancer | DEGs, DAGs | 1.707 | 2.56E-125 | LRMDA/GH10J076335 | eQTLs |
| *LRMDA* | chr10 | 75409408 | Astrocyte 1 | -21795 | 18.77 | 0.009 | Distal enhancer | DEGs, DAGs | 1.707 | 2.56E-125 | LRMDA/GH10J076306 | eQTLs |
| *LRMDA* | chr10 | 75409408 | Astrocyte 1 | -21795 | 18.77 | 0.009 | Distal enhancer | DEGs, DAGs | 1.707 | 2.56E-125 | LRMDA/GH10J076302 | eQTLs |
| *LRMDA* | chr10 | 75409408 | Astrocyte 1 | -21795 | 18.77 | 0.009 | Distal enhancer | DEGs, DAGs | 1.707 | 2.56E-125 | LRMDA/GH10J076033 | C-HiC,eRNA_co-expression |
| *LRMDA* | chr10 | 75409408 | Astrocyte 1 | -21795 | 18.77 | 0.009 | Distal enhancer | DEGs, DAGs | 1.707 | 2.56E-125 | LRMDA/GH10J075904 | C-HiC |
| *LRMDA* | chr10 | 75409408 | Astrocyte 1 | -21795 | 18.77 | 0.009 | Distal enhancer | DEGs, DAGs | 1.707 | 2.56E-125 | LRMDA/GH10J075788 | C-HiC |
| *LRMDA* | chr10 | 75409408 | Astrocyte 1 | -21795 | 18.77 | 0.009 | Distal enhancer | DEGs, DAGs | 1.707 | 2.56E-125 | LRMDA/GH10J075771 | C-HiC |
| *LRMDA* | chr10 | 75409408 | Astrocyte 1 | -21795 | 18.77 | 0.009 | Distal enhancer | DEGs, DAGs | 1.707 | 2.56E-125 | LRMDA/GH10J075764 | C-HiC |
| *LRMDA* | chr10 | 75409408 | Astrocyte 1 | -21795 | 18.77 | 0.009 | Distal enhancer | DEGs, DAGs | 1.707 | 2.56E-125 | LRMDA/GH10J075612 | eQTLs,C-HiC |
| *LRMDA* | chr10 | 75409408 | Astrocyte 1 | -21795 | 18.77 | 0.009 | Distal enhancer | DEGs, DAGs | 1.707 | 2.56E-125 | LRMDA/GH10J075610 | eQTLs,C-HiC |
| *LRMDA* | chr10 | 75409408 | Astrocyte 1 | -21795 | 18.77 | 0.009 | Distal enhancer | DEGs, DAGs | 1.707 | 2.56E-125 | LRMDA/GH10J075542 | C-HiC |
| *LRMDA* | chr10 | 75409408 | Astrocyte 1 | -21795 | 18.77 | 0.009 | Distal enhancer | DEGs, DAGs | 1.707 | 2.56E-125 | LRMDA/GH10J075543 | C-HiC |
| *LRMDA* | chr10 | 75409408 | Astrocyte 1 | -21795 | 18.77 | 0.009 | Distal enhancer | DEGs, DAGs | 1.707 | 2.56E-125 | LRMDA/GH10J075528 | eQTLs |
| *LRMDA* | chr10 | 75409408 | Astrocyte 1 | -21795 | 18.77 | 0.009 | Distal enhancer | DEGs, DAGs | 1.707 | 2.56E-125 | LRMDA/GH10J075507 | eQTLs |
| *LRMDA* | chr10 | 75409408 | Astrocyte 1 | -21795 | 18.77 | 0.009 | Distal enhancer | DEGs, DAGs | 1.707 | 2.56E-125 | LRMDA/GH10J075494 | eQTLs |
| *LRMDA* | chr10 | 75409408 | Astrocyte 1 | -21795 | 18.77 | 0.009 | Distal enhancer | DEGs, DAGs | 1.707 | 2.56E-125 | LRMDA/GH10J075470 | C-HiC |
| *LRMDA* | chr10 | 75409408 | Astrocyte 1 | -21795 | 18.77 | 0.009 | Distal enhancer | DEGs, DAGs | 1.707 | 2.56E-125 | LRMDA/GH10J075468 | eQTLs,C-HiC |
| *CHMP6* | chr17 | 80958791 | Astrocyte 1 | -32800 | 4.25 | 0.014 | Distal enhancer | DEGs, DAGs | -1.335 | 5.05E-55 | CHMP6/GH17J080889 | eQTLs,TF_co-expression |
| *CHMP6* | chr17 | 80958791 | Astrocyte 1 | -32800 | 4.25 | 0.014 | Distal enhancer | DEGs, DAGs | -1.335 | 5.05E-55 | CHMP6/GH17J080958 | eQTLs,C-HiC |

| | | | | | | | | | | | | |
|---|---|---|---|---|---|---|---|---|---|---|---|---|
| *CHMP6* | chr17 | 80958791 | Astrocyte 1 | -32800 | 4.25 | 0.014 | Distal enhancer | DEGs, DAGs | -1.335 | 5.05E-55 | CHMP6/GH17J080968 | eQTLs,C-HiC |
| *CHMP6* | chr17 | 80958791 | Astrocyte 1 | -32800 | 4.25 | 0.014 | Distal enhancer | DEGs, DAGs | -1.335 | 5.05E-55 | CHMP6/GH17J080980 | eQTLs,TF_co-expression |
| *CHMP6* | chr17 | 80958791 | Astrocyte 1 | -32800 | 4.25 | 0.014 | Distal enhancer | DEGs, DAGs | -1.335 | 5.05E-55 | CHMP6/GH17J080986 | eQTLs |
| *CHMP6* | chr17 | 80958791 | Astrocyte 1 | -32800 | 4.25 | 0.014 | Distal enhancer | DEGs, DAGs | -1.335 | 5.05E-55 | CHMP6/GH17J080988 | eQTLs |
| *CHMP6* | chr17 | 80958791 | Astrocyte 1 | -32800 | 4.25 | 0.014 | Distal enhancer | DEGs, DAGs | -1.335 | 5.05E-55 | CHMP6/GH17J080991 | eQTLs,TF_co-expression |
| *CHMP6* | chr17 | 80958791 | Astrocyte 1 | -32800 | 4.25 | 0.014 | Distal enhancer | DEGs, DAGs | -1.335 | 5.05E-55 | CHMP6/GH17J081017 | eQTLs,TF_co-expression |
| *TCF20* | chr22 | 42365156 | Astrocyte 1 | -149964 | -4.70 | 0.015 | Distal enhancer | DEGs, DAGs | -1.233 | 3.20E-41 | TCF20/GH22J042176 | C-HiC,eRNA_co-expression |
| *TCF20* | chr22 | 42365156 | Astrocyte 1 | -149964 | -4.70 | 0.015 | Distal enhancer | DEGs, DAGs | -1.233 | 3.20E-41 | TCF20/GH22J042268 | C-HiC |
| *TCF20* | chr22 | 42365156 | Astrocyte 1 | -149964 | -4.70 | 0.015 | Distal enhancer | DEGs, DAGs | -1.233 | 3.20E-41 | TCF20/GH22J042271 | C-HiC |
| *TCF20* | chr22 | 42365156 | Astrocyte 1 | -149964 | -4.70 | 0.015 | Distal enhancer | DEGs, DAGs | -1.233 | 3.20E-41 | TCF20/GH22J042273 | C-HiC |
| *TCF20* | chr22 | 42365156 | Astrocyte 1 | -149964 | -4.70 | 0.015 | Distal enhancer | DEGs, DAGs | -1.233 | 3.20E-41 | TCF20/GH22J042282 | C-HiC |
| *TCF20* | chr22 | 42365156 | Astrocyte 1 | -149964 | -4.70 | 0.015 | Distal enhancer | DEGs, DAGs | -1.233 | 3.20E-41 | TCF20/GH22J042298 | TF_co-expression |
| *TCF20* | chr22 | 42365156 | Astrocyte 1 | -149964 | -4.70 | 0.015 | Distal enhancer | DEGs, DAGs | -1.233 | 3.20E-41 | TCF20/GH22J042320 | C-HiC |
| *TCF20* | chr22 | 42365156 | Astrocyte 1 | -149964 | -4.70 | 0.015 | Distal enhancer | DEGs, DAGs | -1.233 | 3.20E-41 | TCF20/GH22J042444 | C-HiC,eRNA_co-expression |
| *TCF20* | chr22 | 42365156 | Astrocyte 1 | -149964 | -4.70 | 0.015 | Distal enhancer | DEGs, DAGs | -1.233 | 3.20E-41 | TCF20/GH22J042436 | C-HiC,eRNA_co-expression |
| *TCF20* | chr22 | 42365156 | Astrocyte 1 | -149964 | -4.70 | 0.015 | Distal enhancer | DEGs, DAGs | -1.233 | 3.20E-41 | TCF20/GH22J042364 | C-HiC,eRNA_co-expression |
| *PTPRD* | chr9 | 10612526 | Astrocyte 1 | -578986 | 4.86 | 0.015 | Distal enhancer | DEGs, DAGs | -1.300 | 1.20E-68 | PTPRD/GH09J010612 | eQTLs |
| *FGFR3* | chr4 | 1784490 | Astrocyte 1 | -8572 | 2.39 | 0.019 | Distal enhancer | DEGs, DAGs | 1.330 | 1.07E-88 | FGFR3/GH04J001757 | eQTLs,eRNA_co-expression,Distance |
| *FGFR3* | chr4 | 1784490 | Astrocyte 1 | -8572 | 2.39 | 0.019 | Distal enhancer | DEGs, DAGs | 1.330 | 1.07E-88 | FGFR3/GH04J001764 | eRNA_co-expression |
| *FGFR3* | chr4 | 1784490 | Astrocyte 1 | -8572 | 2.39 | 0.019 | Distal enhancer | DEGs, DAGs | 1.330 | 1.07E-88 | FGFR3/GH04J001775 | eQTLs |
| *FGFR3* | chr4 | 1784490 | Astrocyte 1 | -8572 | 2.39 | 0.019 | Distal enhancer | DEGs, DAGs | 1.330 | 1.07E-88 | FGFR3/GH04J001784 | eQTLs |
| *FGFR3* | chr4 | 1784490 | Astrocyte 1 | -8572 | 2.39 | 0.019 | Distal enhancer | DEGs, DAGs | 1.330 | 1.07E-88 | FGFR3/GH04J001792 | eQTLs |
| *AKAP10* | chr17 | 20072563 | Astrocyte 1 | -94976 | 4.56 | 0.019 | Distal enhancer | DEGs, DAGs | -1.245 | 1.19E-135 | AKAP10/GH17J019957 | eQTLs |
| *AKAP10* | chr17 | 20072563 | Astrocyte 1 | -94976 | 4.56 | 0.019 | Distal enhancer | DEGs, DAGs | -1.245 | 1.19E-135 | AKAP10/GH17J019976 | eQTLs |
| *AKAP10* | chr17 | 20072563 | Astrocyte 1 | -94976 | 4.56 | 0.019 | Distal enhancer | DEGs, DAGs | -1.245 | 1.19E-135 | AKAP10/GH17J020531 | eQTLs,TF_co-expression |
| *AKAP10* | chr17 | 20072563 | Astrocyte 1 | -94976 | 4.56 | 0.019 | Distal enhancer | DEGs, DAGs | -1.245 | 1.19E-135 | AKAP10/GH17J020225 | eQTLs,C-HiC |
| *AKAP10* | chr17 | 20072563 | Astrocyte 1 | -94976 | 4.56 | 0.019 | Distal enhancer | DEGs, DAGs | -1.245 | 1.19E-135 | AKAP10/GH17J020188 | eQTLs,C-HiC |
| *AKAP10* | chr17 | 20072563 | Astrocyte 1 | -94976 | 4.56 | 0.019 | Distal enhancer | DEGs, DAGs | -1.245 | 1.19E-135 | AKAP10/GH17J020089 | eQTLs,TF_co-expression |
| *AKAP10* | chr17 | 20072563 | Astrocyte 1 | -94976 | 4.56 | 0.019 | Distal enhancer | DEGs, DAGs | -1.245 | 1.19E-135 | AKAP10/GH17J020072 | eQTLs,C-HiC |
| *AKAP10* | chr17 | 20072563 | Astrocyte 1 | -94976 | 4.56 | 0.019 | Distal enhancer | DEGs, DAGs | -1.245 | 1.19E-135 | AKAP10/GH17J020008 | eQTLs,eRNA_co-expression,TF_co-expression |
| *ZFP36* | chr19 | 39397581 | Astrocyte 1 | -8982 | -3.35 | 0.024 | Distal enhancer | DEGs, DAGs | -1.425 | 9.33E-50 | ZFP36/GH19J039397 | eRNA_co-expression,TF_co-expression |
| *ZFP36* | chr19 | 39397581 | Astrocyte 1 | -8982 | -3.35 | 0.024 | Distal enhancer | DEGs, DAGs | -1.425 | 9.33E-50 | ZFP36/GH19J039552 | C-HiC,eRNA_co-expression |

| | | | | | | | | | | | | |
|---|---|---|---|---|---|---|---|---|---|---|---|---|
| *TCF20* | chr22 | 42283305 | Astrocyte 1 | -68113 | 6.17 | 0.034 | Distal enhancer | DEGs, DAGs | -1.233 | 3.20E-41 | TCF20/GH22J042176 | C-HiC,eRNA_co-expression |
| *TCF20* | chr22 | 42283305 | Astrocyte 1 | -68113 | 6.17 | 0.034 | Distal enhancer | DEGs, DAGs | -1.233 | 3.20E-41 | TCF20/GH22J042268 | C-HiC |
| *TCF20* | chr22 | 42283305 | Astrocyte 1 | -68113 | 6.17 | 0.034 | Distal enhancer | DEGs, DAGs | -1.233 | 3.20E-41 | TCF20/GH22J042271 | C-HiC |
| *TCF20* | chr22 | 42283305 | Astrocyte 1 | -68113 | 6.17 | 0.034 | Distal enhancer | DEGs, DAGs | -1.233 | 3.20E-41 | TCF20/GH22J042273 | C-HiC |
| *TCF20* | chr22 | 42283305 | Astrocyte 1 | -68113 | 6.17 | 0.034 | Distal enhancer | DEGs, DAGs | -1.233 | 3.20E-41 | TCF20/GH22J042282 | C-HiC |
| *TCF20* | chr22 | 42283305 | Astrocyte 1 | -68113 | 6.17 | 0.034 | Distal enhancer | DEGs, DAGs | -1.233 | 3.20E-41 | TCF20/GH22J042298 | TF_co-expression |
| *TCF20* | chr22 | 42283305 | Astrocyte 1 | -68113 | 6.17 | 0.034 | Distal enhancer | DEGs, DAGs | -1.233 | 3.20E-41 | TCF20/GH22J042320 | C-HiC |
| *TCF20* | chr22 | 42283305 | Astrocyte 1 | -68113 | 6.17 | 0.034 | Distal enhancer | DEGs, DAGs | -1.233 | 3.20E-41 | TCF20/GH22J042444 | C-HiC,eRNA_co-expression |
| *TCF20* | chr22 | 42283305 | Astrocyte 1 | -68113 | 6.17 | 0.034 | Distal enhancer | DEGs, DAGs | -1.233 | 3.20E-41 | TCF20/GH22J042436 | C-HiC,eRNA_co-expression |
| *TCF20* | chr22 | 42283305 | Astrocyte 1 | -68113 | 6.17 | 0.034 | Distal enhancer | DEGs, DAGs | -1.233 | 3.20E-41 | TCF20/GH22J042364 | C-HiC,eRNA_co-expression |
| *ANKDD1A* | chr15 | 64845826 | Astrocyte 1 | -65826 | 6.24 | 0.034 | Distal enhancer | DEGs, DAGs | 1.614 | 2.33E-151 | ANKDD1A/GH15J064840 | eQTLs,C-HiC |
| *ANKDD1A* | chr15 | 64845826 | Astrocyte 1 | -65826 | 6.24 | 0.034 | Distal enhancer | DEGs, DAGs | 1.614 | 2.33E-151 | ANKDD1A/GH15J064891 | eQTLs |
| *ANKDD1A* | chr15 | 64845826 | Astrocyte 1 | -65826 | 6.24 | 0.034 | Distal enhancer | DEGs, DAGs | 1.614 | 2.33E-151 | ANKDD1A/GH15J064903 | eQTLs |
| *JUNB* | chr19 | 12782903 | Astrocyte 1 | -8343 | 3.47 | 0.035 | Distal enhancer | DEGs, DAGs | -1.512 | 1.07E-62 | JUNB/GH19J012781 | eRNA_co-expression |
| *JUNB* | chr19 | 12782903 | Astrocyte 1 | -8343 | 3.47 | 0.035 | Distal enhancer | DEGs, DAGs | -1.512 | 1.07E-62 | JUNB/GH19J013160 | eQTLs,eRNA_co-expression |
| *JUNB* | chr19 | 12782903 | Astrocyte 1 | -8343 | 3.47 | 0.035 | Distal enhancer | DEGs, DAGs | -1.512 | 1.07E-62 | JUNB/GH19J013148 | eQTLs,eRNA_co-expression |
| *FGFR3* | chr4 | 1763318 | Astrocyte 1 | -29744 | 2.55 | 0.036 | Distal enhancer | DEGs, DAGs | 1.330 | 1.07E-88 | FGFR3/GH04J001757 | eQTLs,eRNA_co-expression |
| *FGFR3* | chr4 | 1763318 | Astrocyte 1 | -29744 | 2.55 | 0.036 | Distal enhancer | DEGs, DAGs | 1.330 | 1.07E-88 | FGFR3/GH04J001764 | eRNA_co-expression |
| *FGFR3* | chr4 | 1763318 | Astrocyte 1 | -29744 | 2.55 | 0.036 | Distal enhancer | DEGs, DAGs | 1.330 | 1.07E-88 | FGFR3/GH04J001775 | eQTLs |
| *FGFR3* | chr4 | 1763318 | Astrocyte 1 | -29744 | 2.55 | 0.036 | Distal enhancer | DEGs, DAGs | 1.330 | 1.07E-88 | FGFR3/GH04J001784 | eQTLs |
| *FGFR3* | chr4 | 1763318 | Astrocyte 1 | -29744 | 2.55 | 0.036 | Distal enhancer | DEGs, DAGs | 1.330 | 1.07E-88 | FGFR3/GH04J001792 | eQTLs |
| *NEK6* | chr9 | 124223400 | Astrocyte 1 | -34313 | 3.32 | 0.037 | Distal enhancer | DEGs, DAGs | -1.406 | 2.80E-71 | NEK6/GH09J124207 | eQTLs |
| *NEK6* | chr9 | 124223400 | Astrocyte 1 | -34313 | 3.32 | 0.037 | Distal enhancer | DEGs, DAGs | -1.406 | 2.80E-71 | NEK6/GH09J124210 | eQTLs |
| *NEK6* | chr9 | 124223400 | Astrocyte 1 | -34313 | 3.32 | 0.037 | Distal enhancer | DEGs, DAGs | -1.406 | 2.80E-71 | NEK6/GH09J124212 | eQTLs |
| *NEK6* | chr9 | 124223400 | Astrocyte 1 | -34313 | 3.32 | 0.037 | Distal enhancer | DEGs, DAGs | -1.406 | 2.80E-71 | NEK6/GH09J124213 | eQTLs |
| *NEK6* | chr9 | 124223400 | Astrocyte 1 | -34313 | 3.32 | 0.037 | Distal enhancer | DEGs, DAGs | -1.406 | 2.80E-71 | NEK6/GH09J124215 | eQTLs,eRNA_co-expression |
| *NEK6* | chr9 | 124223400 | Astrocyte 1 | -34313 | 3.32 | 0.037 | Distal enhancer | DEGs, DAGs | -1.406 | 2.80E-71 | NEK6/GH09J124220 | eQTLs,C-HiC |
| *NEK6* | chr9 | 124223400 | Astrocyte 1 | -34313 | 3.32 | 0.037 | Distal enhancer | DEGs, DAGs | -1.406 | 2.80E-71 | NEK6/GH09J124238 | eQTLs,eRNA_co-expression |
| *NEK6* | chr9 | 124223400 | Astrocyte 1 | -34313 | 3.32 | 0.037 | Distal enhancer | DEGs, DAGs | -1.406 | 2.80E-71 | NEK6/GH09J124242 | eQTLs |
| *NEK6* | chr9 | 124223400 | Astrocyte 1 | -34313 | 3.32 | 0.037 | Distal enhancer | DEGs, DAGs | -1.406 | 2.80E-71 | NEK6/GH09J124255 | eQTLs |
| *NEK6* | chr9 | 124223400 | Astrocyte 1 | -34313 | 3.32 | 0.037 | Distal enhancer | DEGs, DAGs | -1.406 | 2.80E-71 | NEK6/GH09J124256 | eQTLs |
| *NEK6* | chr9 | 124223400 | Astrocyte 1 | -34313 | 3.32 | 0.037 | Distal enhancer | DEGs, DAGs | -1.406 | 2.80E-71 | NEK6/GH09J124360 | eQTLs |

| | | | | | | | | | | | | |
|---|---|---|---|---|---|---|---|---|---|---|---|---|
| *NEK6* | chr9 | 124223400 | Astrocyte 1 | -34313 | 3.32 | 0.037 | Distal enhancer | DEGs, DAGs | -1.406 | 2.80E-71 | NEK6/GH09J124340 | eQTLs |
| *NEK6* | chr9 | 124223400 | Astrocyte 1 | -34313 | 3.32 | 0.037 | Distal enhancer | DEGs, DAGs | -1.406 | 2.80E-71 | NEK6/GH09J124308 | eQTLs |
| *NEK6* | chr9 | 124223400 | Astrocyte 1 | -34313 | 3.32 | 0.037 | Distal enhancer | DEGs, DAGs | -1.406 | 2.80E-71 | NEK6/GH09J124305 | eQTLs |
| *NEK6* | chr9 | 124223400 | Astrocyte 1 | -34313 | 3.32 | 0.037 | Distal enhancer | DEGs, DAGs | -1.406 | 2.80E-71 | NEK6/GH09J124303 | eQTLs |
| *NEK6* | chr9 | 124223400 | Astrocyte 1 | -34313 | 3.32 | 0.037 | Distal enhancer | DEGs, DAGs | -1.406 | 2.80E-71 | NEK6/GH09J124277 | eQTLs,eRNA_co-expression |
| *NEK6* | chr9 | 124223400 | Astrocyte 1 | -34313 | 3.32 | 0.037 | Distal enhancer | DEGs, DAGs | -1.406 | 2.80E-71 | NEK6/GH09J124270 | eQTLs |
| *STARD3* | chr17 | 39629899 | Astrocyte 1 | -6916 | 4.27 | 0.044 | Distal enhancer | DEGs, DAGs | -1.204 | 5.71E-37 | STARD3/GH17J039449 | eQTLs,TF_co-expression |
| *STARD3* | chr17 | 39629899 | Astrocyte 1 | -6916 | 4.27 | 0.044 | Distal enhancer | DEGs, DAGs | -1.204 | 5.71E-37 | STARD3/GH17J039460 | eQTLs,TF_co-expression |
| *STARD3* | chr17 | 39629899 | Astrocyte 1 | -6916 | 4.27 | 0.044 | Distal enhancer | DEGs, DAGs | -1.204 | 5.71E-37 | STARD3/GH17J039626 | eQTLs |
| *STARD3* | chr17 | 39629899 | Astrocyte 1 | -6916 | 4.27 | 0.044 | Distal enhancer | DEGs, DAGs | -1.204 | 5.71E-37 | STARD3/GH17J039631 | eQTLs |
| *STARD3* | chr17 | 39629899 | Astrocyte 1 | -6916 | 4.27 | 0.044 | Distal enhancer | DEGs, DAGs | -1.204 | 5.71E-37 | STARD3/GH17J039635 | eQTLs,Distance |
| *STARD3* | chr17 | 39629899 | Astrocyte 1 | -6916 | 4.27 | 0.044 | Distal enhancer | DEGs, DAGs | -1.204 | 5.71E-37 | STARD3/GH17J039658 | TF_co-expression |
| *STARD3* | chr17 | 39629899 | Astrocyte 1 | -6916 | 4.27 | 0.044 | Distal enhancer | DEGs, DAGs | -1.204 | 5.71E-37 | STARD3/GH17J039651 | eQTLs |
| *STARD3* | chr17 | 39629899 | Astrocyte 1 | -6916 | 4.27 | 0.044 | Distal enhancer | DEGs, DAGs | -1.204 | 5.71E-37 | STARD3/GH17J039649 | TF_co-expression |
| *STARD3* | chr17 | 39629899 | Astrocyte 1 | -6916 | 4.27 | 0.044 | Distal enhancer | DEGs, DAGs | -1.204 | 5.71E-37 | STARD3/GH17J039644 | eQTLs |
| *STARD3* | chr17 | 39629899 | Astrocyte 1 | -6916 | 4.27 | 0.044 | Distal enhancer | DEGs, DAGs | -1.204 | 5.71E-37 | STARD3/GH17J039641 | TF_co-expression |
| *STARD3* | chr17 | 39629899 | Astrocyte 1 | -6916 | 4.27 | 0.044 | Distal enhancer | DEGs, DAGs | -1.204 | 5.71E-37 | STARD3/GH17J039639 | eQTLs |
| *STARD3* | chr17 | 39631661 | Astrocyte 1 | -5154 | 3.23 | 0.046 | Distal enhancer | DEGs, DAGs | -1.204 | 5.71E-37 | STARD3/GH17J039449 | eQTLs,TF_co-expression |
| *STARD3* | chr17 | 39631661 | Astrocyte 1 | -5154 | 3.23 | 0.046 | Distal enhancer | DEGs, DAGs | -1.204 | 5.71E-37 | STARD3/GH17J039460 | eQTLs,TF_co-expression |
| *STARD3* | chr17 | 39631661 | Astrocyte 1 | -5154 | 3.23 | 0.046 | Distal enhancer | DEGs, DAGs | -1.204 | 5.71E-37 | STARD3/GH17J039626 | eQTLs |
| *STARD3* | chr17 | 39631661 | Astrocyte 1 | -5154 | 3.23 | 0.046 | Distal enhancer | DEGs, DAGs | -1.204 | 5.71E-37 | STARD3/GH17J039631 | eQTLs |
| *STARD3* | chr17 | 39631661 | Astrocyte 1 | -5154 | 3.23 | 0.046 | Distal enhancer | DEGs, DAGs | -1.204 | 5.71E-37 | STARD3/GH17J039635 | eQTLs |
| *STARD3* | chr17 | 39631661 | Astrocyte 1 | -5154 | 3.23 | 0.046 | Distal enhancer | DEGs, DAGs | -1.204 | 5.71E-37 | STARD3/GH17J039658 | TF_co-expression |
| *STARD3* | chr17 | 39631661 | Astrocyte 1 | -5154 | 3.23 | 0.046 | Distal enhancer | DEGs, DAGs | -1.204 | 5.71E-37 | STARD3/GH17J039651 | eQTLs |
| *STARD3* | chr17 | 39631661 | Astrocyte 1 | -5154 | 3.23 | 0.046 | Distal enhancer | DEGs, DAGs | -1.204 | 5.71E-37 | STARD3/GH17J039649 | TF_co-expression |
| *STARD3* | chr17 | 39631661 | Astrocyte 1 | -5154 | 3.23 | 0.046 | Distal enhancer | DEGs, DAGs | -1.204 | 5.71E-37 | STARD3/GH17J039644 | eQTLs |
| *STARD3* | chr17 | 39631661 | Astrocyte 1 | -5154 | 3.23 | 0.046 | Distal enhancer | DEGs, DAGs | -1.204 | 5.71E-37 | STARD3/GH17J039641 | TF_co-expression |
| *STARD3* | chr17 | 39631661 | Astrocyte 1 | -5154 | 3.23 | 0.046 | Distal enhancer | DEGs, DAGs | -1.204 | 5.71E-37 | STARD3/GH17J039639 | eQTLs |
| *MGRN1* | chr16 | 4614688 | Astrocyte 1 | -9852 | 2.79 | 0.007 | Distal enhancer | DEGs, DAGs | -1.201 | 4.68E-53 | MGRN1/GH16J004368 | eQTLs,eRNA_co-expression |
| *MGRN1* | chr16 | 4614688 | Astrocyte 1 | -9852 | 2.79 | 0.007 | Distal enhancer | DEGs, DAGs | -1.201 | 4.68E-53 | MGRN1/GH16J004611 | eQTLs |
| *MGRN1* | chr16 | 4614688 | Astrocyte 1 | -9852 | 2.79 | 0.007 | Distal enhancer | DEGs, DAGs | -1.201 | 4.68E-53 | MGRN1/GH16J004681 | eQTLs |
| *MGRN1* | chr16 | 4614688 | Astrocyte 1 | -9852 | 2.79 | 0.007 | Distal enhancer | DEGs, DAGs | -1.201 | 4.68E-53 | MGRN1/GH16J004675 | eQTLs |

| | | | | | | | | | | | | |
|---|---|---|---|---|---|---|---|---|---|---|---|---|
| *MGRN1* | chr16 | 4614688 | Astrocyte 1 | -9852 | 2.79 | 0.007 | Distal enhancer | DEGs, DAGs | -1.201 | 4.68E-53 | MGRN1/GH16J004672 | eQTLs |
| *MGRN1* | chr16 | 4614688 | Astrocyte 1 | -9852 | 2.79 | 0.007 | Distal enhancer | DEGs, DAGs | -1.201 | 4.68E-53 | MGRN1/GH16J004658 | eQTLs,eRNA_co-expression |
| *MGRN1* | chr16 | 4614688 | Astrocyte 1 | -9852 | 2.79 | 0.007 | Distal enhancer | DEGs, DAGs | -1.201 | 4.68E-53 | MGRN1/GH16J004651 | eQTLs |
| *MGRN1* | chr16 | 4614688 | Astrocyte 1 | -9852 | 2.79 | 0.007 | Distal enhancer | DEGs, DAGs | -1.201 | 4.68E-53 | MGRN1/GH16J004641 | eQTLs |
| *MGRN1* | chr16 | 4614688 | Astrocyte 1 | -9852 | 2.79 | 0.007 | Distal enhancer | DEGs, DAGs | -1.201 | 4.68E-53 | MGRN1/GH16J004639 | eQTLs |
| *MGRN1* | chr16 | 4614688 | Astrocyte 1 | -9852 | 2.79 | 0.007 | Distal enhancer | DEGs, DAGs | -1.201 | 4.68E-53 | MGRN1/GH16J004623 | eQTLs |
| *JUNB* | chr19 | 12796444 | Astrocyte 1 | 5198 | 6.13 | 0.011 | Distal enhancer | DEGs, DAGs | -1.512 | 1.07E-62 | JUNB/GH19J012781 | eRNA_co-expression |
| *JUNB* | chr19 | 12796444 | Astrocyte 1 | 5198 | 6.13 | 0.011 | Distal enhancer | DEGs, DAGs | -1.512 | 1.07E-62 | JUNB/GH19J013160 | eQTLs,eRNA_co-expression |
| *JUNB* | chr19 | 12796444 | Astrocyte 1 | 5198 | 6.13 | 0.011 | Distal enhancer | DEGs, DAGs | -1.512 | 1.07E-62 | JUNB/GH19J013148 | eQTLs,eRNA_co-expression |
| *SLC3A2* | chr11 | 62889982 | Astrocyte 1 | 34121 | 2.51 | 0.012 | Distal enhancer | DEGs, DAGs | 1.394 | 1.07E-83 | SLC3A2/GH11J062852 | eRNA_co-expression |
| *SLC3A2* | chr11 | 62889982 | Astrocyte 1 | 34121 | 2.51 | 0.012 | Distal enhancer | DEGs, DAGs | 1.394 | 1.07E-83 | SLC3A2/GH11J062887 | C-HiC |
| *SLC3A2* | chr11 | 62889982 | Astrocyte 1 | 34121 | 2.51 | 0.012 | Distal enhancer | DEGs, DAGs | 1.394 | 1.07E-83 | SLC3A2/GH11J062878 | C-HiC |
| *FOS* | chr14 | 75296006 | Astrocyte 1 | 17482 | 4.58 | 0.016 | Distal enhancer | DEGs, DAGs | -1.547 | 1.82E-60 | FOS/GH14J075273 | eRNA_co-expression |
| *FOS* | chr14 | 75296006 | Astrocyte 1 | 17482 | 4.58 | 0.016 | Distal enhancer | DEGs, DAGs | -1.547 | 1.82E-60 | FOS/GH14J075293 | eQTLs,eRNA_co-expression |
| *GADD45B* | chr19 | 2633229 | Astrocyte 1 | 157352 | 9.36 | 0.031 | Distal enhancer | DEGs, DAGs | -1.226 | 1.00E-23 | GADD45B/GH19J002474 | eRNA_co-expression |
| *GADD45B* | chr19 | 2633229 | Astrocyte 1 | 157352 | 9.36 | 0.031 | Distal enhancer | DEGs, DAGs | -1.226 | 1.00E-23 | GADD45B/GH19J002615 | eQTLs,eRNA_co-expression |
| *NXN* | chr17 | 896976 | Astrocyte 1 | 82544 | 7.18 | 0.034 | Distal enhancer | DEGs, DAGs | -1.299 | 3.52E-32 | NXN/GH17J000857 | eQTLs |
| *NXN* | chr17 | 896976 | Astrocyte 1 | 82544 | 7.18 | 0.034 | Distal enhancer | DEGs, DAGs | -1.299 | 3.52E-32 | NXN/GH17J000868 | eQTLs |
| *NXN* | chr17 | 896976 | Astrocyte 1 | 82544 | 7.18 | 0.034 | Distal enhancer | DEGs, DAGs | -1.299 | 3.52E-32 | NXN/GH17J000879 | eQTLs |
| *NXN* | chr17 | 896976 | Astrocyte 1 | 82544 | 7.18 | 0.034 | Distal enhancer | DEGs, DAGs | -1.299 | 3.52E-32 | NXN/GH17J000886 | eQTLs |
| *NXN* | chr17 | 896976 | Astrocyte 1 | 82544 | 7.18 | 0.034 | Distal enhancer | DEGs, DAGs | -1.299 | 3.52E-32 | NXN/GH17J000895 | eQTLs,C-HiC |
| *NXN* | chr17 | 896976 | Astrocyte 1 | 82544 | 7.18 | 0.034 | Distal enhancer | DEGs, DAGs | -1.299 | 3.52E-32 | NXN/GH17J000903 | eQTLs,C-HiC |
| *NXN* | chr17 | 896976 | Astrocyte 1 | 82544 | 7.18 | 0.034 | Distal enhancer | DEGs, DAGs | -1.299 | 3.52E-32 | NXN/GH17J000908 | eQTLs,C-HiC |
| *NXN* | chr17 | 896976 | Astrocyte 1 | 82544 | 7.18 | 0.034 | Distal enhancer | DEGs, DAGs | -1.299 | 3.52E-32 | NXN/GH17J000914 | eQTLs |
| *NXN* | chr17 | 896976 | Astrocyte 1 | 82544 | 7.18 | 0.034 | Distal enhancer | DEGs, DAGs | -1.299 | 3.52E-32 | NXN/GH17J000924 | eQTLs,C-HiC |
| *NXN* | chr17 | 896976 | Astrocyte 1 | 82544 | 7.18 | 0.034 | Distal enhancer | DEGs, DAGs | -1.299 | 3.52E-32 | NXN/GH17J000931 | eQTLs,C-HiC |
| *NXN* | chr17 | 896976 | Astrocyte 1 | 82544 | 7.18 | 0.034 | Distal enhancer | DEGs, DAGs | -1.299 | 3.52E-32 | NXN/GH17J000936 | eQTLs,C-HiC |
| *NXN* | chr17 | 896976 | Astrocyte 1 | 82544 | 7.18 | 0.034 | Distal enhancer | DEGs, DAGs | -1.299 | 3.52E-32 | NXN/GH17J000937 | eQTLs,C-HiC |
| *NXN* | chr17 | 896976 | Astrocyte 1 | 82544 | 7.18 | 0.034 | Distal enhancer | DEGs, DAGs | -1.299 | 3.52E-32 | NXN/GH17J000945 | C-HiC |
| *NXN* | chr17 | 896976 | Astrocyte 1 | 82544 | 7.18 | 0.034 | Distal enhancer | DEGs, DAGs | -1.299 | 3.52E-32 | NXN/GH17J000948 | eQTLs,C-HiC |
| *NXN* | chr17 | 896976 | Astrocyte 1 | 82544 | 7.18 | 0.034 | Distal enhancer | DEGs, DAGs | -1.299 | 3.52E-32 | NXN/GH17J000949 | eQTLs,C-HiC |
| *NXN* | chr17 | 896976 | Astrocyte 1 | 82544 | 7.18 | 0.034 | Distal enhancer | DEGs, DAGs | -1.299 | 3.52E-32 | NXN/GH17J000964 | eQTLs |

| | | | | | | | | | | | | |
|---|---|---|---|---|---|---|---|---|---|---|---|---|
| *HSPA1A* | chr6 | 31821171 | Astrocyte 1 | 5957 | 4.96 | 0.044 | Distal enhancer | DEGs, DAGs | 1.636 | 2.51E-30 | HSPA1A/GH06J031733 | C-HiC,eRNA_co-expression |
| *HSPA1A* | chr6 | 31821171 | Astrocyte 1 | 5957 | 4.96 | 0.044 | Distal enhancer | DEGs, DAGs | 1.636 | 2.51E-30 | HSPA1A/GH06J031813 | eRNA_co-expression |

FC: Fold change; DEGs: Differentially Expressed Genes; DAGs: Differentially Accessible Genes; TSS: Transcription Staring Site; Negative value in Distance to TSS reflects peaks falling upstream of TSS; Negative FC reflects higher accessibility or expression in ALA; NEG: not ELITE enhancer found

**Supplementary Table 7.** Subset of the differential functional analysis of the astrocyte cluster 1 DEG-DAG with positive correlation between expression and accessibility using EnrichR software

| **KEGG** Term | *p*-value | Adjusted *p*-value | Odds Ratio | Combined Score | Genes |
|---|---|---|---|---|---|
| Adrenergic signaling in cardiomyocytes | 3E-06 | 4E-04 | 2E+01 | 211.783 | *CAMK2B;CACNA2D3;ATP1A2;ATP1B2;ADRA1A; RAPGEF3* |
| Cardiac muscle contraction | 1E-04 | 5E-03 | 2E+01 | 173.599 | *ASPH;CACNA2D3;ATP1A2;ATP1B2* |
| Protein digestion and absorption | 2E-04 | 7E-03 | 2E+01 | 135.147 | *COL5A3;ATP1A2;SLC3A2;ATP1B2* |
| Calcium signaling pathway | 5E-04 | 1E-02 | 8E+00 | 63.597 | *CAMK2B;ASPH;ADRA1A;FGFR3;VEGFA* |
| Mineral absorption | 6E-04 | 1E-02 | 2E+01 | 149.244 | *ATP1A2;ATP1B2;FTL* |
| Gastric acid secretion | 1E-03 | 2E-02 | 2E+01 | 105.620 | *CAMK2B;ATP1A2;ATP1B2* |
| MAPK signaling pathway | 1E-03 | 2E-02 | 7E+00 | 45.380 | *CACNA2D3;HSPB1;FGFR3;VEGFA;HSPA1A* |
| Insulin secretion | 2E-03 | 2E-02 | 1E+01 | 87.931 | *CAMK2B;ATP1A2;ATP1B2* |
| Proximal tubule bicarbonate reclamation | 2E-03 | 2E-02 | 4E+01 | 226.067 | *ATP1A2;ATP1B2* |
| Salivary secretion | 2E-03 | 2E-02 | 1E+01 | 78.214 | *ATP1A2;ATP1B2;ADRA1A* |
| Aldosterone synthesis and secretion | 2E-03 | 2E-02 | 1E+01 | 72.281 | *CAMK2B;ATP1A2;ATP1B2* |
| Rap1 signaling pathway | 3E-03 | 2E-02 | 8E+00 | 44.588 | *RAPGEF5;FGFR3;RAPGEF3;VEGFA* |
| cAMP signaling pathway | 3E-03 | 3E-02 | 7E+00 | 42.573 | *CAMK2B;ATP1A2;ATP1B2;RAPGEF3* |
| Aldosterone-regulated sodium reabsorption | 5E-03 | 4E-02 | 2E+01 | 115.296 | *ATP1A2;ATP1B2* |
| Bladder cancer | 6E-03 | 4E-02 | 2E+01 | 99.579 | *FGFR3;VEGFA* |
| Ferroptosis | 6E-03 | 4E-02 | 2E+01 | 99.579 | *SLC3A2;FTL* |
| Carbohydrate digestion and absorption | 7E-03 | 5E-02 | 2E+01 | 81.840 | *ATP1A2;ATP1B2* |
| Cell adhesion molecules | 8E-03 | 5E-02 | 8E+00 | 38.172 | *IGSF11;SDC4;NRXN1* |
| Endocrine and other factor-regulated calcium reabsorption | 9E-03 | 5E-02 | 1E+01 | 68.772 | *ATP1A2;ATP1B2* |

| **GO-BP** Term | *p*-value | Adjusted *p*-value | Odds Ratio | Combined Score | Genes |
|---|---|---|---|---|---|
| regulation of protein kinase C signaling (GO:0090036) | 7E-06 | 6E-03 | 1E+02 | 1240.796 | *DGKG;ADRA1A;VEGFA* |
| positive regulation of cellular component organization (GO:0051130) | 2E-05 | 7E-03 | 2E+01 | 201.815 | *CAMK2B;NRXN1;APOE;RAPGEF3;VEGFA* |
| positive regulation of endothelial cell chemotaxis by VEGF-activated vascular endothelial growth factor receptor signaling pathway (GO:0038033) | 7E-05 | 2E-02 | 3E+02 | 2386.360 | *HSPB1;VEGFA* |

| | | | | | |
|---|---|---|---|---|---|
| positive regulation of developmental process (GO:0051094) | 1E-04 | 3E-02 | 1E+01 | 103.340 | *CAMK2B;NRXN1;APOE;RAPGEF3;VEGFA* |
| positive regulation of cell migration by vascular endothelial growth factor signaling pathway (GO:0038089) | 2E-04 | 3E-02 | 1E+02 | 1064.547 | *HSPB1;VEGFA* |
| postsynaptic density organization (GO:0097106) | 2E-04 | 3E-02 | 1E+02 | 1064.547 | *NRXN1;TMEM108* |
| positive regulation of protein kinase C signaling (GO:0090037) | 3E-04 | 3E-02 | 1E+02 | 885.601 | *ADRA1A;VEGFA* |
| positive regulation of synapse maturation (GO:0090129) | 3E-04 | 3E-02 | 1E+02 | 885.601 | *CAMK2B;NRXN1* |
| regulation of axon guidance (GO:1902667) | 3E-04 | 3E-02 | 9E+01 | 754.043 | *ZSWIM6;VEGFA* |
| regulation of synapse maturation (GO:0090128) | 4E-04 | 4E-02 | 8E+01 | 653.601 | *CAMK2B;NRXN1* |
| regulation of blood vessel endothelial cell migration (GO:0043535) | 5E-04 | 4E-02 | 2E+01 | 169.285 | *HSPB1;APOE;VEGFA* |
| regulation of cardiac conduction (GO:1903779) | 5E-04 | 4E-02 | 2E+01 | 169.285 | *ASPH;ATP1A2;ATP1B2* |
| cellular potassium ion homeostasis (GO:0030007) | 6E-04 | 4E-02 | 7E+01 | 511.060 | *ATP1A2;ATP1B2* |
| sodium ion export across plasma membrane (GO:0036376) | 6E-04 | 4E-02 | 7E+01 | 511.060 | *ATP1A2;ATP1B2* |
| diterpenoid metabolic process (GO:0016101) | 7E-04 | 4E-02 | 2E+01 | 135.874 | *SDC4;GPC5;APOE* |
| cellular sodium ion homeostasis (GO:0006883) | 8E-04 | 4E-02 | 6E+01 | 415.398 | *ATP1A2;ATP1B2* |
| positive regulation of endothelial cell chemotaxis (GO:2001028) | 8E-04 | 4E-02 | 6E+01 | 415.398 | *HSPB1;VEGFA* |
| retrograde axonal transport (GO:0008090) | 9E-04 | 4E-02 | 5E+01 | 378.628 | *DST;TMEM108* |
| membrane repolarization (GO:0086009) | 9E-04 | 4E-02 | 5E+01 | 378.628 | *ATP1A2;ATP1B2* |
| regulation of cholesterol biosynthetic process (GO:0045540) | 1E-03 | 4E-02 | 5E+01 | 347.182 | *SREBF1;APOE* |
| cell communication by electrical coupling involved in cardiac conduction (GO:0086064) | 1E-03 | 4E-02 | 5E+01 | 347.182 | *ATP1A2;ATP1B2* |
| neurotransmitter transport (GO:0006836) | 1E-03 | 4E-02 | 2E+01 | 112.074 | *NRXN1;SLC1A2;ATP1A2* |
| postsynaptic membrane organization (GO:0001941) | 1E-03 | 5E-02 | 4E+01 | 296.332 | *NRXN1;APOE* |
| cellular monovalent inorganic cation homeostasis (GO:0030004) | 1E-03 | 5E-02 | 4E+01 | 275.527 | *ATP1A2;ATP1B2* |
| positive regulation of heart contraction (GO:0045823) | 1E-03 | 5E-02 | 4E+01 | 275.527 | *ATP1A2;ADRA1A* |
| negative regulation of cellular component organization (GO:0051129) | 1E-03 | 5E-02 | 1E+01 | 97.907 | *RAPGEF3;VEGFA;HSPA1A* |
| regulation of cholesterol metabolic process (GO:0090181) | 2E-03 | 5E-02 | 4E+01 | 257.121 | *SREBF1;APOE* |
| regulation of dendritic spine development (GO:0060998) | 2E-03 | 5E-02 | 4E+01 | 257.121 | *CAMK2B;APOE* |

| | | | | | |
|---|---|---|---|---|---|
| vascular transport (GO:0010232) | 2E-03 | 5E-02 | 1E+01 | 91.067 | *SLC1A2;ATP1A2;ATP1B2* |
| positive regulation of axon extension (GO:0045773) | 2E-03 | 5E-02 | 4E+01 | 240.736 | *MACF1;VEGFA* |
| positive regulation of dendritic spine development (GO:0060999) | 2E-03 | 5E-02 | 4E+01 | 240.736 | *CAMK2B;APOE* |
| transport across blood-brain barrier (GO:0150104) | 2E-03 | 5E-02 | 1E+01 | 87.931 | *SLC1A2;ATP1A2;ATP1B2* |
| regulation of transcription from RNA polymerase II promoter in response to stress (GO:0043618) | 2E-03 | 5E-02 | 1E+01 | 86.427 | *HIF3A;VEGFA;HSPA1A* |
| sodium ion transmembrane transport (GO:0035725) | 2E-03 | 5E-02 | 1E+01 | 86.427 | *SLC6A11;ATP1A2;ATP1B2* |
| positive regulation of neuron projection development (GO:0010976) | 2E-03 | 5E-02 | 1E+01 | 84.965 | *CAMK2B;ATP1B2;APOE* |
| amino acid import across plasma membrane (GO:0089718) | 2E-03 | 5E-02 | 3E+01 | 212.867 | *SLC1A2;SLC3A2* |
| potassium ion homeostasis (GO:0055075) | 2E-03 | 5E-02 | 3E+01 | 212.867 | *ATP1A2;ATP1B2* |
| retinoid metabolic process (GO:0001523) | 2E-03 | 5E-02 | 1E+01 | 79.493 | *SDC4;GPC5;APOE* |
| sodium ion homeostasis (GO:0055078) | 2E-03 | 5E-02 | 3E+01 | 200.934 | *ATP1A2;ATP1B2* |
| regulation of heart contraction (GO:0008016) | 2E-03 | 5E-02 | 1E+01 | 78.214 | *ASPH;ATP1A2;ATP1B2* |
| L-alpha-amino acid transmembrane transport (GO:1902475) | 2E-03 | 5E-02 | 3E+01 | 180.226 | *SLC1A2;SLC3A2* |

| **GO-CC** Term | *p*-value | Adjusted *p*-value | Odds Ratio | Combined Score | Genes |
|---|---|---|---|---|---|
| astrocyte projection (GO:0097449) | 2E-04 | 2E-02 | 1E+02 | 1064.547 | *SLC1A2;ATP1B2* |
| sodium:potassium-exchanging ATPase complex (GO:0005890) | 3E-04 | 2E-02 | 9E+01 | 754.043 | *ATP1A2;ATP1B2* |
| glial cell projection (GO:0097386) | 7E-04 | 2E-02 | 6E+01 | 458.896 | *SLC1A2;ATP1B2* |
| cation-transporting ATPase complex (GO:0090533) | 9E-04 | 2E-02 | 5E+01 | 378.628 | *ATP1A2;ATP1B2* |

| **RNA-Seq Disease Gene and Drug Signatures from GEO** Term | *p*-value | Adjusted *p*-value | Odds Ratio | Combined Score | Genes |
|---|---|---|---|---|---|
| Huntington's (Grade 3) BA4 (Motor Cortex) Grade 3 GSE79666 up | 2E-21 | 2E-18 | 3E+01 | 1307.651 | *MACF1;SDC4;SLC1A2;HSPB1;ATP1A2;HIF3A;MTRNR2L8;ATP1B2;VEGFA;ADGRG1;COL5A3;SAFB2;CPE;FAM171B;CKB;APOE;RAPGEF5;FGFR3;RAPGEF3;FTL;B4GALNT4;HSPA1A* |
| Prmt5 Neural Stem/Progenitors Cells knockout GSE45284 up | 1E-14 | 7E-12 | 2E+01 | 582.369 | *CAMK2B;MACF1;SDC4;NRXN1;NTM;SLC1A2;SLC6A11;MMD2;ATP1A2;SLC3A2;ATP1B2;KLF6;SAFB2;CPE;FAM171B;CKB;APOE* |

| | | | | | |
|---|---|---|---|---|---|
| Lin28 Neural stem cells Overexpression GSE77851 down | 3E-12 | 8E-10 | 2E+01 | 404.225 | *CAMK2B;SREBF1;MACF1;SLC1A2;SLC6A11;MMD2;ATP1A2;SLC3A2;ATP1B2;VEGFA;CPE;FAM171B;CKB;APOE;FGFR3* |
| LPS Astrocyte GSE75246 down | 3E-12 | 8E-10 | 2E+01 | 400.339 | *NRXN1;NTM;SLC1A2;SLC6A11;MMD2;ATP1A2;ATP1B2;VEGFA;IGSF11;ADGRG1;GPC5;FAM171B;CKB;APOE;FGFR3* |
| Topotecan Cortical neurons 300 nM GSE43526 down | 3E-12 | 8E-10 | 2E+01 | 400.339 | *CAMK2B;MACF1;DST;SDC4;NRXN1;NTM;CACNA2D3;SLC1A2;MMD2;ATP1A2;SLC3A2;ATP1B2;CPE;CKB;APOE* |
| ALS (Familial) Whole lumbar Spinal Cord GSE52946 down | 6E-10 | 1E-07 | 1E+01 | 265.686 | *MACF1;DST;SDC4;SLC1A2;SLC6A11;ATP1A2;MTRNR2L8;ATP1B2;PURA;CKB;RAPGEF5;FGFR3;FTL* |
| OPC Cerebral Cortex Knockdown GSE74647 up | 8E-09 | 1E-06 | 1E+01 | 210.046 | *MACF1;KLF6;SDC4;NRXN1;CACNA2D3;SLC1A2;SLC6A11;ATP1A2;SLC3A2;FAM171B;ATP1B2;CKB* |
| TAF15 Striatum Knockdown GSE77703 down | 8E-09 | 1E-06 | 1E+01 | 210.046 | *CAMK2B;MACF1;DST;NRXN1;NTM;CACNA2D3;SLC1A2;CPE;SLC6A11;CKB;APOE;CDK14* |
| Chd7 Spinal Cord Conditional Knockout GSE72726 up | 8E-09 | 1E-06 | 1E+01 | 208.655 | *ADGRG1;SDC4;NRXN1;NTM;CPE;SLC6A11;MMD2;ATP1A2;FAM171B;ATP1B2;APOE;FGFR3* |
| Acromegaly Adipose GSE57803 down | 8E-09 | 1E-06 | 1E+01 | 207.965 | *ZFP36;KLF6;ASPH;COL5A3;CPE;SLC3A2;HIF3A;MTRNR2L8;RAPGEF3;FTL;AKAP1;HSPA1A* |
| Nfix Brain Knockout GSE65337 up | 9E-08 | 8E-06 | 1E+01 | 163.596 | *CAMK2B;SREBF1;MACF1;ADGRG1;NRXN1;CPE;SLC6A11;ATP1A2;SLC3A2;ATP1B2;CKB* |
| Olig2 Brain tumor conditional Knockout GSE71493 up | 9E-08 | 8E-06 | 1E+01 | 163.596 | *IGSF11;MACF1;DST;NRXN1;SLC1A2;SLC6A11;MMD2;ATP1A2;SLC3A2;APOE;VEGFA* |
| MeCP2 Hypothalamus Transgenic GSE66871 up | 9E-08 | 8E-06 | 1E+01 | 161.954 | *MACF1;ADGRG1;SLC1A2;CPE;TCF20;SLC6A11;MMD2;ATP1A2;ATP1B2;APOE;FGFR3* |

| **HDSigDB human 2021** Term | *p*-value | Adjusted *p*-value | Odds Ratio | Combined Score | Genes |
|---|---|---|---|---|---|
| Genes Changed In Astrocytes Of HD Patients Vs Control PMID32681824 | 2E-22 | 3E-19 | 2E+01 | 811.321 | *DGKG;IQCA1;SDC4;NRXN1;NTM;SLC1A2;HSPB1;TCF20;ATP1A2;SLC3A2;HIF3A;LRMDA;ADGRG1;GPC5;ZSWIM6;CKB;APOE;B4GALNT4;ANKDD1A;CACNA2D3;ZNF98;SLC6A11;MMD2;ATP1B2;FRMD4B;VEGFA;IGSF11;SDK1;KLF6;ASPH;KANSL1;COL5A3;CPE;FAM171B;RAPGEF5;FGFR3;FTL* |
| Genes Changed In Human HD Astrocytes Vs Control Astrocytes PMID32070434 | 3E-20 | 2E-17 | 1E+01 | 634.466 | *MACF1;IQCA1;NRXN1;NTM;SLC1A2;HSPB1;ATP1A2;MTRNR2L8;AKAP1;LRMDA;LCNL1;ZFP36;ADGRG1;TMEM108;CKB;APOE;B4GALNT4;ANKDD1A;SREBF1;DST;ZNF98;SLC6A11;MMD2;V* |

| | | | | | |
|---|---|---|---|---|---|
| | | | | | *EGFA;KLF6;KANSL1;COL5A3;CPE;FAM171B;RAPGEF5;FGFR3;RAPGEF3;FTL;HSPA1A* |
| Top Astrocyte-Enriched Genes In Humans And Mice PMID29892006 | 2E-18 | 9E-16 | 2E+01 | 664.072 | *MACF1;IQCA1;SDC4;NRXN1;NTM;SLC1A2;ATP1A2;SLC3A2;HIF3A;ADRA1A;GPC5;CKB;APOE;SREBF1;SLC6A11;MMD2;ATP1B2;VEGFA;IGSF11;ASPH;COL5A3;CPE;FAM171B;FGFR3;RAPGEF3* |
| Genes Down-Regulated In Astrocytes Of HD Patients Vs Control PMID32681824 | 3E-18 | 1E-15 | 1E+01 | 544.027 | *DGKG;IQCA1;SDC4;NRXN1;NTM;SLC1A2;ATP1A2;SLC3A2;HIF3A;LRMDA;ADGRG1;GPC5;ZSWIM6;CKB;APOE;B4GALNT4;ANKDD1A;ZNF98;SLC6A11;MMD2;ATP1B2;VEGFA;IGSF11;ASPH;COL5A3;CPE;FAM171B;FGFR3;FTL* |
| Genes Changed In Astrocytes Of R6/2 Vs WT PMID32681824 | 2E-16 | 6E-14 | 4E+01 | 1597.803 | *MACF1;NRXN1;NTM;SLC1A2;MMD2;ATP1A2;ATP1B2;VEGFA;LRMDA;CPE;GPC5;CKB;APOE* |
| Top Astrocyte-Specific Genes In Humans And Mice PMID29892006 | 4E-16 | 9E-14 | 1E+01 | 495.611 | *SREBF1;MACF1;IQCA1;SDC4;NRXN1;NTM;SLC1A2;SLC6A11;MMD2;ATP1A2;HIF3A;ATP1B2;VEGFA;IGSF11;ASPH;COL5A3;CPE;GPC5;FAM171B;CKB;APOE;FGFR3;RAPGEF3* |
| Genes Down-Regulated In Astrocytes Of R6/2 Vs WT PMID32681824 | 4E-16 | 9E-14 | 5E+01 | 1835.168 | *MACF1;NRXN1;NTM;SLC1A2;CPE;MMD2;ATP1A2;GPC5;ATP1B2;CKB;APOE;VEGFA* |
| Genes Changed In OPCs Of HD Patients Vs Control PMID32681824 | 5E-16 | 9E-14 | 2E+01 | 553.114 | *DGKG;MACF1;DST;NRXN1;CACNA2D3;SLC1A2;HSPB1;MMD2;HIF3A;VEGFA;LRMDA;ADGRG1;KANSL1;TMEM108;CPE;GPC5;CKB;APOE;RAPGEF5;FTL;HSPA1A* |
| Top Astrocyte-Expressed Genes In Humans And Mice PMID29892006 | 8E-13 | 1E-10 | 1E+01 | 308.676 | *MACF1;SDC4;NRXN1;NTM;SLC1A2;SLC6A11;MMD2;ATP1A2;SLC3A2;ATP1B2;IGSF11;PURA;ZFP36;ASPH;CPE;GPC5;FAM171B;CKB;APOE;FGFR3* |

| **ENCODE Histone Modifications 2015 Term** | *p*-value | Adjusted *p*-value | Odds Ratio | Combined Score | Genes |
|---|---|---|---|---|---|
| H3K27me3 astrocyte hg19 | 8E-04 | 3E-02 | 3E+00 | 22.001 | *ANKDD1A;MACF1;IQCA1;NRXN1;KLHL32;MMD2;HIF3A;ADRA1A;CPE;GPC5;CKB;APOE;RAPGEF5;FGFR3* |

Colors indicate pathways relevant to Alzheimer Disease

**Supplementary Figure 1.** SnATAC-seq data analysis for sample bias.

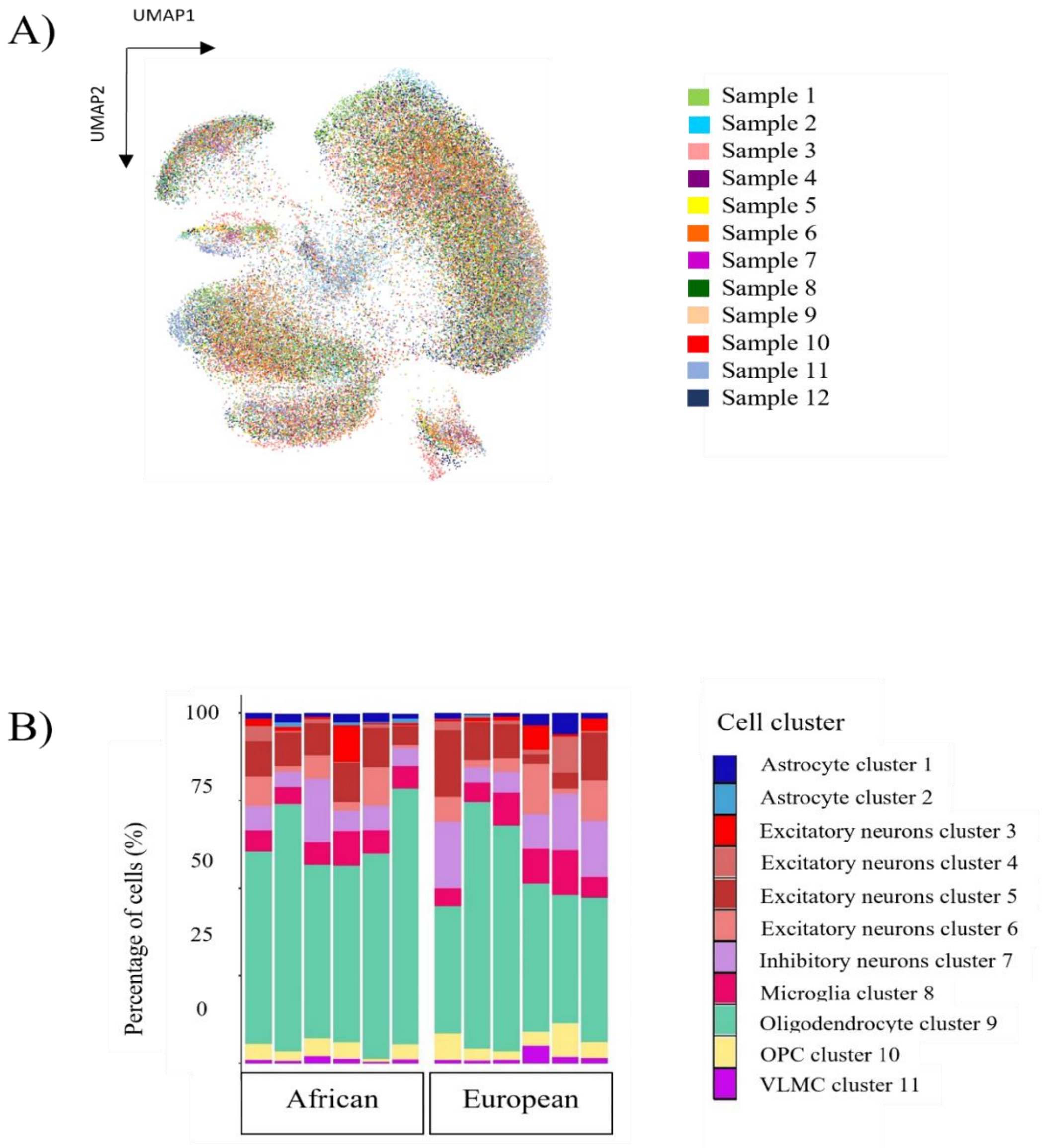


A) UMAP representation of integrated snATAC-seq and snRNA-seq of individual samples across all clusters, each color represents a sample; B) Ancestry distribution of percentage of cells per cluster from snATAC-seq data.

**Supplementary Figure 2.** SnATAC-seq cell marker cluster identification by chromatin accessibility.

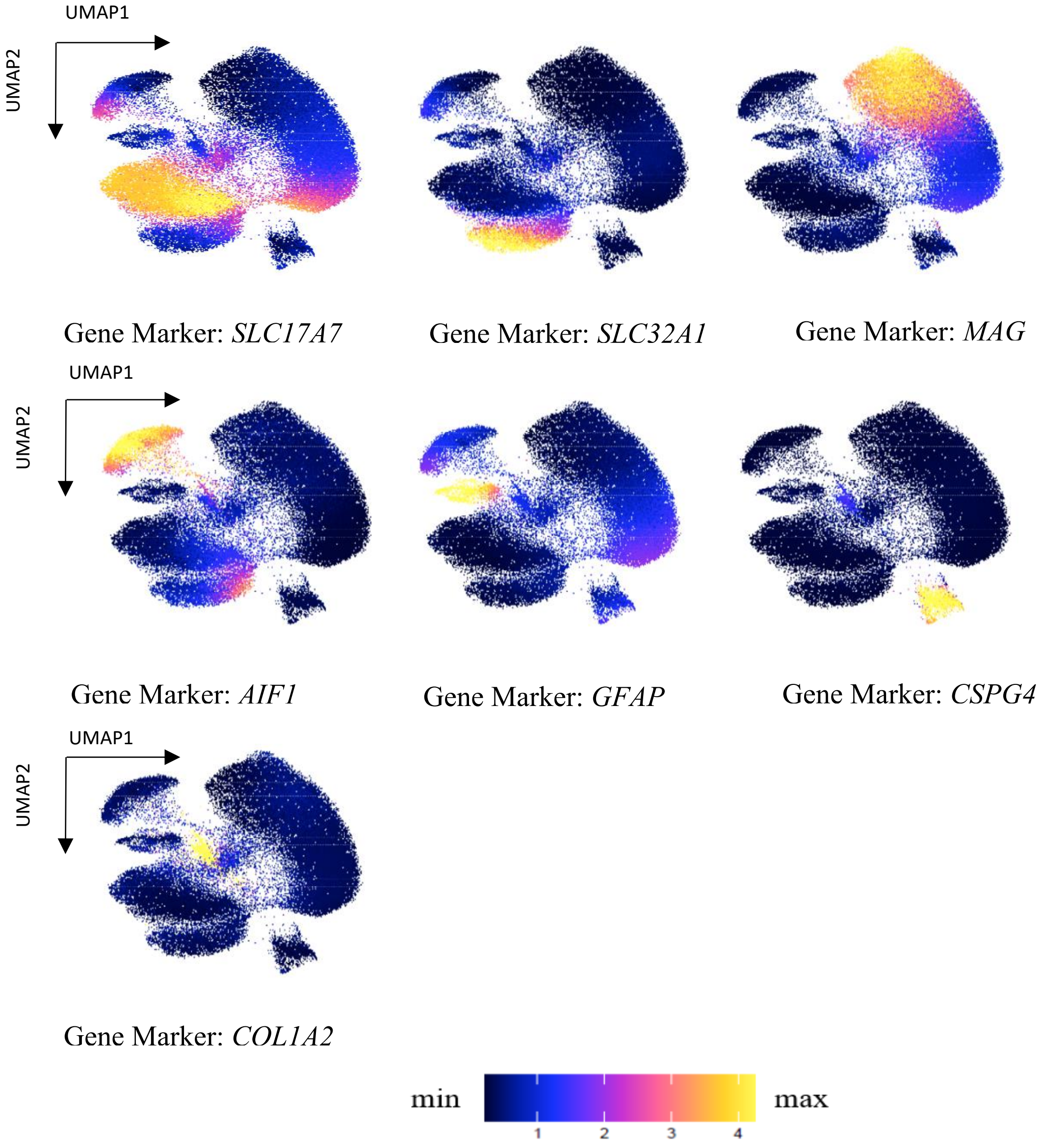


Marker genes used for cell type identification: *SLC1717* for excitatory neurons; *SLC32A1* for inhibitory neurons; *MAG* for oligodendrocytes; *AIF1* for microglia; *GFAP* for astrocytes, *CSPG4* for OPCs and *COL1A2* for VLMC.

**Supplementary Figure 3.** Astrocyte subtype classification.

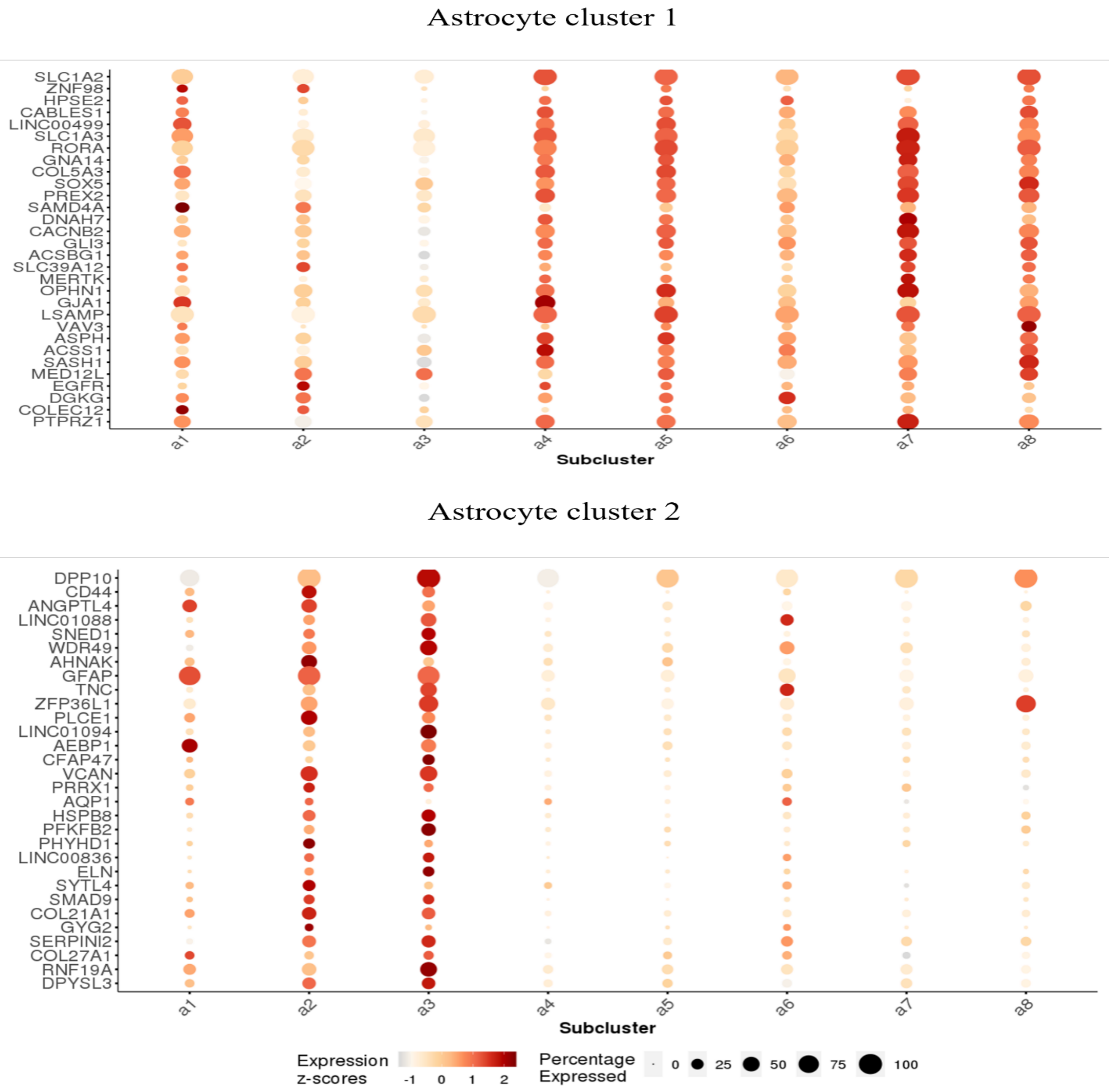


Bubble plot showing the gene markers for astrocyte cluster 1 and astrocyte cluster 2 and their expression in the single cell atlas of the Entorhinal Cortex in Human Alzheimer's Disease (http://adsn.ddnetbio.com). The color represents the relative gene expression, and the bubble size is the proportion of cells in the group expressing the gene.

**Supplementary Figure 4.** *APOE* chromatin accessibility in the other cell clusters where *APOE* is highly expressed but not differentially accessible between the ancestries.

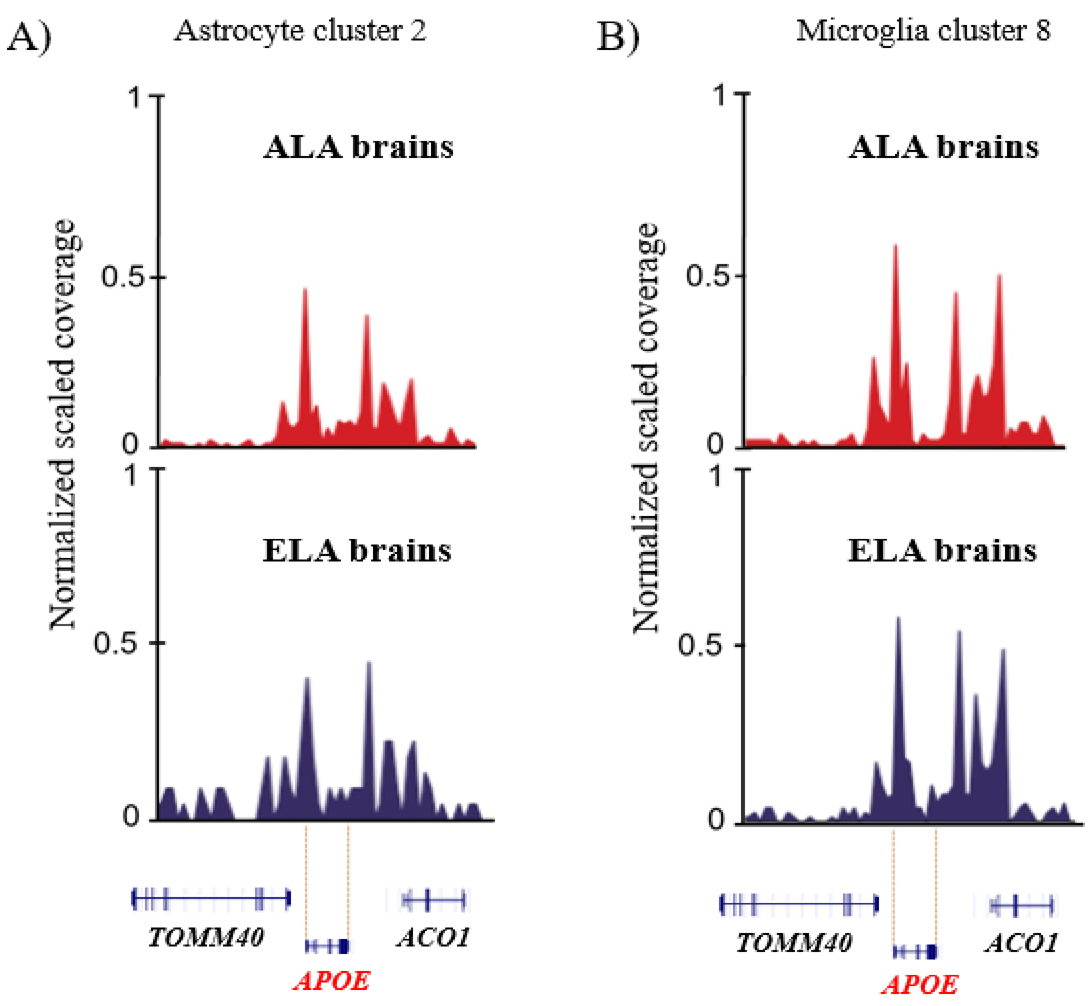


Visualization of chromatin accessible peaks in the *APOE* local ancestry region between ancestries from Astrocyte cluster 2 (A) and Microglia cluster 8 (B). No significant differentially accessible peaks were observed between ancestries in either cluster.

**Supplementary Figure 5.** Visualization of local ancestry across all chromosomes per individual sample.

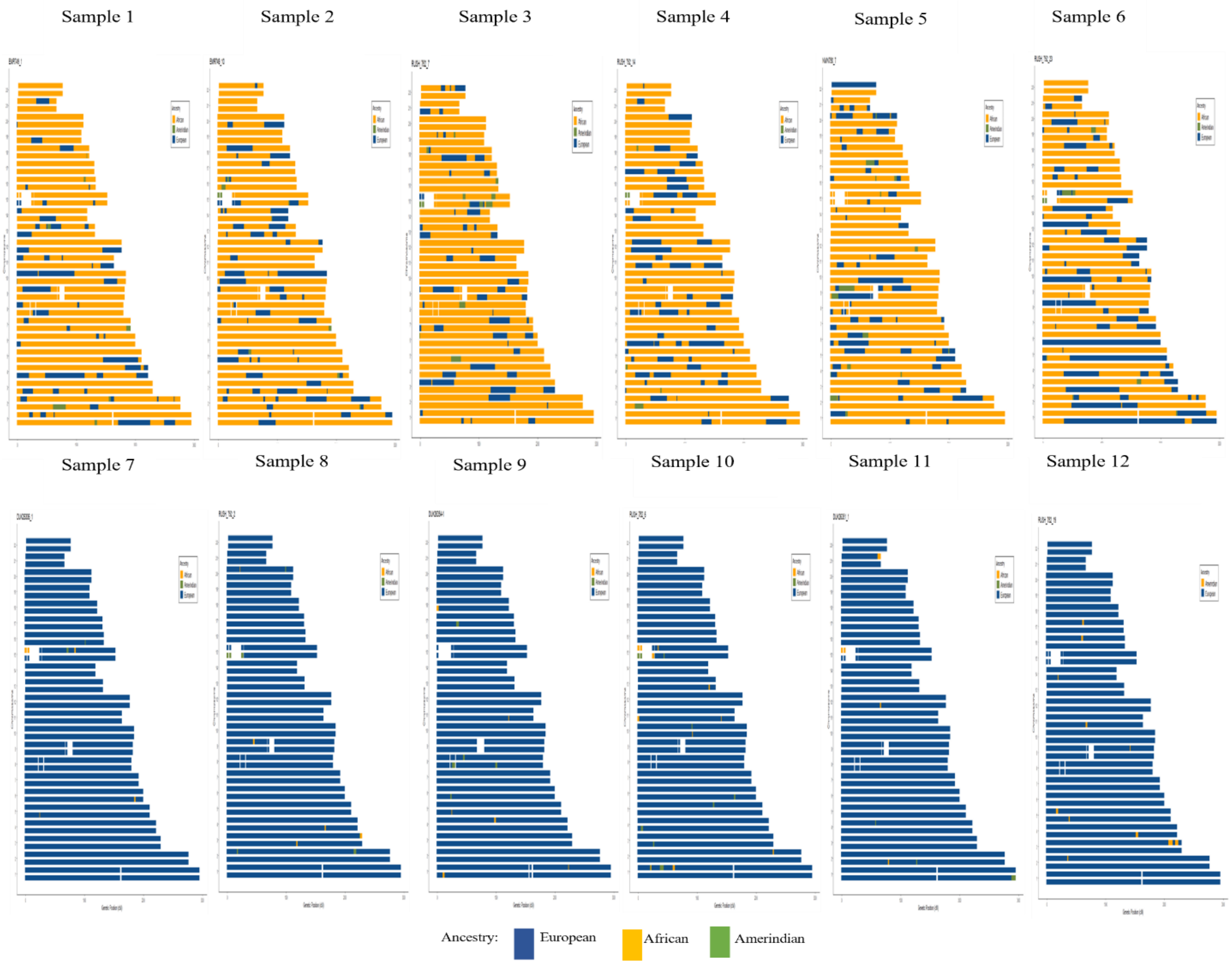


Bar-plot showing the local ancestry across the genome. The y-axis represents the chromosomes and the x-axis represent the chromosomal position. Ancestries are represented by colors Blue (European), Yellow (African) and Green (Amerindian).

**Supplementary Figure 6.** Enrichment of differentially accessible peak between ancestries in chromosome 19.

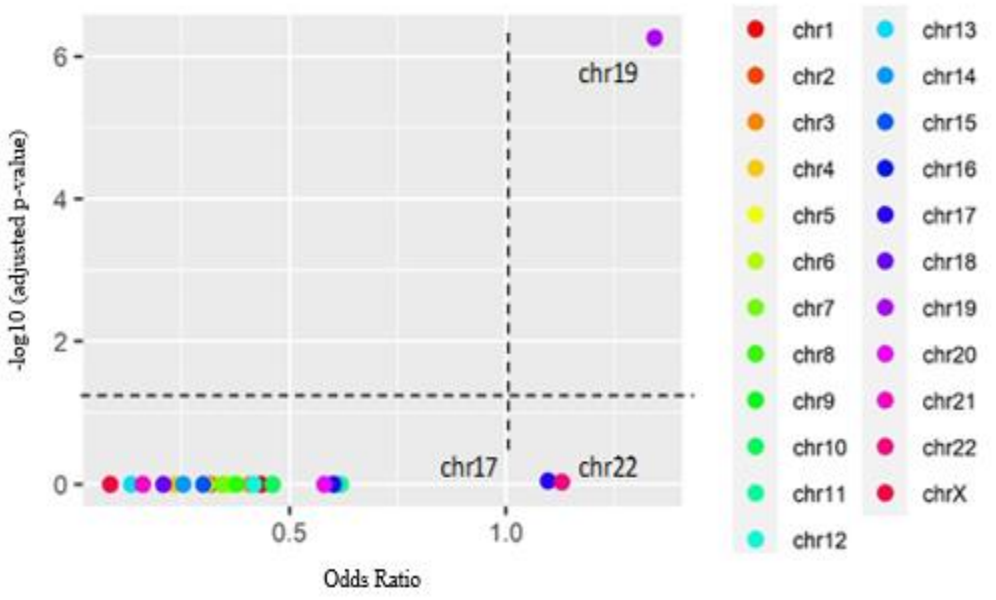


The plot shows the significance of each gene set (genes associated to differentially accessible peaks by GREAT and HOMER software, located in a given chromosome) versus its odds ratio. Each point represents a single gene set (chromosome); the x-axis measures the odds ratio (0, inf) calculated for the gene set, while the y-axis gives the -log10 (adjusted *p*-value) of the gene set. The colored points represent each chromosome. The dotted lines represent statistical significance thresholds, the horizontal line represents the -log10 of an adjusted *p*-value <0.05. The -log10 values >1.3 show statistical significance. The vertical line represents that Odds ratios higher than 1 can be considered significant.
